# Systematic mutagenesis assay promotes comprehension of the strand-bias laws for mutations induced by oxidative DNA damage

**DOI:** 10.1101/2024.02.29.581290

**Authors:** Hidehiko Kawai, Shungo Ebi, Ryusei Sugihara, Chiho Fujiwara, Yoshihiro Fujikawa, Shingo Kimura, Hiroyuki Kamiya

**Author notes:** To whom correspondence should be addressed.; (H. Kawai) or (H. Kamiya).

## Abstract

We have recently developed an efficient and sensitive method for analyzing mutations caused by various environmental and endogenous factors which utilizes nucleotide-barcoded *supF* shuttle vector libraries with a multiplexed NGS assay, referred to hereafter as *supF* NGS assay. Ionizing-radiation-induced cancer is known to be difficult to distinguish from spontaneous cancer, especially in the case of low-dose and low-dose-rate exposure, and discerning the underlying mechanisms of ionizing-radiation-induced cancer, especially the relationship between mutagenesis and carcinogenesis, is likely to be an arduous task. In the present study, we have attempted to address the mutations characteristic for exposure to low levels of ionizing radiation by using the *supF* NGS assay. As a result, a significant increase in mutations was detected at cytosines and guanines within 5’-TC-3’:5’-GA-3’ sites following chronic gamma-irradiation at a dose-rate of 1 Gy per day for the duration of 2 days. Since the number of detected mutations exceeded the expectations based on the quantity of DNA-damage induced by irradiation, we proceeded to explore the possibilities that a single DNA-lesion induced by irradiation may cause amplification of mutations. For this purpose, we utilized shuttle vector libraries with a single 8-oxo-7,8-dihydroguanine (8-oxo-G)-damaged residue introduced at different sites via an *in vitro* enzymatic method. Through a set of experiments, we revealed that a single 8-oxo-G-damaged residue can become a trigger for peripheral mutagenesis; intense generation of strand-biased mutations occured at 5’-TC-3’:5’-GA-3’ sites with specific localization in the secondary structures of single-stranded DNA, more frequently than not at sites different from the 8-oxo-G-damaged sites. Thus, this study provides a novel prospect for the role of DNA-lesions induced by environmentally or endogenously generated ROS in additional mutations. The high-performance mutagenesis assay presented in this study will advance research aimed at uncovering the mechanisms of mutagenesis and the intricacies relevant to carcinogenesis.

## Introduction

The progressive co-development of NGS technologies and cancer genomics databases have enabled the identification of distinct mutational signatures in various cancers, and so far about a half of the identified mutational signatures have underlying processes been assigned to them (1–4). Mutational signatures originate from the effects and interactions of a vast range of environmental and intrinsic sources, such as exposure to chemicals or physical factors like ultraviolet radiation among others, endogenously generated reactive oxygen species (ROS), errors in DNA replication, or defects in specific DNA repair mechanisms (5,6). Elucidating the specific vulnerabilities of cancer cells to different drugs and therapies can facilitate the development of more effective and personalized cancer treatments (7–10). Therefore, in addition to recognizing specific mutational signatures, it is also crucial to understand the processes responsible for them. Continuous research is expected to lead to full understanding of the individual mutational processes in sufficient detail to maximize their potential use in biomedical applications.

Ionizing radiation has been known to be mutagenic ever since Muller demonstrated that X-rays induce mutations in Drosophila sperm (11). The extensive studies on cancer incidence and mortality in atomic-bomb survivors from Hiroshima and Nagasaki have established exposure to ionizing radiation as an independent risk factor for many different types of cancer (12–14). It is now clear beyond doubt that high doses of ionizing radiation from medical radiation therapy or nuclear accidents can increase the risk of developing cancers such as leukemia, thyroid cancer and breast cancer (12–15). In addition, a large body of animal studies has also demonstrated that exposure to ionizing radiation can increase the risk of developing cancer in a dose-dependent manner (16–19). Numerous studies using murine germlines and *Arabidopsis thaliana* cells have demonstrated that ionizing radiation can cause various types of mutations, among which enrichment of indels (20–24). It is important to note that the risk of developing radiation-induced cancer depends on a complex multitude of factors, such as radiation dose and the dose-rate, duration of exposure, age at exposure, the cell types in the tissue and the level of their differentiation, as well as the individual’s genetic susceptibility. Ionizing radiation can damage DNA through direct or indirect ionization mechanisms, resulting in various types of DNA lesions including damage to single bases, single-stranded and double-stranded DNA breaks (SSBs and DSBs) (25). The type of mutations induced in the process of mutagenesis is thought to have a decisive role in cancer risk. DSBs and clustered DNA lesions induced by ionizing radiation are considered to be the most severe types of DNA damage, having the highest toxicity to cells and mutagenicity (26). Notably, the repair of DSBs and cluster damage can be substantially delayed or compromised, which determines the higher likelihood of mutations and further genomic instability (27,28). Overall, comprehending the mutational signatures resulting from exposure to ionizing radiation is a formidable undertaking, which despite an ample amount of existing studies, including whole genome sequencing, is likely to continue in the foreseeable future.

Radiation exposure can be detected and measured down to very low levels, and according to the linear non-threshold (LNT) hypothesis, radiation exposure even at low doses is associated with a finite risk of cancer induction (29). Unlike other mutagens, radiation can produce complex DNA damage through both the direct ionization of DNA molecules or by indirect oxidation via reactive oxygen species (ROS), which are also continuously generated in living cells under physiological conditions (30,31). As a product of normal oxygen metabolism, ROS are a major source of endogenous DNA lesions *in vivo*, and their levels can substantially increase even further by exposure to various environmental stressors and inflammation (32). It has been postulated that one-third of radiation-induced DNA lesions result from direct radiation absorption and two-thirds result from the ionization of water molecules and the subsequent production of ROS (33). The chemical nature of the DNA lesions generated by ROS is indistinguishable between spontaneously generated ROS and indirect radiation effects. The rate of production of endogenous DNA lesions has been estimated for SSBs and apurinic/apyrimidinic sites (AP-sites or abasic sites) at more than 2,000 and 10,000, respectively; and for 8-oxo-7,8-dihydroguanine (8-oxo-G, also known as 8-hydroxyguanine) at 1,000 to 100,000 per cell per day (31,34–38). These amounts are equivalent to those induced by gamma-radiation delivered at dose rates of at least 1 to 10 Gy per day (31). In particular, 8-oxo-G is known as a major DNA lesion generated by ROS, and it is well known that 8-oxo-G on the replicating strand leads to misincorporations of adenine opposite the lesion at a rate of 10-75 %, subsequently resulting in G:C to T:A (C:G to A:T) transversion mutations (37,39,40).

Present-day large-scale integrated genomic analyses are able to distinguish the radiation-induced events associated with specific types of cancer (41). However, the fact that indirect ionization via ROS is the predominant mechanism by which radiation induces multiple types of lesions in genomic DNA is the reason why it is exceedingly difficult to distinguish between most radiation-induced cancers and spontaneous cancers. In recent years, whole-genome sequencing studies of clonal lines from somatic cells in various tissues of mice and humans have identified spontaneous somatic mutations (42,43). In these studies, C:G to T:A transitions and C:G to A:T transversions at non-CpG sites were observed as the majority of spontaneous mutations in addition to C:G to T:A transitions at CpG dinucleotides that occur from the deamination of 5-methylcytosine to thymine, which may reflect cell division (2). Furthermore, a recent study of somatic mutations using clonal cell populations from hematopoietic stem cells of whole-body X-ray-irradiated mice demonstrated that radiation exposure significantly increased small deletions and insertions, and also both C:G to T:A transitions and C:G to A:T transversions at non-CpG sites in a dose dependent manner (44). Moreover, experiments on the *gpt* delta mouse system with potassium bromate used as a ROS-inducing agent have also suggested that the same mutations at non-CpG sites can be induced by ROS (45). The mechanisms behind the mutations induced by ROS seem to be well explained with the damage or modification of either G or C bases, and their repair processes and errors. However, the prominent mutation hotspots are expected to be influenced by many factors in addition to the sequence context and gene’s function, such as the DNA repair process, chromatin structure, cell cycle, and more.

Recently, we reported the development of an advanced mutagenesis assay using a random-DNA-barcoded *supF* shuttle vector library with non-SOS-induced *E. coli*, which we called “*supF* NGS assay” (46). In the study, we demonstrated that the *supF* NGS assay allowed us to quickly accumulate a very substantial amount of data and to contribute some new findings regarding UV-induced mutation signature and mutational processes in mammalian cells, which was surprising given that UV-associated mutagenesis have been studied in great depth and is considered well-known. Thus, we demonstrated that the *supF* NGS assay has advantages in sensitivity and throughput; it is also especially useful as a damage-specific mutagenesis assay, since different DNA lesions of interest can be introduced at different sites and sequences and in different numbers in shuttle vectors using *in vitro* construction. Various experiments can be designed based on environmental mutagen exposure and/or genetic perturbations, which will help to elucidate the mechanisms behind mutational signatures. In a previous study, we have demonstrated that chronic Cs-137 gamma-irradiation at a low dose-rate of 1 Gy per day efficiently induces senescence via ROS-mediated activation of an ATM-P53-P21 pathway in human fibroblast cells, but not in the many other types of cell lines that we tested (47,48). Unlike fibroblasts, other cell lines showed continuous growth with little mitotic cell death under conditions of chronic irradiation. Despite this different outcome, radiation-induced DNA damage detected by gamma-H2A.X and 53BP1 were similarly accumulated in a dose-rate dependent manner in all cell lines. In this study, using our recently developed *supF* NGS assay under experimental conditions of chronic Cs-137 gamma-irradiation, we have attempted to address the mutational signatures characteristic for continuous exposure to low levels of radiation.

## RESULTS AND DISCUSSION

### Mutation frequencies were increased in response to chronic gamma-irradiation at a dose-rate of 1 or 2 Gy per day

In order to test whether we can investigate the mutational signatures associated with exposure to chronic gamma-irradiation at a dose-rate as low as 1 Gy per day in mammalian cells, at first the *supF* mutant frequencies were analyzed by using the RF01 *supF*-mutation indicator *E. coli* strain (49) and a conventional *supF* mutagenesis assay, which compares the number of colonies on selection plates versus titer plates. Briefly, the random 12-nucleotide barcoded (N_12_-BC) pNGS2-K2 shuttle vector libraries containing the *supF* gene were constructed to be used with the NGS mutagenesis assay (Figures 1A and 1B). The constructed libraries were transfected into the human osteosarcoma U2OS cells, allowing the libraries to generate mutations with their replication in the cells. The cells were cultured for 2 days in a 5% CO_2_, 37°C incubator at a dose-rate of 0, 1 or 2 Gy per day using a Cs-137 gamma-ray source (Figure 1C) (47). The shuttle vector libraries replicated in the cells during the 2 days of incubation were purified by digestion of unreplicated vectors with *N*^6^-methyladenine-dependent restriction endonuclease *Dpn* I, and then were electroporated into RF01 and seeded on either titer or selection plates in appropriate dilutions. The *supF* mutant frequencies were obtained by counting colonies grown on the plates. The results from three independent transfection experiments indicate that chronic gamma-irradiation at a dose-rate of 1 to 2 Gy per day can produce *supF* mutants with frequencies 0.5 to 1.5 × 10^-3^ higher than the spontaneous one (Figure 1D).

**Figure 1.**
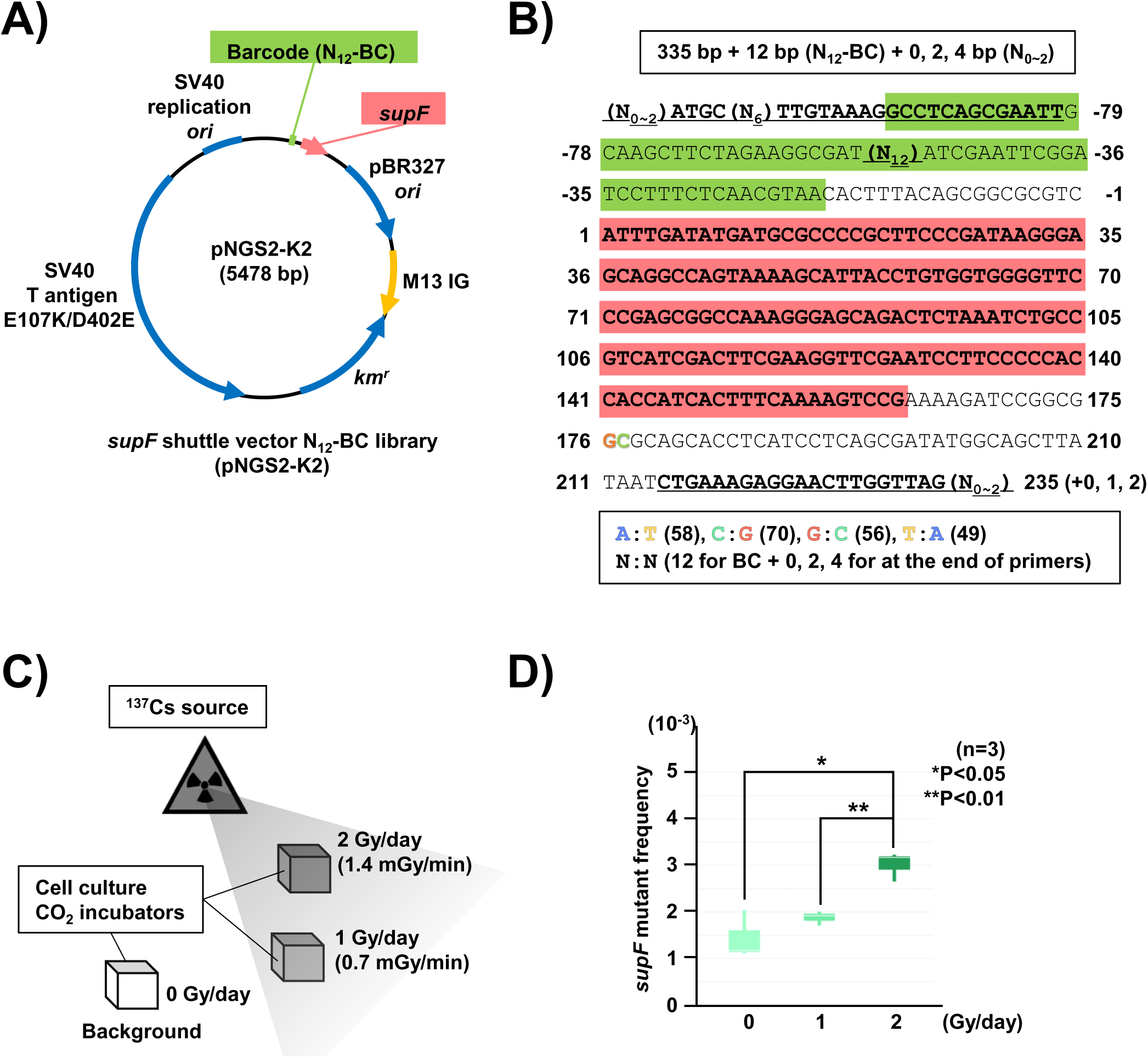
Increase of *supF* mutation frequencies in response to chronic gamma-irradiation. (A) Schematic map of the *supF* shuttle vector N_12_-BC library (pNGS2-K2). The pNGS2 encodes the amber suppressor tRNA (*supF*) gene, the TP53-/Rb-binding-deficient mutant SV40 large T antigen (SV40 T antigen E107K/D402E) gene, the SV40 replication origin (SV40 replication *ori*), the pBR327 origin of replication (pBR327 *ori*), the M13 intergenic region (M13 IG), the kanamycin-resistance gene (*km^r^*), and the randomized 12-nucleotide barcode DNA sequence (N_12_-BC). The N_12_-BC is inserted before the sequence of the *supF* gene. (B) The nucleotide sequence for NGS analysis encoded in pNGS2-N_12_-BC libraries (335 bp +12 bp for the N_12_-BC). The underlined letters indicate the primer set for PCR amplification. Primer sets contain 0, 1, or 2 random nucleotides at the 5’ end to ensure proper signal detection for amplicon sequencing. A pre-designed 6-nucleotide sequence (N_6_) in the forward primers serves as an index sequence for multiplexed NGS. The sequence with green background represents commercially synthesized oligonucleotides used for the insertion of N_12_-BC sequences (N_12_) into the pNGS2. The sequence with red background indicates the *supF* gene starting at position 1. (C) A diagram of the low-dose-rate gamma-radiation facility at the Radiation Research Center for Frontier Science (RIRBM, Hiroshima University) equipped with a 1.11 TBq ^137^Cs source. Cell cultures were maintained in conventional incubators at 37 °C, 5% CO_2_ under gamma-irradiation conditions at the indicated dose-rates (1 or 2 Gy/day). (D) Box plot of the *supF*-mutant frequency in response to gamma-irradiation. The *supF* mutant frequency was determined by the conventional *supF* assay by dividing the number of colonies grown on selection plates by the number of colonies grown on titer plates (n=3).

### Chronic gamma-irradiation induced strand-biased mutations at C:G sites were observed in the mutation spectra from colonies grown on *supF*-selection plates

Next, the mutation spectra were analyzed using the *supF* NGS assay, the procedure for which was described in detail in our previous report (46). Briefly, to obtain a substantial amount of data, including data on spontaneous mutations that occurred endogenously in cells, roughly two thousand colonies of RF01 grown on *supF*-selection plates and containing the N_12_-BC shuttle vectors with a variety of mutations in the *supF* gene were harvested for each sample. Using the libraries purified from the harvested colonies as PCR templates, DNA fragments including the *supF* gene (Figure 1B) were amplified by two independent PCRs with the index-tagged primers and analyzed by the MGI DNBSEQ platform. The sequencing data containing the mutation spectra were classified and error-corrected by using the N_12_-BCs and the variant frequencies (VFs). The VFs are the frequencies of identical variant call(s) found in a unique N_12_-BC. Variant sequences exceeding 0.4 VF can be assumed to be true mutations and not errors that have occurred in the process of NGS analysis. In addition, to eliminate PCR errors from the data, only the identical N_12_-BC with recurring mutations between two independent PCR samples were extracted and used for analysis. The number of mutations at each position in the sequence was plotted and shown in Figure 2A. The data from three independent transfections (TF1, TF2, and TF3) appear to have good reproducibility, and the mutations were located in the region of the *supF* gene (positions from 56 to145) and its promoter region (position -15 is the first nucleotide of the promoter), consistent with the results previously reported by many groups, including us (46). The positional bias for the number of detected mutations was caused by the *supF* selection process, which relies on the destruction of the function of transfer RNAs. This positional bias was observed in irradiated samples as well, as chronic gamma-irradiation seems to induce mutations at the same positions and some other specific positions (Figure 2A). In order to compare appropriately the mutation spectra and frequencies across samples, one hundred N_12_-BCs with at least one mutation from the group with the highest number of reads, or the maximum number of available N_12_-BCs with mutations were subjected to further analysis. These approximately one hundred unique N_12_-BCs for each sample contain 176-211 mutations, indicating that more than one mutations were present in some of the N_12_-BCs (Figure 2B). Single nucleotide substitutions (SNSs) were the predominant type of mutation, and the proportion of SNSs was not significantly altered by irradiation (Figure 2C). It is important to note that SNSs had occurred mainly from C:G (Non-transcribed:Transcribed strand, respectively) or G:C, where C and G were substituted into various bases (Figure 2D). Even though not significant, it was observed consistently that chronic gamma-irradiation induced mutations mainly from C:G in a dose-rate dependent manner, suggesting that due to some underlying reason, radiation-induced mutations were strand-biased. The mutational spectra and positions for SNSs were depicted in Figure 2E, and it was confirmed that mutations at spontaneous hotspot positions were increased by irradiation, and C:G mutations were observed more frequently than G:C mutations.

**Figure 2.**
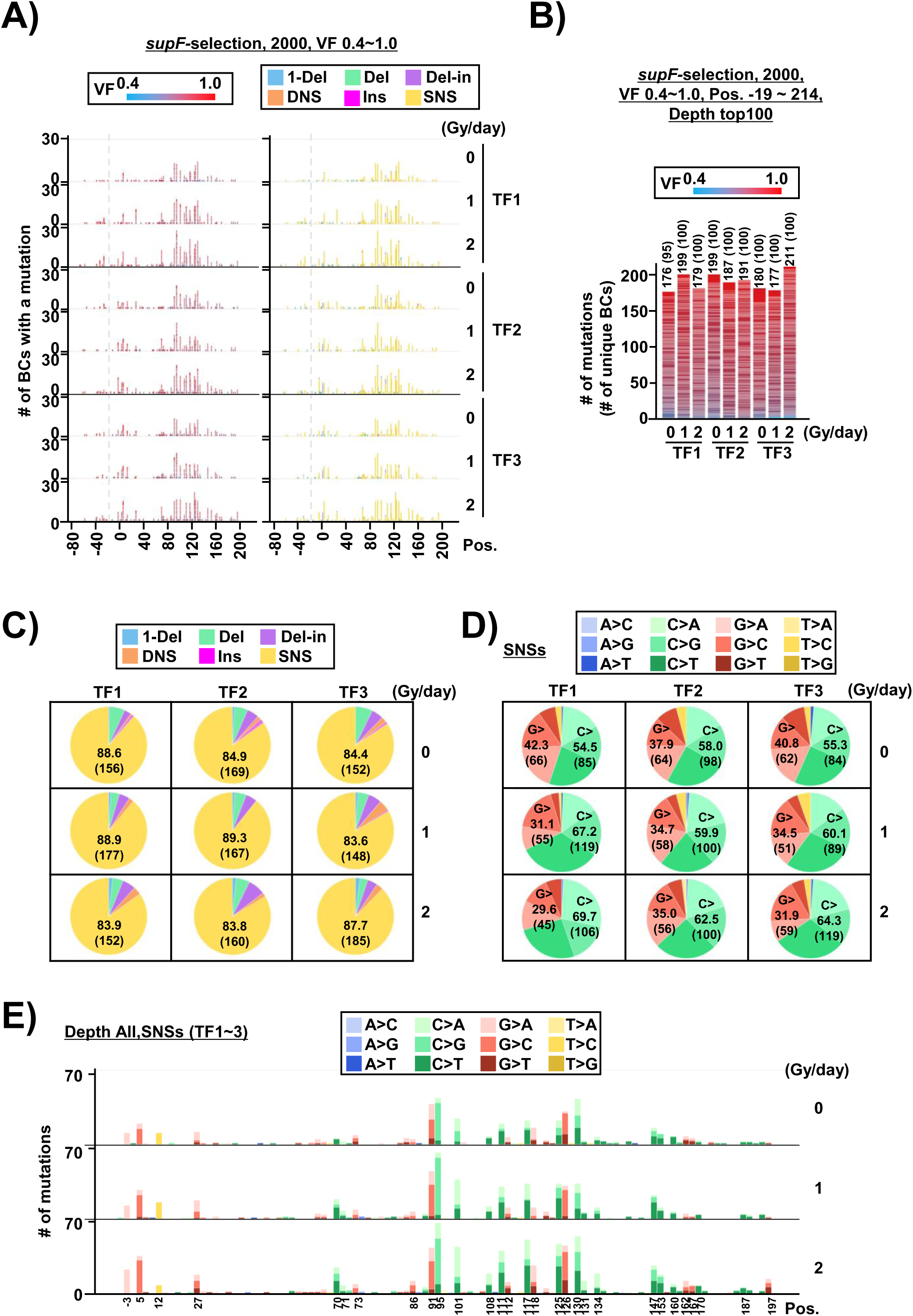
Analysis by the *supF* NGS assay of the mutations induced by chronic gamma-irradiation in cells from 2,000 RF01 colonies grown on *supF*-selection plates (*supF*-selection, VF 0.4∼1.0) (A) Number of N_12_-BC sequences with a variant exceeding VF 0.4 according to their nucleotide positions (Pos.) − data from three independent transfection experiments (TF1, TF2, and TF3) and different dose-rates of gamma-irradiation (0, 1, and 2 Gy/day for 2 days). In the left side graph, the colors reflect the variant frequency, as indicated by the heatmap on top. On the right side graph, the variant types are shown in different colors as indicated in the legend. The variant calls are categorized into six types: one-nucleotide deletion (1-Del), deletions larger than one nucleotide (Del), deletions with insertions (Del-in), dinucleotide substitutions (DNS), insertions (INS), and single nucleotide substitutions (SNS). (B) Stack bar graph representing the number of N_12_-BC sequences with mutations from each sample used for further data analysis. Included are the one hundred mutation-bearing N_12_-BCs with the highest number of reads per unique N_12_-BC or the maximum number of available N_12_-BC with mutations. The number in parentheses refers to the number of unique N_12_-BC sequences with mutations. (C) Pie charts of the proportions of mutation types shown in different colors as indicated in the legend on top. The percentage and number (in parentheses) of SNSs are provided in each pie chart. (D) Pie charts of different base substitutions as proportions of SNSs. The individual base substitutions are shown in different colors as indicated in the legend on top. The percentage and number (in parentheses) of substitutions from guanine (G>) or cytosine (C>) are provided in each pie chart. (E) Number of SNSs according to their nucleotide position (Pos.) at different dose-rates of gamma-irradiation (0, 1, and 2 Gy/day for 2 days). The numbers represent data combined from three transfection experiments. The individual base substitutions are shown in different colors as indicated in the legend on top.

### The mutation spectra of C:G substitutions differ between single mutations per N_12_-BC and multiple mutations per N_12_-BC

In order to address the mechanisms underlying the C:G strand-biased mutations induced by irradiation, the individual SNSs were analyzed in more detail. As shown in Figure 2E, the majority of mutations from C or G were detected at specific positions that are considered required for the *supF*-selection. The detected mutations were classified as single mutations in a unique N_12_-BC only when they were selected as a standalone single mutation, while the multiple mutations per N_12_-BC included those that were co-detected along with the aforementioned single mutations. It should be noted that the multiple mutations were able to be detected spontaneously and the frequencies of mutational events were as low as 10^-3^. These facts implied indirectly that most multiple mutations were produced in a single process, thereby, it was anticipated that some of the single mutations in the *supF*-responsible region could be distinguished as mutations directly produced from DNA damage induced by irradiation. Therefore, the mutations detected at different radiation dose rates were presented separately for the single or more than one mutation per N_12_-BC (Figure 3). The percentages of N_12_-BCs with either single or multiple mutations were not significantly altered by irradiation (Figures 3A, 3B, and 3C), suggesting that the strand-biased mutation induction by irradiation took place in both cases. For an in-depth look, the mutations were classified by number per N_12_-BC (1, 2, 3, and 4∼ mutations per N_12_-BC) and depicted according to their positions (Figure 3-figure supplement 1). It was anticipated that single mutations would be exclusively located at specific positions in either the *supF*-tRNA cloverleaf or promoter region, multiple mutations would also include other regions, and C:G mutations would be dominant throughout all regions. However, contrary to expectations, a significant difference was observed between single and multiple mutations in the proportion of mutated bases at C:G/G:C sites (Figure 3D). Based on the data, C:G to G:C transversions were major for single mutations, while C:G to T:A transitions were major for multiple mutations. It appears that exposure to radiation may increase the frequency of C:G to A:T transversions as single mutations and C:G to T:A transitions as multiple mutations, while decreasing the frequency of G:C to A:T transitions in multiple mutations (Figure 3E). There is a remote possibility that by increasing C:G to A:T transversions, which are a characteristic feature of 8-oxo-G, exposure to radiation may become a trigger for the strand-biased C:G to T:A cluster mutations. In previous reports by us and others, the majority of the spontaneous mutations were located at 5’-T<u>C</u>-3’:5’-<u>G</u>A-3’ sites in both DNA strands, which is well known as COSMIC APOBEC (apolipoprotein B mRNA editing enzyme, catalytic polypeptide-like) signatures SBS2/SBS13 (7,50,51). In order to investigate in more detail the SNS mutation spectrum, all possible 192 trinucleotide contexts of mutations were compared between the complementary DNA strands, and it became clear that the SNSs occurred mainly at 5’-T<u>C</u>N-3’:5’-N<u>G</u>A-3’ sites in both DNA strands in all samples (Figure 3-figure supplement 2), without a noticeable difference between non-irradiated and irradiated samples. From this data analysis it became clear that a deeper exploration of these complex mutation spectra was required to address the mutations induced at lower frequency.

**Figure 3.**
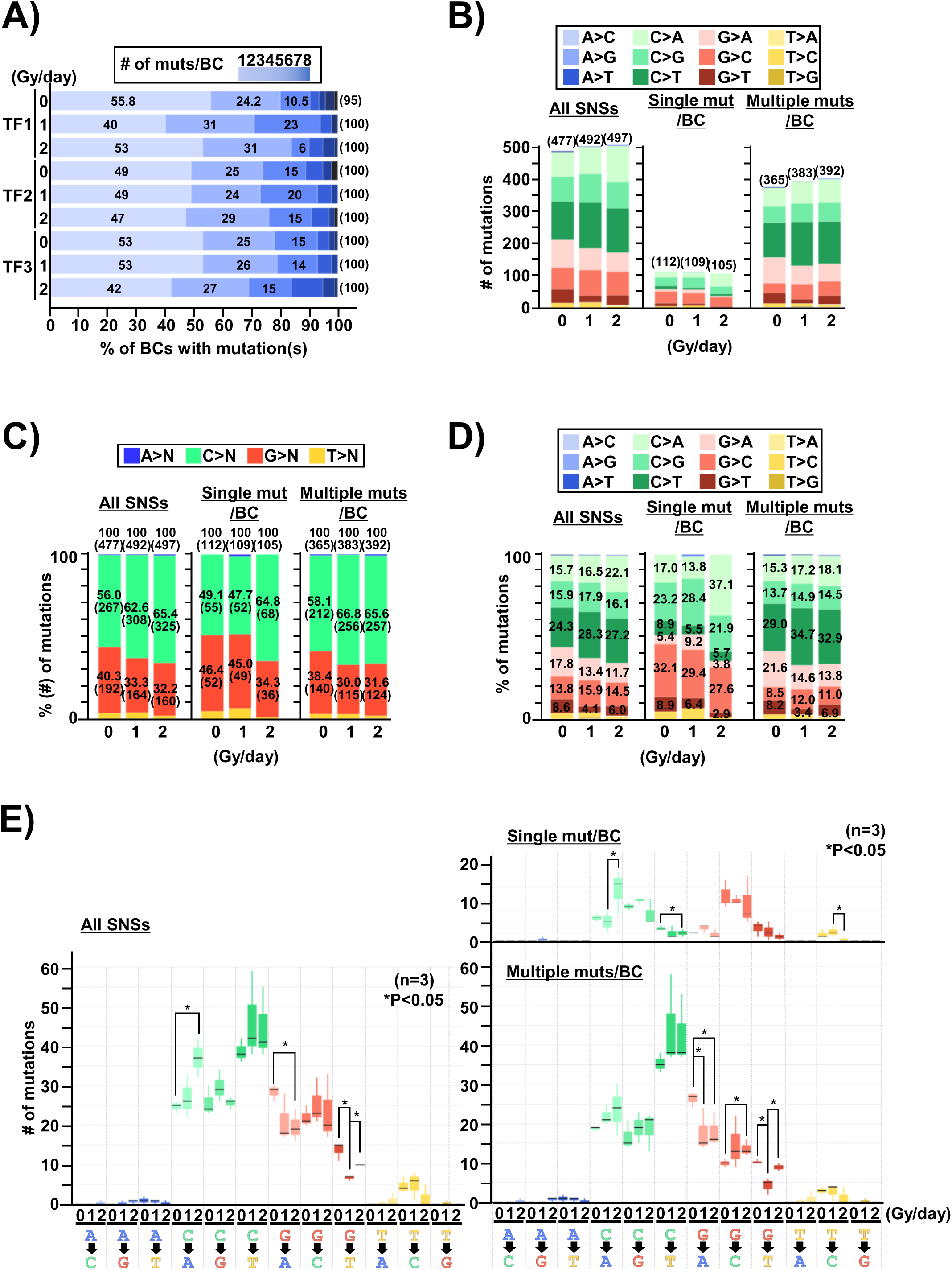
Analysis of base-substitutions in single and multiple *supF* mutations per N_12_-BC (*supF*-selection, VF 0.4∼1.0) (A) Proportion of the number of N_12_-BC sequences with single (1) or multiple (2∼8) mutations from three independent transfection experiments (TF1, TF2, and TF3) at different dose-rates of gamma-irradiation (0, 1, and 2 Gy/day). The percentage of N_12_-BCs with different number of mutations is indicated inside each bar, and the total number of mutations are indicated outside each bar in parentheses. (B) Number of N_12_-BC sequences, combined data from three independent transfection experiments. The number of all SNSs, single mutations per N_12_-BC, and multiple mutations (more than one mutation) per N_12_-BC are separately presented. The total number of SNSs are shown above the bars in parentheses. The proportion of individual base substitutions is shown in different colors as indicated in the legend on top. (C) Stacked bar graph representing the relative proportion of the four bases with reference to the SNSs plotted in (B). The percentage and number (in parentheses) of SNSs from either C or G is provided inside each segment. The four bases are shown in different colors as indicated in the legend on top. (D) Stacked bar graph representing the relative proportion of SNSs, data set identical to (B). The percentage of each SNSs from either C or G is provided inside each segment. The individual base substitutions are shown in different colors as indicated in the legend on top. (E) Box plot of the number of individual SNSs in response to different gamma-irradiation conditions (n=3). The horizontal line in each box plot represents the median. The number of all SNSs is presented in the left-side graph. Single-mutations per N_12_-BC and multiple-mutations per N_12_-BC are separately presented in the right-side graph. The asterisks indicate statistical significance determined by paired student’s *t* test (\**P* < 0.05).

### Mutational spectra of single versus multiple mutations provide evidence for the *supF*-selection bias

Next, the mutational positions within the sequence were ranked based on the number of detected mutations, and the mutational bases and positions with their flanking sequences were shown in Figure 4A. The ranks of single mutations were not significantly changed by irradiation. It proved challenging to identify any discernible features of irradiation, but consistent with the above analysis, in irradiated samples the number of multiple mutations located at 5’-T<u>C</u>-3’:5’-<u>G</u>A-3’ sites was increased more than those at 5’-<u>G</u>A-3’:5’-T<u>C</u>-3’ sites. It became clear that mutations at specific positions such as 111 and 125 can be detected only as multiple mutations, even though they are located in the *supF*-tRNA cloverleaf region (Figure 4A and Figure 4-figure supplement 1A) (52). The method of *supF*-selection has an advantage over other drug selection markers in the range of mutations it allows us to identify; however, there exists a significant bias in the analysis of mutation spectra due to its reliance on the *supF*-tRNA properties. The spectra of single mutations are depicted with the *supF*-tRNA cloverleaf structure in Figure 4B. As described in our previous report, the potential hairpin loop of the quasi-palindromic sequence between positions -8 and 12, GGCGCGTCATTTGATATGATGCGCC, in the promoter region of the *supF* gene may lead to template switching and give rise to complementary cluster mutations (Figure 4-figure supplement 1B). It is obvious that the mutations were distributed unequally at different positions, even at the same 5’-T<u>C</u>-3’:5’-<u>G</u>A-3’ sequence in the cloverleaf-region, suggesting that other factors in addition to the functional bias of the *supF*-gene may be involved in the mutation bias, such as the secondary structure of single-stranded DNA during the replication, repair, or transcription process, etc. Therefore, we predicted the single-stranded DNA secondary structures of the analyzed region by Mfold web server (Figure 4-figure supplements 2 and 3) (53,54), and the number and substituted bases of the identified mutations at each position in the analyzed sequence were depicted separately for single or multiple mutations per N_12_-BC (Figure 4-figure supplements 4-9). However, it proved difficult to find any correlation between mutations and sequences or their structures, which may be due to the *supF*-selection bias.

**Figure 4.**
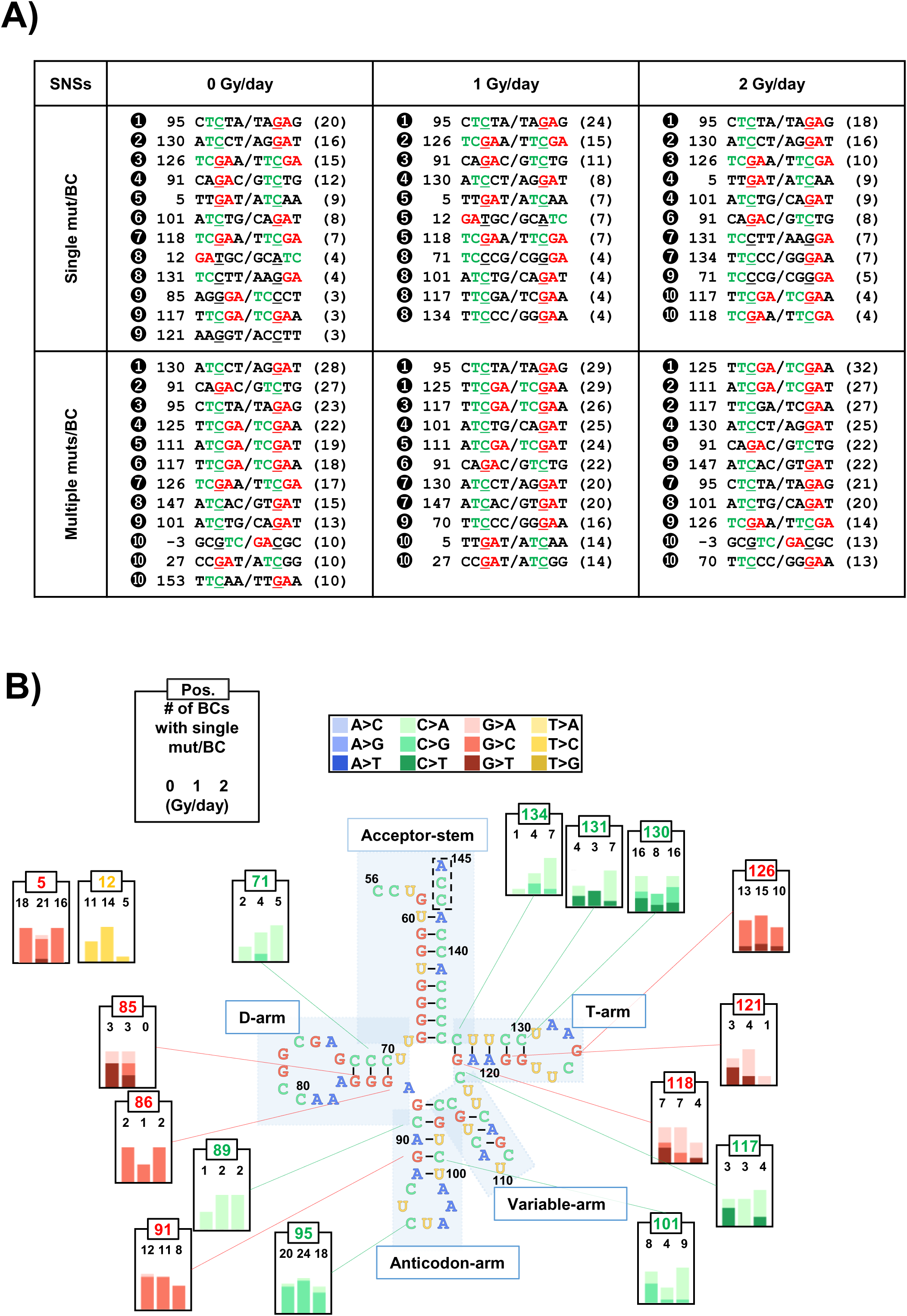
Summary of SNSs induced in cells – data from *supF*-selection. (A) SNSs listed by rank according to the frequency of their detection (number of N_12_-BCs) from the combined data of three transfections for *supF*-selection. The ranks of the SNSs (shown in full black circles) are sorted separately for single mutation or multiple mutations per N_12_-BC and for each radiation dose-rate (0, 1, and 2 Gy/day). Each SNS is denoted as an underlined nucleotide in a pentanucleotide sequence (including two nucleotides upstream and downstream of the mutation, and the sequence of the non-transcribed/transcribed strand). The 5’-TC-3’ sites are shown in green color, and the 5’-GA-3’ sites are shown in red color. The number next to the rank is the position of each mutation in the nucleotide sequence. The number in parentheses represents the number of detected N_12_-BCs with the mutation. (B) The number of single mutations per N_12_-BC according to their positions in the cloverleaf secondary structure of the *supF*-tRNA (positions from 56 to 145). The bar graphs in each frame provide the number of single mutations per N_12_-BC for different radiation dose rates (the numbers are denoted above each bar), and their positions are indicated by dotted lines. The proportion of individual base substitutions is shown in different colors as indicated in the legend on top

### The NGS mutation analysis without *supF*-selection provides mutation frequency and non-biased mutation spectra

The process of *supF*-selection is effective to obtain a substantial amount of data on mutation spectra through concentrating the mutants at a magnitude of thousands. An appropriate *supF*-selection process provides accurate mutant frequency and trends. However, the mutation spectra obtained by *supF*-selection exhibit a strong bias in favor of certain positions and substituted bases that are associated with the function of the *supF* gene. On the other hand, without the selection it is difficult to investigate thoroughly mutation spectra, especially at very low frequencies where it is hard to differentiate from background mutations. We hypothesized that our newly-developed NGS analysis can be applied to samples directly prepared from purified libraries digested by *Dpn* I without transformation into the indicator *E.coli*. Henceforth, the *Dpn* I-digested libraries purified from the cells were directly analyzed by using DNB-seq, following the PCR for multiplex tagging and amplification. The threshold for VF was set to 0.2. As expected, the regions between -67 and -20, where N_12_-BC sequence oligonucleotides were inserted, display very similar pattern of variant sequences with high VFs close to 1.0, reproducible between libraries and independent transfection experiments (Figures 5A and 5B). These sequences turned out to be a suitable internal standard for the NGS analysis. The types of mutations were clearly different between the constructed N_12_-BC region, positions from -67 to -20, and the *supF* gene, positions from -19 to 214 (Figure 5A). The mutational positions were spread relatively evenly in the sequence compared to the data of *supF*-selection shown in Figure 2A, and seemed to have a degree of similarity between samples. As expected, a similar number of N_12_-BCs were classified across all samples, but the number of sequences with variants were significantly different and appeared to be increased by irradiation (Figure 5B). Interestingly, only N_12_-BCs bearing a variant sequence with lower VFs were increased at positions from -19 to 214, and even from -67 to -20. Therefore, we obtained the mutant frequencies separately for the two regions (-67 to -20 and -19 to 214) from the number of reads with a variant sequence (only the highest number of reads with a variant sequence was selected if the unique N_12_-BCs contains multiple variant sequences) divided by the number of total reads (Figure 5C). The results shown that the mutant frequencies were slightly but statistically significantly increased by chronic irradiation, which very closely resembles the results from the *supF* colony assay shown in Figure 1D. The proportions of mutation types and the sequences of SNSs at positions from -19 to 214 remained very similar between non-irradiated and irradiated samples (Figures 5D and 5E), indicating that the strand-biased mutations from C:G observed in the *supF*-selection condition were not present here. However, when the mutational positions were examined in detail, positions 5 and 27 emerged as significant hotspots in all samples, and their prominence among other hotspots was more evident compared to *supF*-selection. It appears that the mutations from G:C at positions at 5 and 27 counteracted the strand-biased mutations from C:G at multiple positions detected in *supF*-selection. When all possible 192 trinucleotide contexts of mutations were compared between the complementary DNA strands (Figure 5-figure supplement 1), it became clear that the results could be a reflection of positional biases. The data set demonstrates that chronic exposure to radiation leads to an increase in the frequency of SNSs at 5’-T<u>C</u>N-3’:3’-N<u>G</u>A-5’ sites in both DNA strands, along with multiple other mutations that exhibit a position-dependent strand bias. The positions where SNSs occurred were essentially the same between non-irradiated and irradiated samples. When the mutational sequences were ranked and compared between NGS analysis without selection (w/o-*supF*-selection) and *supF*-selection (Figure 5-figure supplements 2A and 2B), novel mutational hotspots were revealed, including positions 5, 27, 111, and 125, among others. These mutations could not be identified from *supF*-selection where they were masked by other position-biased mutations, although they were detected as multiple mutations per N_12_-BC as shown in Figure 4A.

**Figure 5.**
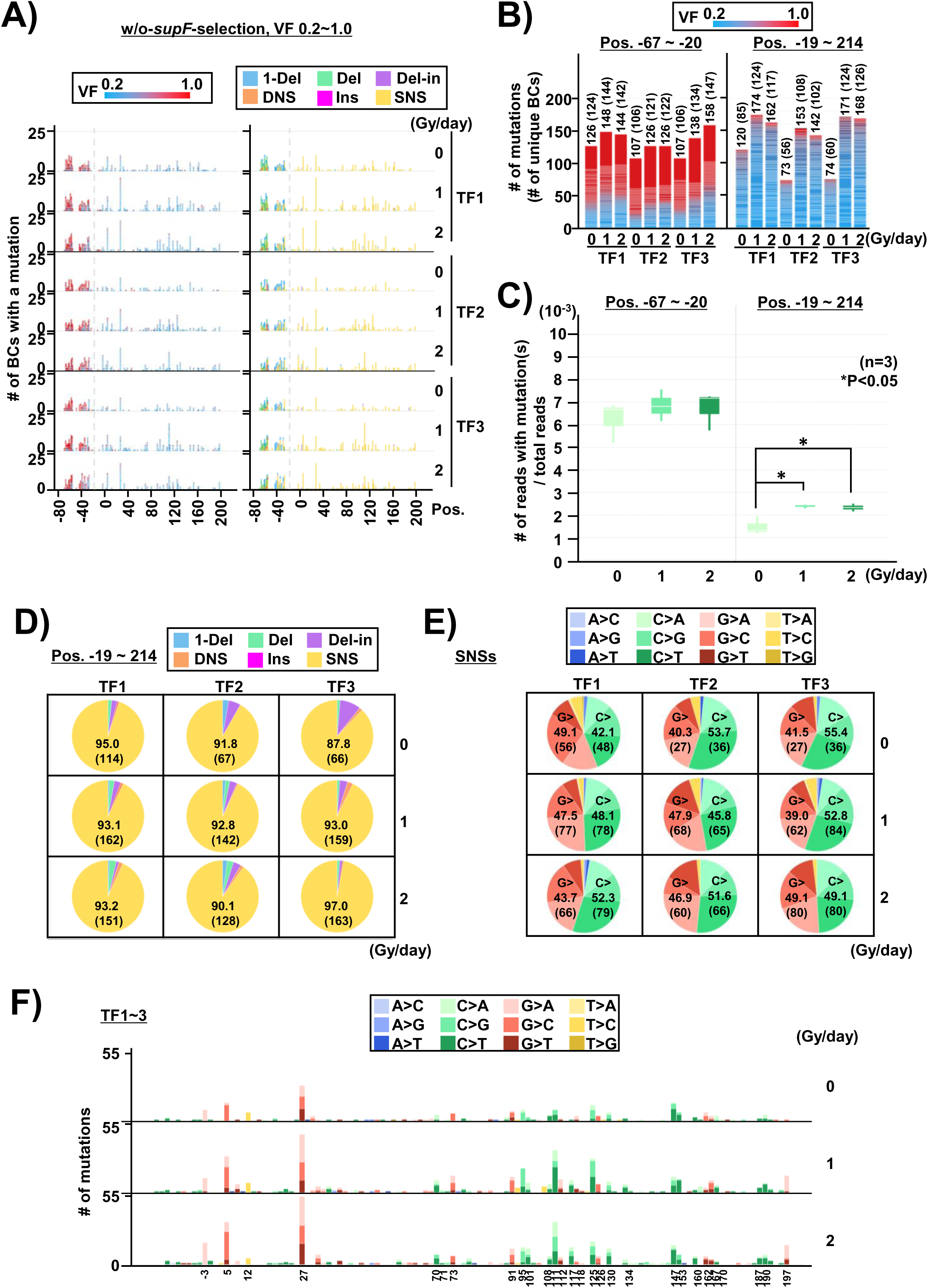
Analysis by *supF* NGS assay of the *Dpn* I-treated libraries extracted from cells (w/o-*supF*-selection, VF 0.2∼1.0) The layout of the figure is essentially analogous to Figure 2, only this time the NGS samples were prepared from the *Dpn* I treated libraries (used for the experiments in Figures 2 : the graph on the left side represents positions from -67 to -20, and the graph on the right side – positions from -19 to 214 containing the *supF* gene. (C) Box plot of the mutant frequency under different irradiation conditions calculated from NGS data. The mutant frequency was determined by the number of reads harboring mutation(s) vs the total number of reads, as described in the text. The data is separated into two graphs: positions from -67 to -20 (left side), and positions from -19 to 214 containing the *supF* gene (right side). The results were obtained from three independent transfection experiments. The asterisks indicate statistical significance determined by paired student’s *t* test (\**P* < 0.05). (D) Pie charts of the proportions of mutation types (refer to the legend in Figure 2C). (E) Pie charts of the proportions of SNSs represented by different base substitutions (refer to the legend in Figure 2D). (F) Number of SNSs according to their nucleotide position (refer to the legend in Figure 2E).

### Positional bias contributes to the discrepancy of the data on strand-biased mutations between *supF*-selection and without *supF*-selection

The NGS data from the w/o-*supF*-selection experiments was analyzed by the number of mutations per N_12_-BC (Figure 6A). About 20 to 30 percent of N_12_-BCs contained multiple mutations, and those seemed to be slightly increased by irradiation, although the limitations of the N_12_-BC-variant sequence classification in the w/o-*supF*-selection condition bring a degree of uncertainty. In fact, both single and multiple mutations per N_12_-BC appeared to be increased by irradiation (Figure 6B), and both were strand-biased; however, the radiation-induced increase occurred from G:C for single mutations, and from C:G for multiple mutations per N_12_-BC (Figures 6C and 6D). The mutations were then depicted within the sequence and with the predicted secondary structure in Figure 6-figure supplements 2 and 3, analogous to Figure 4-figure supplements 4 and 5. For obvious reasons, the number of detected mutations at a single position was relatively less in the w/o-*supF*-selection condition compared to *supF*-selection; on the other hand, the w/o-*supF*-selection analysis could detect mutations at a same number of positions as the *supF*-selection, with some discrepancies. The data between *supF*-selection and w/o-*supF*-selection were compared in Figure 7. It may appear that the data regarding the effect of irradiation is in disagreement between *supF*-selection and w/o-*supF*-selection, but in reality the data were very consistent and highly compelling if strand bias and positional bias for spontaneous mutational hotspots at 5’-T<u>C</u>-3’:5’-<u>G</u>A-3’ sites are considered. The SNSs are the major mutations at positions from -19 to 214 which are generated in cells through replication. The positions of SNSs are relatively evenly spread in the sequence for all samples, but there exist hotspot sites that are shared with background mutations. So far, from the data two noteworthy points have emerged. First, chronic gamma-irradiation for 2 days at a dose-rate as low as 1 Gy per day induces a significant amount of mutations, which are very similar to the spontaneously induced mutations. This may indicate that chronic gamma-irradiation generates the same initial events that are responsible also for spontaneous mutations, that these events are the root cause of the strand bias associated with the 5’-T<u>C</u>-3’:5’-<u>G</u>A-3’ sequence in both strands, and are therefore an essential factor that influences the mutational frequencies and the substituted bases. This can be addressed by the experiments without *supF*-selection because, as already discussed, the strong bias associated with *supF*-selection obstructs the elucidation of these matters. As a second point, the estimated numbers of DNA lesions expected to be induced by gamma-irradiation, including the direct and indirect effects, are almost equal to the number of detected mutations. This may indicate that a DNA lesion directly causes a mutation without repair, or that a single DNA lesion can trigger mutagenesis and lead to amplification of mutations.

**Figure 6.**
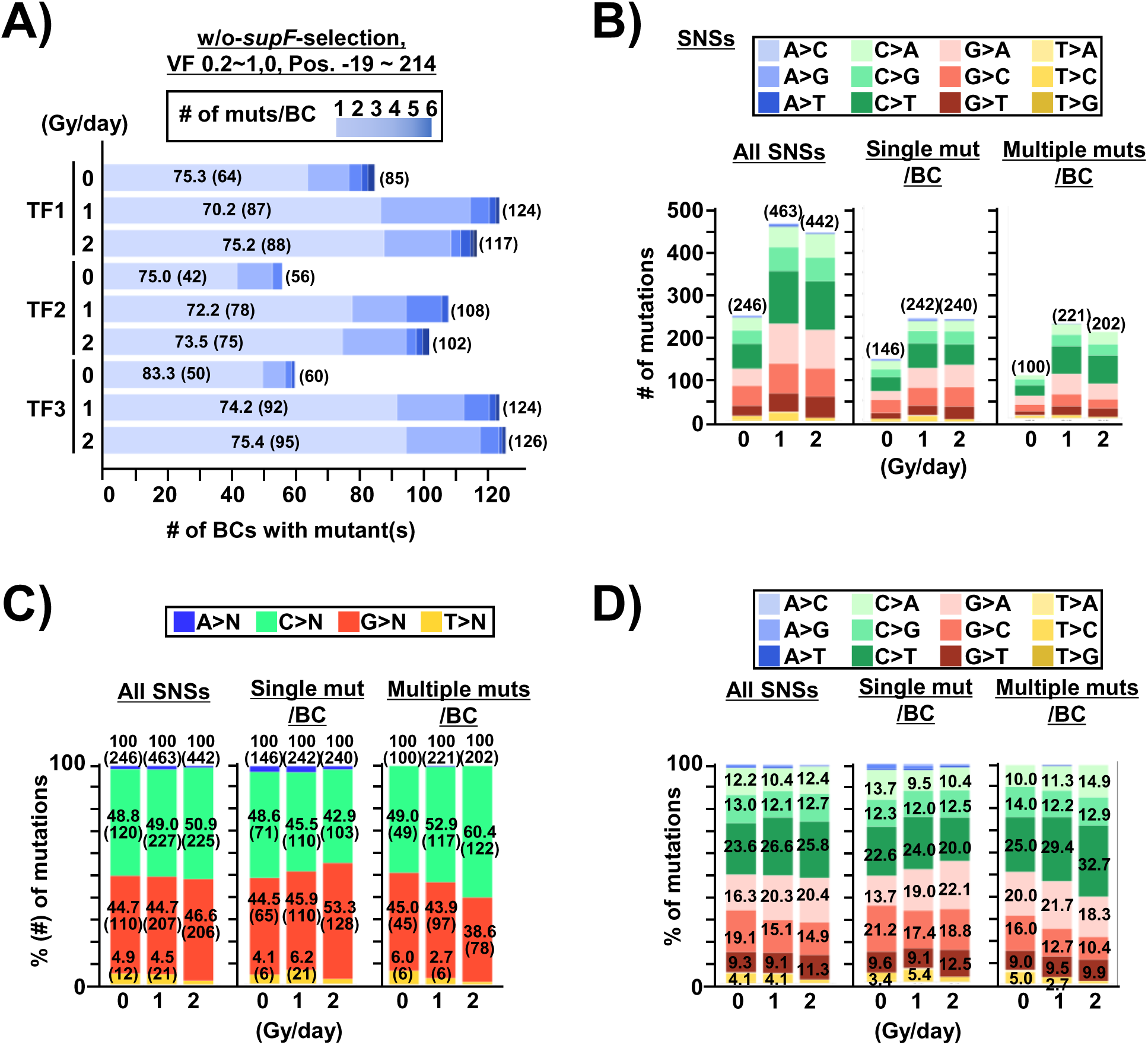
Analysis of base-substitutions in single and multiple *supF* mutations per N_12_-BC in response to chronic gamma-irradiation (w/o-*supF*-selection, VF 0.2∼1.0) The layout of the figure is essentially analogous to Figure 3, only this time the NGS samples were prepared from the *Dpn* I-treated libraries (used for the experiments in Figures 2 and 3) before transformation into RF01. (A) N_12_-BC sequences with single (1) or multiple (2∼8) mutations from three independent transfection-experiments (TF1, TF2, and TF3) at different dose-rates of gamma-irradiation (0, 1, and 2 Gy/day). The percentage and number (in parentheses) of N_12_-BCs with a single mutation per N_12_-BC are presented inside each bar, and the total number of mutations is indicated next to the bars in parentheses. (B) Bar graph of the combined number of N_12_-BC sequences from three independent transfection-experiments (refer to the legend in Figure 3B). (C) Stacked bar graph representing the relative proportion of the four bases in SNSs (refer to the legend in Figure 3C). (D) Stacked bar graph representing the relative proportion of SNSs (refer to the legend in Figure 3D).

**Figure 7.**
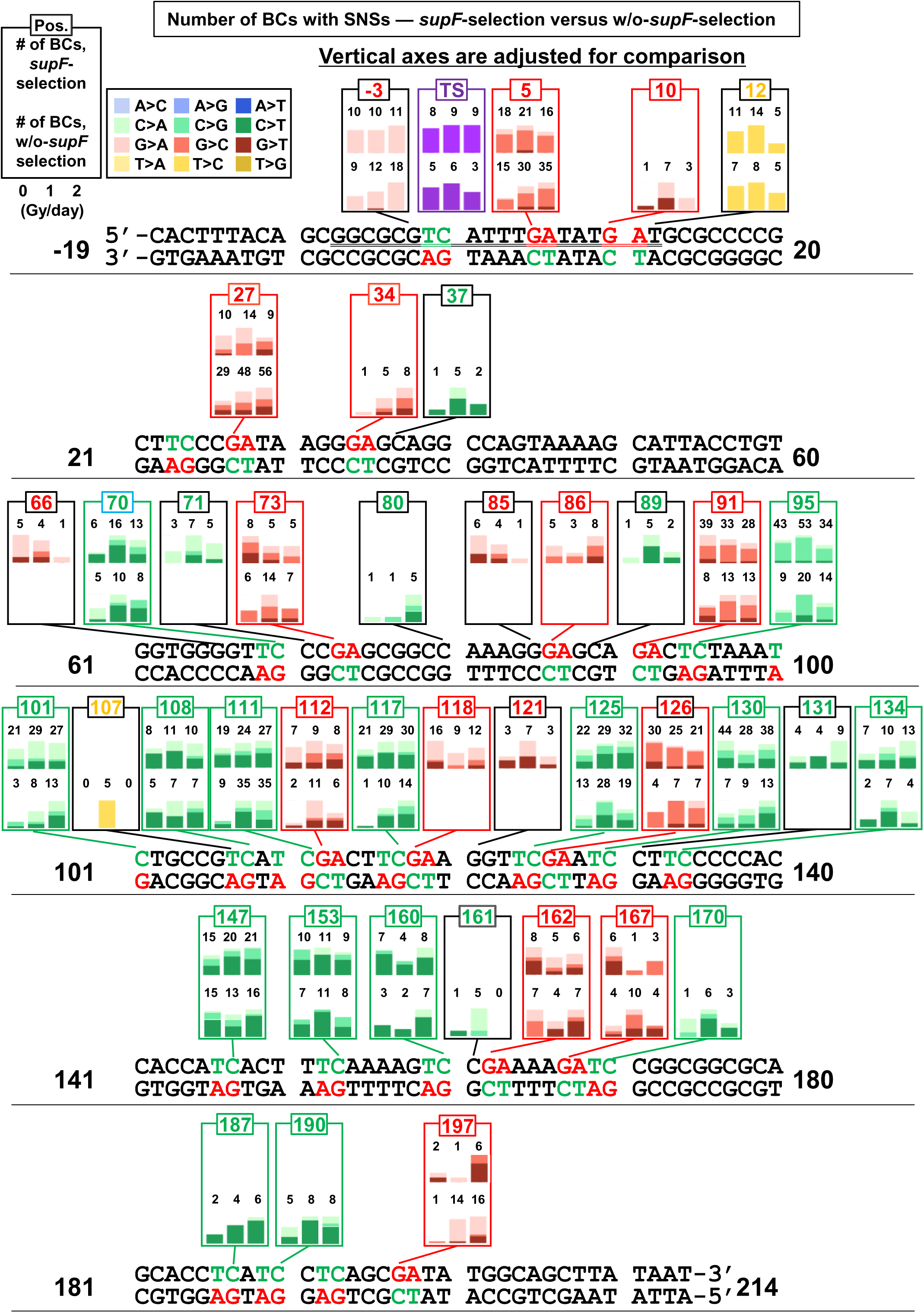
Analysis of SNSs by base substitutions according to positions in the sequence and dose-rates of irradiation (*supF*-selection versus w/o-*supF*-selection) Bar graphs depicting the number of SNSs at the indicated positions on the analyzed sequence (positions from -19 to 214) − combined data from three transfection-experiments (TF1, TF2, and TF3). Each frame depicts the number of SNSs for *supF*-selection (bars on top) and w/o-*supF*-selection (bars on bottom). The colors of the bars and segments of stack bars reflect the type of SNSs as indicated in the legend on the top-left side of the figure. The 5’-TC-3’ and 5’-GA-3’ sites in the sequence and their positions are shown in green and red, respectively.

### A single 8-oxo-G efficiently induces mutations apart from the damaged base, referred to as action-at-a-distance mutations

The 8-oxo-G is known as a major DNA lesion generated by ROS produced endogenously or by gamma-irradiation (37,38). Previously, by using the conventional *supF* assay, we have reported that an 8-oxo-G can serve as a trigger for mutations at 5’-T<u>C</u>-3’:5’-<u>G</u>A-3’ sites, referred to as action-at-a-distance mutations, although the underlying mechanism still remains unclear (51,55). Here, to investigate the effects of 8-oxo-G in inducing SNSs and its potential contribution to radiation-induced mutagenesis, we introduced a single 8-oxo-G outside the region of the *supF* gene, at positions 176 or 177 (refer to Figure 1B), which has no effect on the tRNA function of the *supF*-gene, in a series of pNGS2 shuttle vectors. The schematic diagram of eight different shuttle vectors constructed by the *in vitro* enzymatic method (56) − pNGS2-K1, -K2, -K3, and -K4, containing either non-damaged guanine (G) or 8-oxo-G is depicted in Figure 8A. These vectors contain an M13 intergenic region with opposite orientations relative to the *supF* gene, which allows us to incorporate 8-oxo-G at specific sites in the opposite strands of the shuttle vector library; the vectors also contain a SV40 replication origin on opposite sides relative to the *supF* gene, which changes the direction of replication forks and/or transcription (46). The two different N_12_-BC libraries (with inserted G or 8-oxo-G) for pNGS2-K1 to -K4 were independently prepared, and it was confirmed that no mutation could be detected by NGS analysis. In the preparation process for NGS samples, KOD-one polymerase synthesized a strand complementary to both the damaged and non-damaged sites evenly, which fortuitously allowed for the insertion of 8-oxo-G at positions 176 and 177 in the *in vitro* synthesized library to be confirmed by its detection as thymine instead of guanine in approximately half of the reads (Figure 8-figure supplement 1A). The *supF* mutant frequencies for each library containing either G or 8-oxo-G were analyzed following transfection into cells by using the conventional *supF* assay. As shown in Figure 8B, the *supF* mutant frequencies were increased by the insertion of 8-oxo-G in the region outside the *supF* gene, although the effect was statistically significant only in pNGS2-K3 and -K4. In parallel, the libraries after being purified from the cells and digested by *Dpn* I, were analyzed by NGS, as in w/o-*supF*-selection experiments. The nucleotide positions and types of mutations based on the combined data from three transfections are shown in Figure 8C. Consistent with the data on mutant frequencies obtained from the conventional *supF* assay, more mutations were detected after insertion of 8-oxo-G compared to the control insertion of G for all libraries (Figure 8C). Referring back to what was mentioned in Figure 5A, the region between -67 and -20 showed very similar pattern of variant sequences between G and 8-oxo-G. The number of mutations in the region from -19 to 214, excluding positions 176 and 177 with the introduced G or 8-oxo-G, were clearly increased. Notably, the patterns of mutations following insertion of G were similar in all libraries, while it appears that upon insertion of 8-oxo-G two distinct patterns were observed – one in pNGS2-K1 and -K4, and another in pNGS2-K2 and -K3. The number of N_12_-BCs with at least one mutation for regions -67 to -20 and -19 to 214 indicate that a single 8-oxo-G caused the generation of other SNSs in the region (Figure 8D). The frequency of mutations induced at positions 176 and 177 with inserted 8-oxo-G was only in the order of 10^-3^ reads, suggesting that the inserted in the library 8-oxo-G is efficiently repaired in the cells (Figure 8-figure supplement 1B). The mutant frequencies calculated as a ratio between the number of reads with mutations and the number of total reads for positions from -19 to 214, except for positions 176 and 177, were relatively lower compared to those obtained from the conventional *supF* assay but revealed very similar trends (Figures 8E versus 8B). This is consistent with the irradiation experiments using the conventional assay versus NGS, shown in Figures 1D and 5C. It is possible that the loss of data comes from reads containing true mutations with VFs below 0.2, which were excluded in order to avoid false positives from NGS errors.

**Figure 8.**
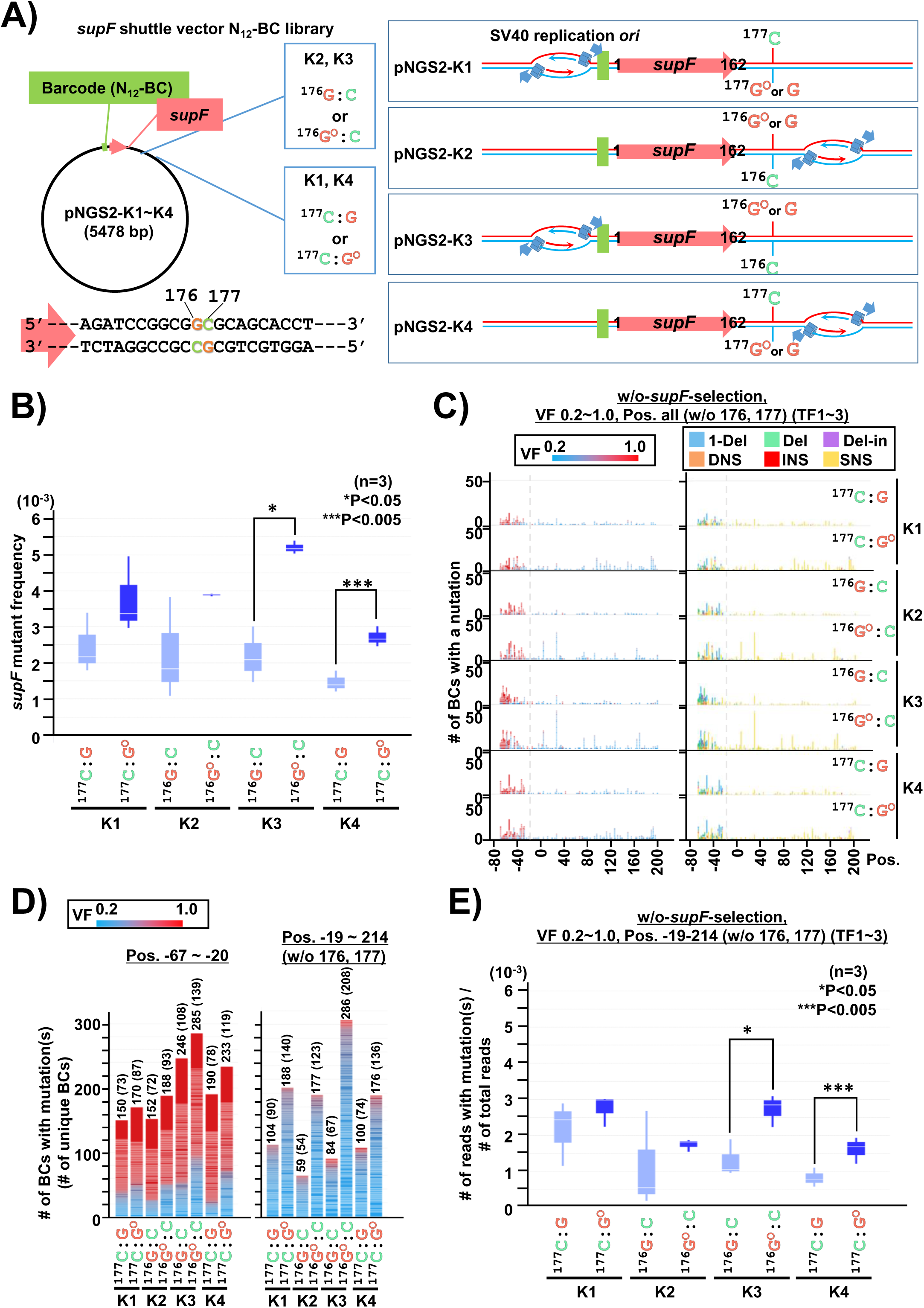
The presence of a single 8-oxo-G lesion induces action-at-a-distance mutations (w/o-*supF*-selection, VF 0.2∼1.0) (A) Schematic map of the N_12_-BC libraries (pNGS2-K1∼K4) with an inserted 8-Oxo-7,8-dihydroguanine (8-oxo-G, G^O^) or a lesion-free G (G). An 8-oxo-G or G was inserted outside the *supF* gene at position 176 for pNGS2-K2 and pNGS2-K3 (represented as ^176^G:C for a lesion-free G-inserted and ^176^G^O^:C for an 8-oxoG-inserted), or at position 177 for pNGS2-K1 and pNGS2-K4 (represented as ^177^C:G for a lesion-free G-inserted and ^177^C:G^O^ for an 8-oxo-G-inserted). The schematic diagrams of the relative localizations of the SV40 replication origin, *supF* gene, and 8-oxo-G or G -inserted position/strand are shown on the right side of figure. (B) Box plot of the *supF*-mutant frequency in transfected cells for G or 8-oxo-G inserted libraries. The *supF* mutant frequency were determined by the conventional *supF* assay, and statistical significance was determined by paired student’s *t* test (n=3, \**P* < 0.05, \*\*\**P* < 0.0005). (C) Number of N_12_-BC sequences with a variant exceeding VF 0.2 according to their nucleotide position (refer to the legend in Figure 2A). The mutations at positions 176 and 177 were excluded. (D) Stack bar graph representing the number of N_12_-BC sequences with mutations from each sample used for further data analysis (refer to the legend in Figure 5B). (E) Box plot of the mutation frequency calculated from NGS data as described in Figure 5C. The asterisks indicate statistical significance determined by paired student’s *t* test (n=3, \**P* < 0.05, \*\*\**P* < 0.005).

### A set of pNGS2 libraries revealed that a single 8-oxo-G induces strand-biased mutations exclusively from G:C but not from C:G sites

The comparison of the types of mutations and the nucleotide bases of SNSs are shown in Figures 9A and 9B, respectively. The distribution of mutation types was not significantly altered between the G and 8-oxo-G inserted libraries, with SNSs being the dominant type of mutations observed (Figure 9A). On the other hand, the SNS patterns significantly varied depending on the library and the DNA-strand in which 8-oxo-G was introduced: while in pNGS2-K1 and -K4 substitutions were predominantly from C, in pNGS2-K2 and -K3 substitutions were from G (Figure 9B). The trinucleotide signatures of SNSs in both groups exhibited a bias towards 5’-T<u>C</u>N-3’:5’-N<u>G</u>A-3’ sites (Figure 9-figure supplement 1); however, this is not conclusive, again due to the strong positional biases as shown in graphs depicting the region (Figure 9C and Figure 9-figure supplements 2 and 3). In pNGS2-K1 and -K4, which have the 8-oxo-G inserted in the top strand of the *supF* gene region, the C to N mutations were uniformly distributed in specific positions throughout the region from 80 to 200. On the other hand, in pNGS2-K2 and -K3, which have the 8-oxo-G inserted in the bottom strand of the *supF* gene region, the G to N mutations were distributed in a wider region but in fewer highly-specific positions, such as 5 and 27. The positions 5 and 27 were previously mentioned as hotspots for spontaneous mutations that were further increased by irradiation. The results clearly demonstrate that the presence of a single 8-oxo-G can induce mutations at sites apart from the original damage-site, and the 5’-T<u>C</u>-3’:5’-<u>G</u>A-3’ site on the same strand emerges as the mutational signature associated with this phenomenon. Also, there is no significant difference between pNGS2-K1 and -K4 versus pNGS2-K2 and -K3, suggesting that the orientation of the replication/transcriptions are not a crucial factor for the strand bias of the mutations. However, the data also indicates that, in addition to the trinucleotide signatures, there is a strong positional bias, since there are 5’-T<u>C</u>N-3’:5’-N<u>G</u>A-3’ sites where mutations cannot be observed, such as positions 24 and 34. The incomplete classifications of the N_12_-BC-variant sequences showed that the insertion of 8-oxo-G led to a slight increase in the proportion of multiple mutations per N_12_-BC (Figure 10A). Nevertheless, in addition to the incomplete classification, it is also difficult to make conclusion due to the relatively low number of mutations; both single and multiple mutations per N_12_-BC show a strand bias, and multiple mutations may even further reinforce this trend (Figure 10B and Figure 10-figure supplement 1).

**Figure 9.**
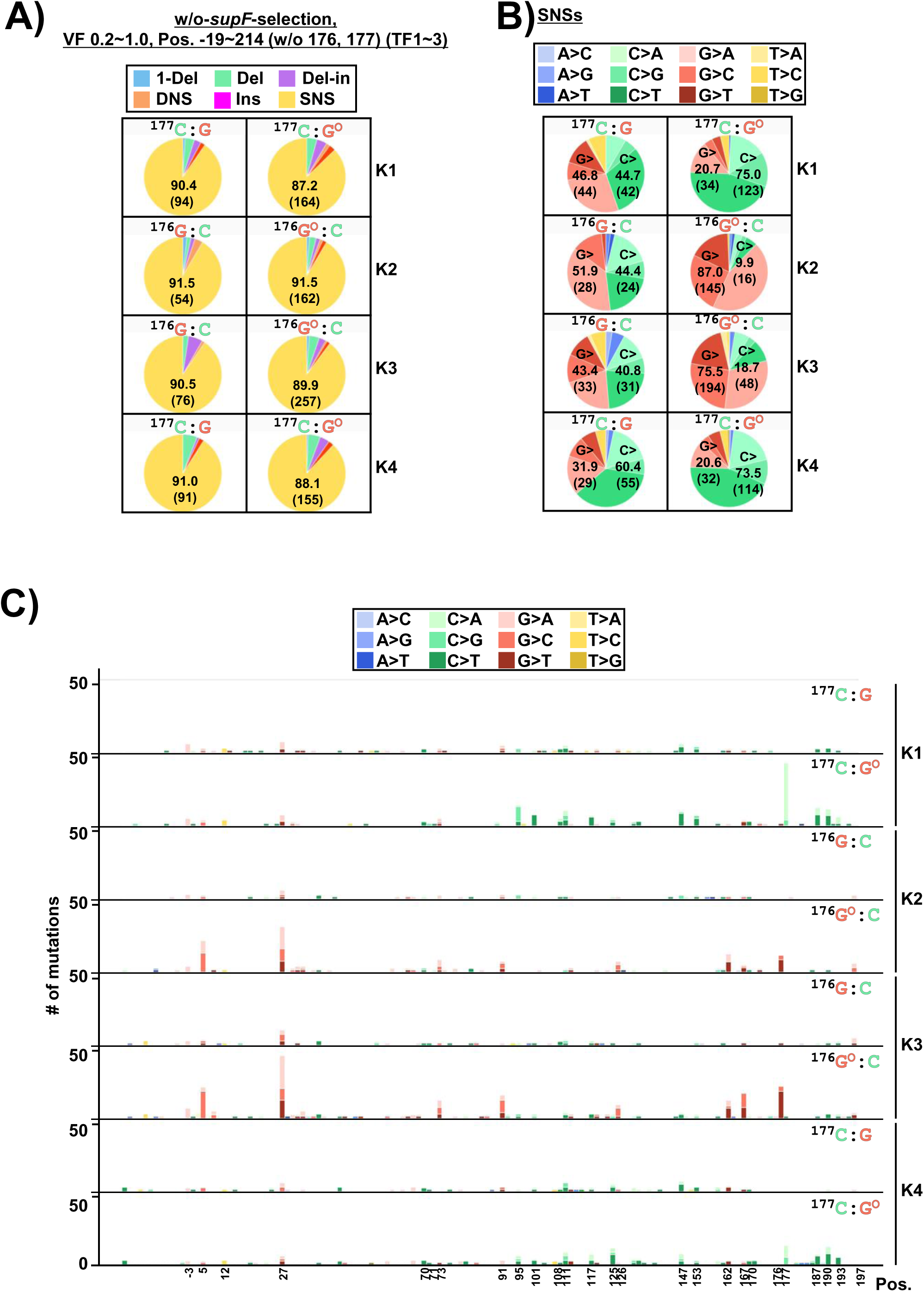
A single 8-oxo-G lesion induces strand-biased single nucleotide substitutions at G:C sites (w/o-*supF*-selection, VF 0.2∼1.0) Analysis of mutations in w/o-*supF*-selection experiments (^176^G:C or ^176^G^O^:C for pNGS2 -K2 and -K3; ^177^C:G or ^177^C:G^O^ for pNGS2 -K1 and -K4). (A) Pie charts of the proportions of different mutation types for pNGS2-K1∼K4 with an inserted G or 8-oxo-G (refer to the legend in Figure 2C). (B) Pie charts of the proportions of different base substitutions in SNSs (refer to the legend in Figure 2D). (C) Number of SNSs according to their nucleotide position (refer to the legend in Figure 2E).

**Figure 10.**
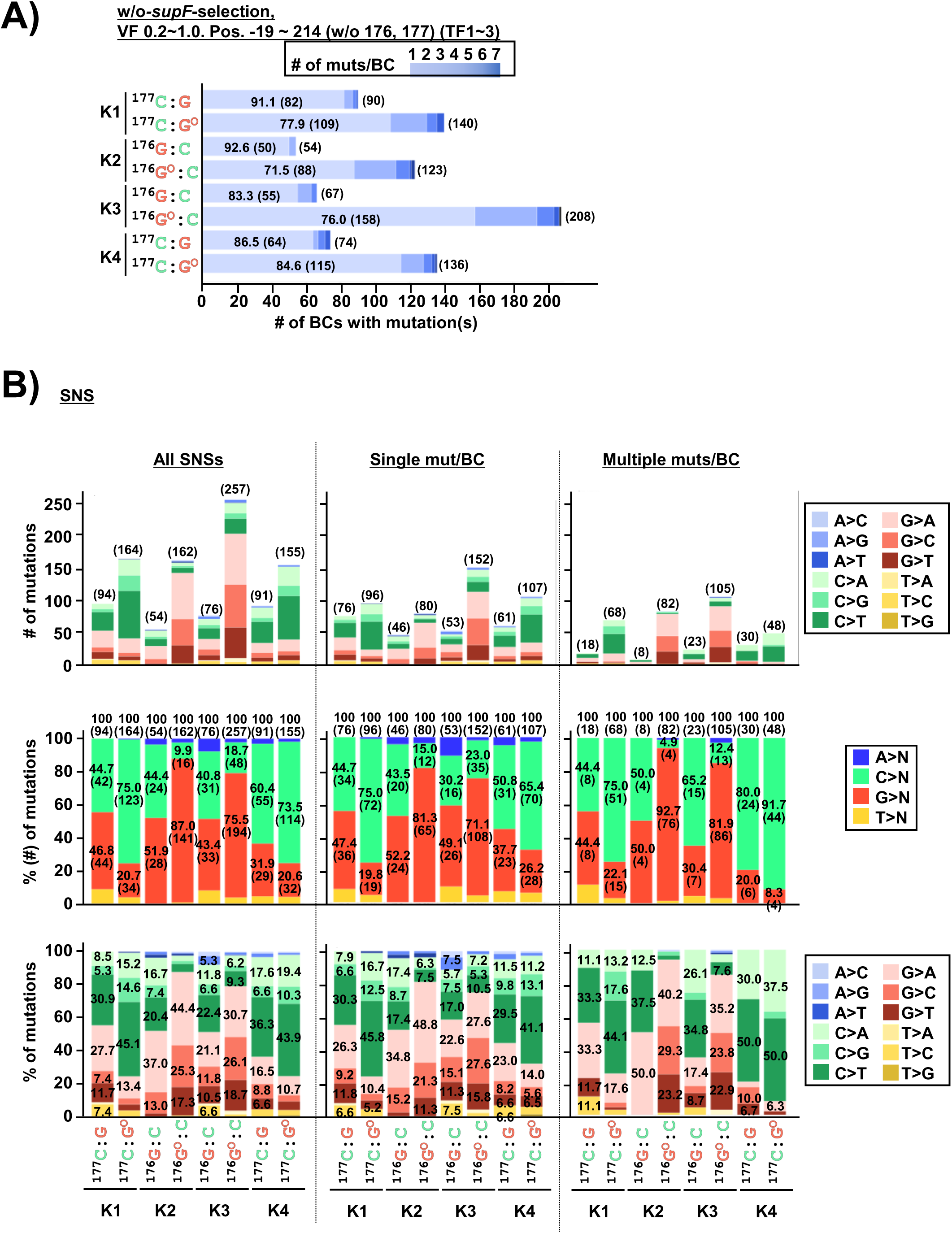
Analysis of 8-oxo-G insertion induced base substitutions in single and multiple *supF* mutations per N_12_-BC (w/o-*supF*-selection, VF 0.2∼1.0). (A) Proportion of N_12_-BC sequences with single (1) or multiple (2∼7) mutations for pNGS2-K1∼K4 with an inserted G or 8-oxo-G. The combined data from three independent transfection-experiments (TF1, TF2, and TF3); mutations at positions 176 and 177 are excluded. The percentage and number of N_12_-BCs with a single mutation are denoted inside each bar, and the total number of N_12_-BCs are indicated outside the bars in the parentheses. (B) Top-row bar graphs represent the number of SNSs in pNGS2-K1∼K4 with an inserted G or 8-oxo-G − combined data from three independent transfection experiments without mutations at positions 176 and 177, for either all mutations (left side), single-mutation per N_12_-BC (middle), or multiple-mutations per N_12_-BC (right side). Middle-row bar graphs represent the relative proportions of the four bases in SNSs. Bottom-row bar graphs represent a detailed breakdown of SNSs according to base substitution, as indicated in the color legend on the right. Throughout panel (B) the numbers inside or above the bars indicate the percentage of SNSs, and the number of detected SNSs is denoted in parentheses.

### The NGS data from *supF*-selection complement the data from w/o-*supF* selection experiments

The NGS analysis in the w/o-*supF*-selection experiments provides less biased data compared to that of *supF*-selection; however, smaller amounts of mutation spectra data could be obtained. Based on the experiments we conducted with chronic irradiation, we can expect that the combined data from both w/o-*supF*-selection and *supF*-selection would provide valuable information if the bias from the selection is taken into account. Therefore, to delve deeper into the mutational spectra at the sites apart from the introduced G or 8-oxo-G in a series of libraries (referred to as G-library or 8-oxo-G library, respectively), the N_12_-BC libraries from roughly 2,000 RF01 colonies grown on *supF* selection plates were analyzed. The N_12_-BCs-variants, excluding positions 176 and 177, were extracted using VF 0.4. The detected number of N_12_-BCs with variant sequence(s) ranged from 76 to 247, and between 120 and 390 mutations were detected at positions -19 to 214 (Figure 11-figure supplement 1A). In order to ensure a fair comparison between samples, the numbers of N_12_-BCs with mutations were limited to about one hundred, as already explained about the data in Figure 2, and the results from TF1, TF2, and TF3 were combined (Figure 11–figure supplement 1B). The positions and types of mutations were very similar among G-libraries, as well as between pNGS2-K1 and -K4, and pNGS2-K2 and -K3 for 8-oxo-G-libraries (Figure 11A). Here the results exhibited a trend similar to the data from the w/o-*supF*-selection experiments shown in Figure 8C. However, there are a few distinct differences, such as the proportion of deletions or deletion-insertions (Figure 11-figure supplement 1 versus Figure 9A). The differences may be due to the *supF*-selection bias, which is often observed especially at lower mutation frequencies. Although the proportion of base substitutions were different, the proportions of individual SNSs seemed to be very similar between the data of *supF*-selection and w/o-*supF*-selection (Figure 11B versus Figure 9B). The proportions of SNSs showed significant difference depending on the 8-oxo-G-library: while in pNGS2-K1 and -K4 substitutions primarily occurred form C, in pNGS2-K2 and -K3 they occurred from G. As expected, the mutations were predominantly located in the tRNA-cloverleaf and promoter regions of the *supF* gene, which causes a noticeable bias on not only positions but also substituted bases. The G-libraries share more similarities in mutations, suggesting that background mutations can be obtained from *supF*-selection, which could help mitigate experimental bias. As described in our previous report, our novel *supF* NGS assay is at least ten-times higher in efficiency than the conventional *supF* assay, and is able to identify practically the whole range of SNSs in the *supF*-selection. Therefore, the quality of the libraries is essential for accurate analysis in the investigation of mutations induced by a variety of mutagens, environmental factors, or deficiency of genes. As already mentioned, all libraries were synthesized by *in vitro* enzymatic reactions and purified following treatment with T5 exonuclease. It has been observed that the frequencies of endogenous mutations in the *supF* gene are associated with the presence of nicks in shuttle vectors (46,57). However, it is impossible to completely eliminate vectors with poor quality, and this is why, in order to ensure the quality of libraries, it is important to thoroughly detect the mutations in the control libraries and treatments. The G-libraries shared similarities between pNGS2-K1 and -K3, with more mutations at positions -3, 5, and 12 compared to the pNGS2-K2 and -K4 libraries (Figure 11C and Figure 11-figure supplement 3). These mutations may have occurred via a template switching process (Figure 4-figure supplement 1B), and there is a possibility that the distance from the SV40 replication origin or the opposite replication strand may have contributed to their frequency. In contrast to G-libraries, 8-oxo-G-libraries exhibited similar patterns for mutations in pNGS2-K1 and -K4, as well as in pNGS2-K2 and -K3. In pNGS2-K1 and -K4 the mutations were detected at the same positions in G-libraries and 8-oxo-G-libraries, but more mutations were observed at C in 8-oxo-G-libraries (Figure 11-figure supplement 2). In pNGS2-K2 and -K3, although the mutations were observed predominantly at G, mutations at positions outside of the *supF* cloverleaf and promoter regions were significantly decreased compared to the data from w/o-*supF*-selection. A strong strand bias was observed in the trinucleotide mutational signatures at 5’-T<u>C</u>N-3’:5’-N<u>G</u>A-3’ sites (Figure 11-figure supplement 3), which is similar to the data from w/o-*supF*-selection upon initial observation (Figure 9-figure supplement 1). In *supF*-selection experiments, the insertion of 8-oxo-G led to a slight increase in the proportion of multiple mutations per N_12_-BC (Figure 11-figure supplement 4A), that is consistent with the data from w/o-*supF*-selection shown in Figure 10A. When the mutations detected by *supF*-selection were classified as single or multiple mutations per N_12_-BC (Figure 11-figure supplement 4B), differences in the mutational spectra were revealed between single and multiple mutations per N_12_-BC, and the multiple mutations were very similar to the case of w/o-*supF*-selection. The same trends were shared with the data from the experiments with gamma-irradiation using pNGS2-K2, shown in Figure 3. These data indicate that single mutations per N_12_-BC may exhibit strong *supF*-selection bias due to the reliance on the loss of function of the *supF* gene as an amber suppressor tRNA.

**Figure 11.**
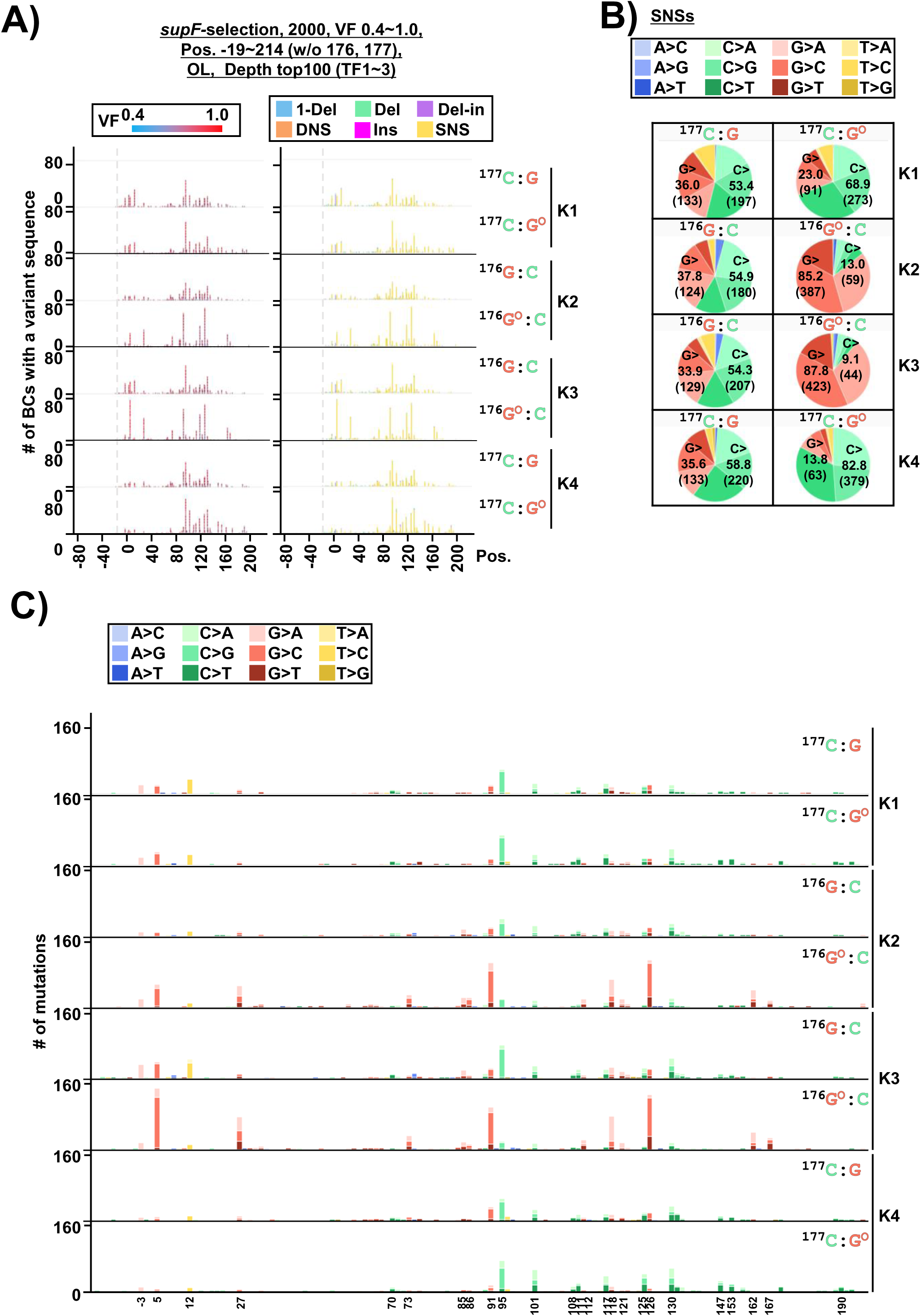
Analysis by *supF* NGS assay of the mutations induced by the insertion of an 8-oxo-G (*supF*-selection, VF 0.4∼1.0). (A) Number of N_12_-BC sequences with a variant exceeding VF 0.4 according to their nucleotide position (Pos.) – combined data from three independent transfection experiments (TF1, TF2, and TF3) for pNGS2-K1∼K4 with an inserted G or 8-oxo-G (refer to the legend of Figure 2A). (B) Pie charts depicting individual base substitutions (in different colors, in the legend on top) as proportions of SNSs. The percentage (and number, in parentheses) of total substitutions from guanine (G>) or cytosine (C>) are indicated in the pie charts. (C) Number of SNSs according to their nucleotide position (Pos.) – combined data from three transfection experiments. The individual base substitutions are shown in different colors as indicated in the legend on top.

The comparison between the positions and spectra of all mutations from *supF*-selection and w/o-*supF*-selection is shown in Figure 12 (Figure 9-figure supplements 2 and 3, and Figure 11-figure supplements 5-10). Relatively non-biased data on the mutations induced by 8-oxo-G were able to be obtained from the combination of w/o-*supF*-selection and multiple mutations per N_12_-BC from *supF*-selection. The following is a summary of the observations: SNSs almost exclusively occurred at 5’-T<u>C</u>-3’:5’-<u>G</u>A-3’ sites throughout the analyzed region in all libraries; The mutations at 5’-T<u>C</u>-3’:5’-<u>G</u>A-3’ sites were generated spontaneously in all G-libraries without strand bias; The insertion of a single 8-oxo-G introduced strongly strand-biased mutations on the same strand at 5’-<u>G</u>A-3’:5’-T<u>C</u>-3’ sites. Furthermore, there were also apparent positional biases even without *supF*-selection. As mentioned above, positions between -8 and 17 represent a distinct and exclusive region for template switching which produced particular mutations: SNSs at -3 (G>A), 5 (G>C), 12 (T>C), and Del-Ins at 3-7 (<u>T</u>T<u>G</u>A<u>T</u>><u>A</u>T<u>C</u>A<u>A</u>); these base substitutions were almost invariably biased and predetermined by fixed bases for a template strand in a quasi-palindromic sequence. Although the underlying molecular mechanisms of template switching are still unclear, it was hypothesized that the 8-oxo-G incidentally caused the replication fork to stall or collapse at a position separate from the 8-oxo-G, followed by disengagement of the lagging strand and invasion of an adjacent replication fork through the annealing to partial homologies in the other strand. Except for this region, the substituted bases in SNSs at 5’-T<u>C</u>-3’:5’-<u>G</u>A-3’ sequences were not predetermined by fixed bases (Figure 12), suggesting these SNSs may have occurred via abasic sites at 5’-T<u>C</u>-3’:5’-<u>G</u>A-3’ sites, which can be assumed with a high probability to have emerged from the uracil generated by APOBEC3 cytosine deaminase targeting the cytosine at 5’-T<u>C</u>-3’ in single-stranded DNA (51). There exists a position-dependent bias for the substituted bases in all the libraries due to unknown reasons. Among the possible factors involved are the efficiency of removal of uracil, the different translesion DNA synthesis (TLS) polymerases, etc. In contrast, it is clear that mutations were never detected at positions -1, 10, 24, and 34 (coldspots), even where the sequences at those positions were 5’-T<u>C</u>-3’:5’-<u>G</u>A-3’, and significantly more were detected at positions 5 and 27 (hotspots), especially in the libraries with an inserted 8-oxo-G.

**Figure 12.**
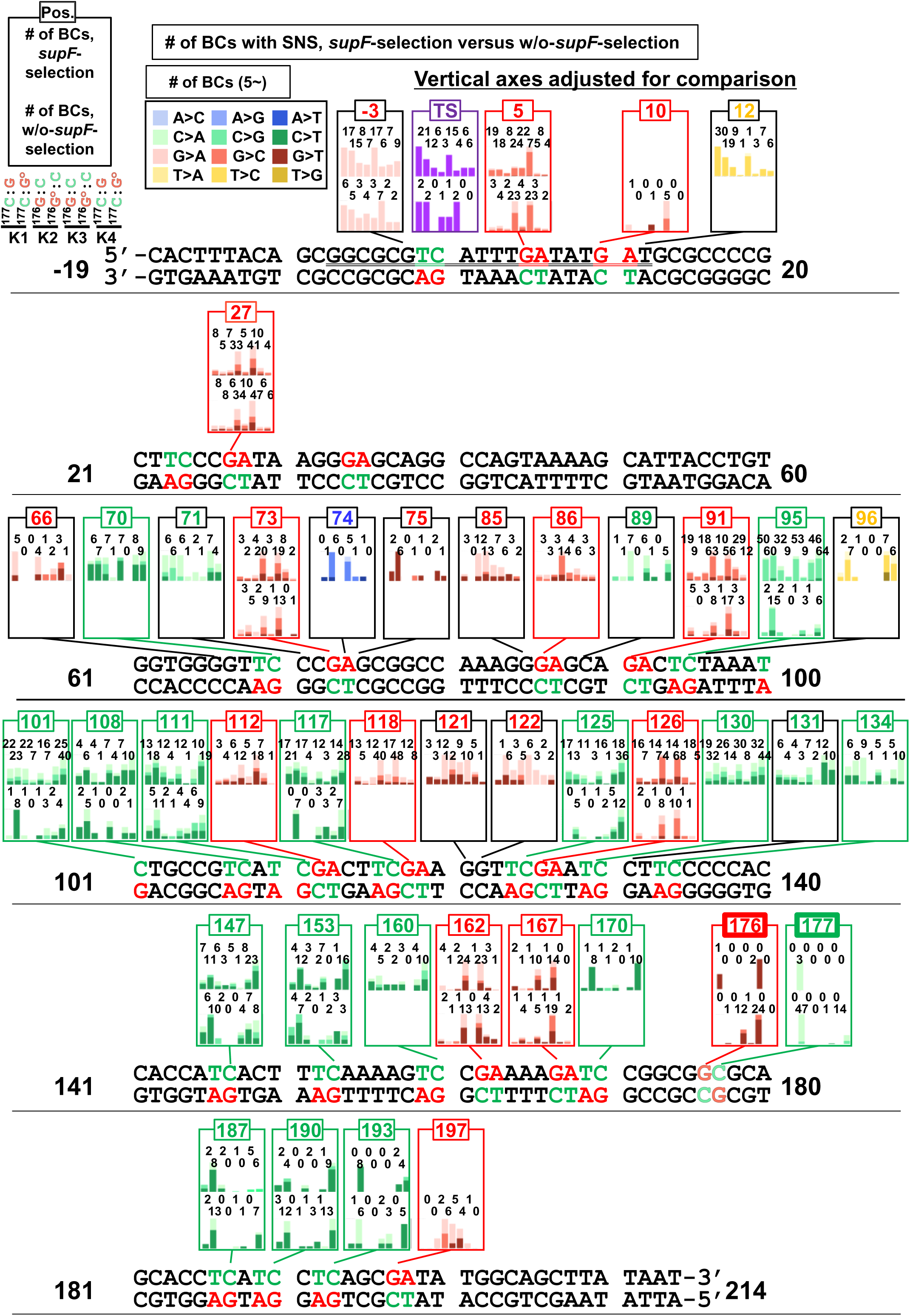
Analysis of SNSs by proportions of substituted bases and mutational positions for pNGS2 libraries -K1 ∼ -K4 with an inserted G or 8-oxo-G (*supF*-selection versus w/o-*supF*-selection) Bar graphs depicting the number of SNSs at the indicated positions on the analyzed sequence, (positions from -19 to 214) − combined data from three transfection experiments (TF1, TF2, and TF3). Each frame depicts the number of SNSs for *supF*-selection (bars on top) and w/o-*supF*-selection (bottom bars). The colors of the bars and segments of stack bars reflect the type of SNSs as indicated in the legend on the top-left side of the figure. The 5’-TC-3’ and 5’-GA-3’ sites in and their positions are shown in green and red, respectively.

### The secondary structure of single-stranded DNA is a crucial factor determining the positions of mutations and possibly the substitutions

In cells, single-stranded DNA is formed during DNA replication, recombination, and transcription. Single-stranded DNA is promptly coated by Replication Protein A (RPA) complexes, which serves as protection to prevent the induction of cluster mutations found in various cancer genomes (1,58,59). Based on the results from the NGS assay, it can be inferred that both template switching and mutational spectra at 5’-T<u>C</u>-3’:5’-<u>G</u>A-3’ sites are important for the mutational signature. It seems highly probable that the single-strand state of DNA may be involved in the mutational process induced by the introduction of 8-oxo-G and irradiation, or even induced spontaneously. Thus, to explore the factors contributing to the strand-biased and position-biased mutations, analysis of the secondary structure of single-stranded DNA was conducted using Mfold web server for the region between positions -20 and 60, which includes six 5’-T<u>C</u>-3’:5’-<u>G</u>A-3 (5’-<u>G</u>A-3’:5’-T<u>C</u>-3) sites with two mutational hotspots (5 and 27) and four coldspots (-1, 10, 24, and 34). The region was divided into overlapping smaller regions of 41 bases by applying a 5-base sliding window (Figure 13). As it was to a certain degree expected, the mutational hotspot positions 5 and 27 were located in the loops of two independent stem-loop structures, regardless of the exact configuration. These stem-loop structures shared the same strand between positions 13 to 26. Also, the above mentioned mutational cold spots, positions -1, 10, 24, and 34, were all located in the stem of the stem-loop structures; position -1 interacted with 10 by G:C base pairing, and the same was observed for 24 and 34. The secondary structures of regions other than -20 to 60 were also predicted by Mfold web server, but these stem-loop structures had a shorter stem, suggesting that they were less fastened, and therefore, it was difficult to find any correlations between the structures and the pattern of mutations (Figure 13-figure supplements 1 and 2).

**Figure 13.**
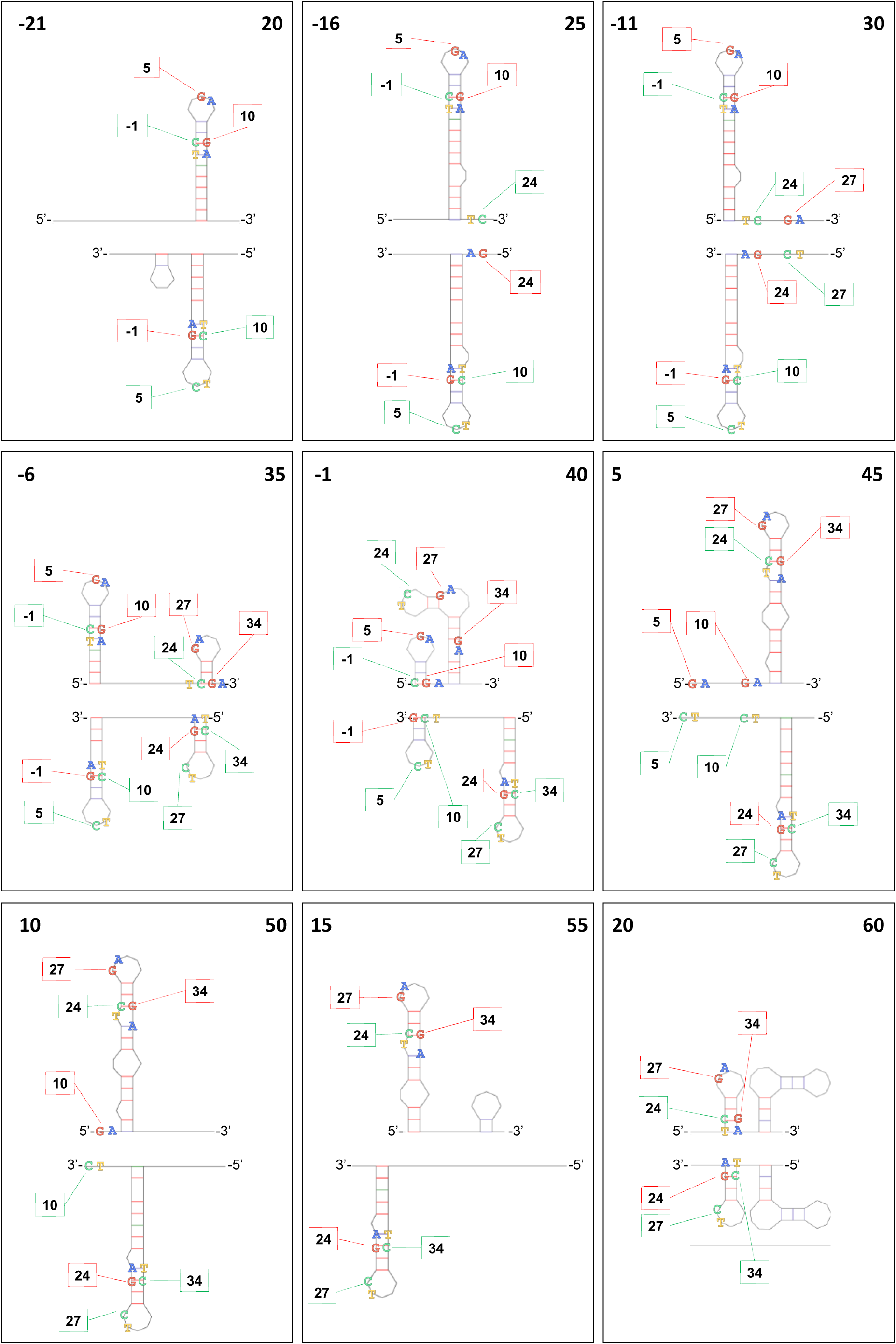
Localization of each 5’-TC-3’:5’-GA-3’ site and the predicted secondary structure of single-stranded DNA. The secondary structure of the single-stranded non-transcribed (top) or transcribed strand (bottom) from position -21 to position 175 of the *supF*-analyzed sequence were predicted by Mfold web server. The analyzed sequence was divided into 27 overlapping regions of 41-bases with a sliding window of 5-bases; the first and last positions of the regions are shown in the top corners of each frame (Fig. 13-figure supplements 1 and 2). The 5’-TC-3’ sites are shown in a green-outline box, and the 5’-GA-3’ sites are shown in a red-outline box for each strand.

### The presence of a single 8-oxo-G lesion increases the frequency of strand-biased multiple mutations at 5’-T<u>C</u>-3’:5’-<u>G</u>A-3’ sites

Finally, to answer the question whether 8-oxo-G influences the induction of mutations by chronic gamma-irradiation exposure, the same set of experiments conducted on pNGS-K2 were carried out using pNGS2-K3 and -K4 libraries with an inserted non-damaged G or an 8-oxo-G. A series of libraries was introduced into cells, and the cells were maintained for 2 days under non-irradiated conditions or chronic gamma-irradiation at a dose-rate of 2 Gy per day. The combined data from two independently prepared libraries and three independent transfection experiments are presented in a set of figures analogous to those shown thus far. The mutations from the w/o *supF*-selection experiments in both libraries pNGS2-K3 and -K4 were increased in response to chronic irradiation as in pNGS2-K2 (Figures 5A and 5B), and interestingly, more mutations were observed in the pNGS2-K3 and -K4 libraries with inserted 8-oxo-G (Figures 14A and 14B). The proportions of mutation types were not significantly altered by irradiation, and SNSs represent the majority of mutations in both libraries (Figure 14-figure supplement 1A); also, the strand bias for the G:C mutations induced by 8-oxo-G was slightly intensified by irradiation (Figure 14-figure supplement 1B). The SNSs seemed to be increased at exactly the same positions at which they were induced by only insertion of an 8-oxo-G. In pNGS2-K3, SNSs were increased at positions 5, 27, 91, 162, 167, and elsewhere at 5’-<u>G</u>A-3’:5’-T<u>C</u>-3’ sites, while in pNGS2-K4 they were increased at positions 95, 111, 187, or 190, and other 5’-T<u>C</u>-3’:5’-<u>G</u>A-3’ sites (Figure 14-figure supplements 1C and 2-6). The increase of both the mutation frequencies obtained from the NGS assay (Figure 14C) and the absolute number of mutations (Figure 14-figure supplement 7A), excluding the mutations at positions 176 and 177, were synergistically greater than the additive effects of 8-oxo-G insertion and irradiation. As expected from the results in figures 6D and 10B, the proportions of SNSs were significantly changed by the insertion of an 8-oxo-G, while a similar effect of radiation could not be clearly inferred for either single or multiple mutations per N_12_-BC (Figure 14-figure supplement 7B). Regarding the 8-oxo-G insertion position 176 for pNGS2-K3 or 177 for pNGS2-K4, the rates of insertion of an A base instead of C were also slightly increased by irradiation for both 8-oxo-G-inserted libraries (Figure 14D). This may suggest that irradiation somehow increases the frequency of incorporation of A by replicative DNA polymerases, or interferes with DNA repair pathways like OGG1-initiated BER for 8-oxo-G:C or MutYH-initiated BER for 8-oxoG:A, where the mechanisms involve many processes, such as damaged base excision or gap filling. The 8-oxo-G:C to T:A mutations were primarily single mutation per N_12_-BC, suggesting that increased frequency of insertion of A is not directly required for the additional mutations (Figure 14-figure supplement 8). Although it has been reported that the mismatch repair pathway and/or OGG1 may be involved in the induction of mutations at positions apart from the damaged base, further studies are necessary to uncover the detailed mechanisms (50,60).

**Figure 14.**
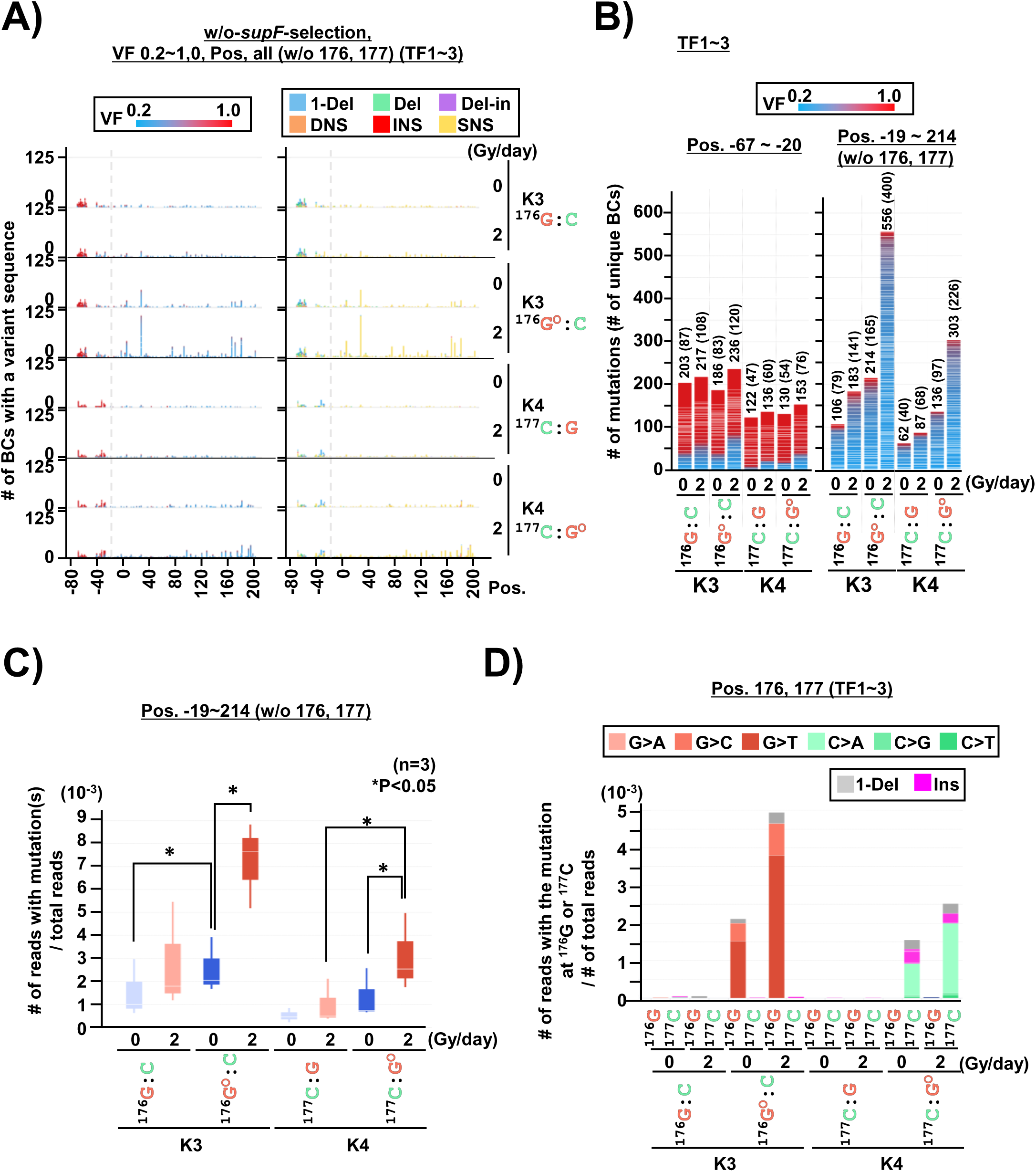
Effects of chronic gamma-irradiation on 8-oxo-G induced action-at-a-distance mutations in cells (pNGS2-K3 and -K4 with an inserted non-damaged G or 8-oxo-G, dose-rate of irradiation 0 or 2 Gy/day for 2 days, w/o-*supF*-selection, VF 0.2∼1.0) (A) Number of N_12_-BC sequences with a variant exceeding VF 0.2 according to their nucleotide position (refer to the legend in Figure 2A). The mutations at positions 176 and 177 were excluded. (B) Stack bar graph representing the number of N_12_-BC sequences with mutations in each sample used for further data analysis (refer to the legend in Figure 5B). The data is separated into two graphs: on the left side positions from -67 to -20, and on the right side – positions from -19 to 214 containing the *supF* gene. (C) Box plot of the mutation frequency calculated from NGS data as described in Figure 5C. The asterisks indicate statistical significance determined by paired student’s *t* test (n=3, \**P* < 0.05). (D) Mutation frequencies for positions 176 and 177 with inserted G or 8-oxo-G − combined data from three transfection experiments. The different base-substitutions or types of mutations are shown in different colors as indicated on in the legend.

### The bias towards hotspot mutations at 5’-T<u>C</u>-3’:5’-<u>G</u>A-3’ sites related to the strand of insertion of 8-oxo-G and irradiation is a result from the process of *supF*-selection

The subtle mutation spectra obtained from *supF*-selection experiments can provide complementary data sets to help us understand the data from the w/o-*supF*-selection experiments. Following the previous analyses of the *supF*-selection data, the mutational spectra for all samples were adjusted depending on the number of reads per N_12_-BC (Figure 15-figure supplement 1A and 1B). As expected from all data sets so far, the major type of mutations was SNSs, and their proportions were slightly increased by both the insertion of an 8-oxo-G and irradiation (Figure 15-figure supplement 1C). A series of mutational biases associated with the insertion of an 8-oxo-G and irradiation were also observed as predicted: the positional bias specific for the *supF*-selection, causing the mutations to be exclusively detected in the promoter and cloverleaf regions of the *supF* gene (Figure 15A, with reference to Figure 2A and Figure 11A); the strand bias manifested as mutations from C:G in opposite directions following the insertion of an 8-oxo-G, which depends on the damaged base inserted strand, and which is slightly intensified by irradiation (Figure 15B, refers to Figure 11B); the bias for hotspots, which were found at 5’-T<u>C</u>-3’:5’-<u>G</u>A-3’ sites (Figure 15C, with reference to Figures 2F, 11C, and 15-figure supplement 2); the multiple mutations per N_12_-BC were increased by the insertion of an 8-oxo-G and slightly increased by irradiation (Figure 15-figure supplement 3A, with reference to Figures 3A and 11-figure supplement 4A) and the proportions of SNSs in single mutation per N_12_-BC were significantly different from the w/o-*supF*-selection data, due to the hotspot bias and preferential substituted bases (Figure 16-figure supplement 3 versus Figure 15-figure supplement 7, with reference to Figure 3D versus Figure 6D, and Figure 10 versus Figure 11-figure supplement 4B).

**Figure 15.**
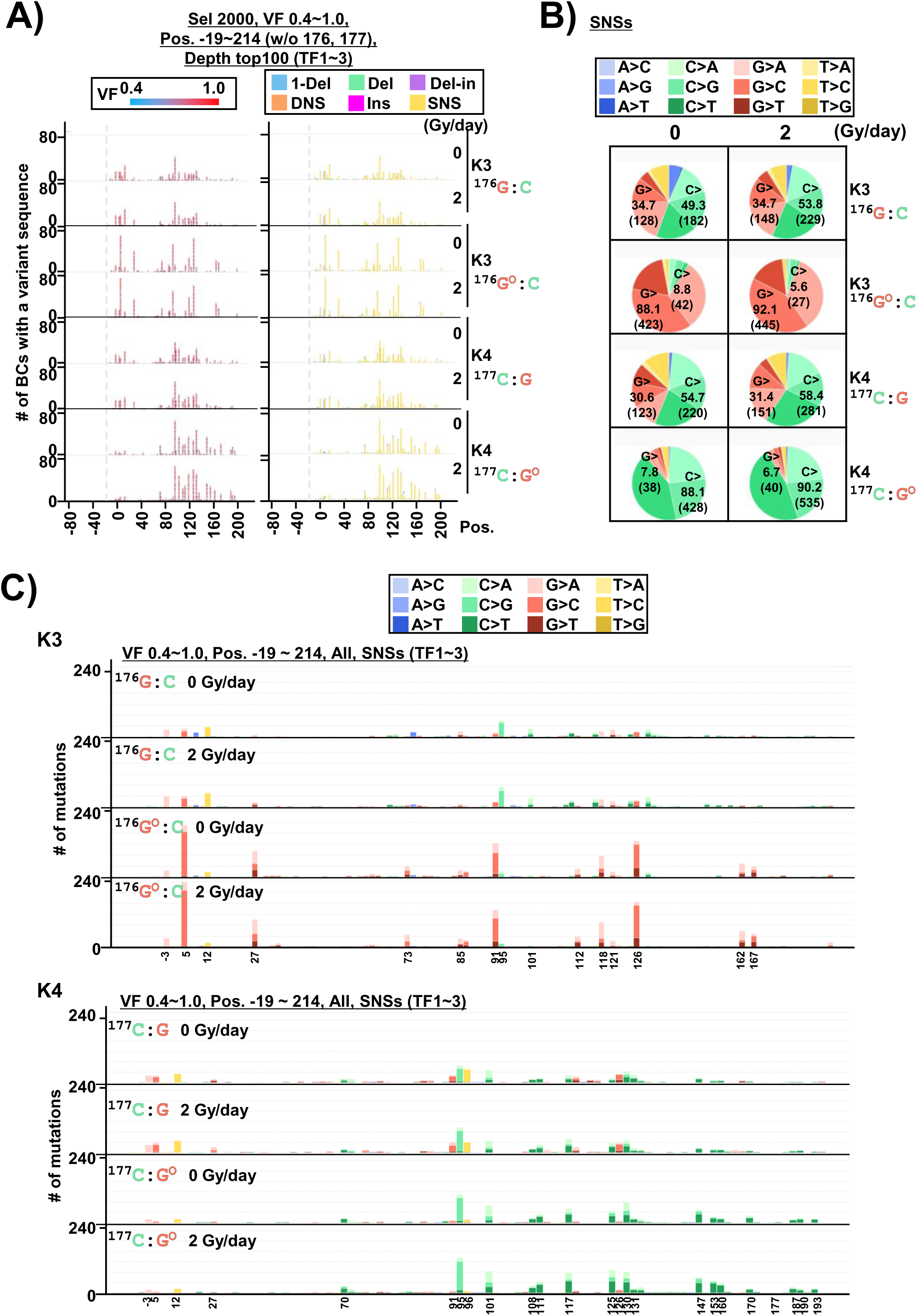
Effects of chronic gamma-irradiation on 8-oxo-G induced action-at-a-distance mutations in cells (pNGS2-K3 and K4 with an inserted non-damaged G or 8-oxo-G, dose-rate of irradiation 0 or 2 Gy/day for 2 days, *supF*-selection, VF 0.4∼1.0) (A) Number of N_12_-BC sequences with a variant exceeding VF 0.4 according to their nucleotide position (refer to the legend in Figure 2A). The mutations at positions 176 and 177 were excluded. (B) Pie charts of the proportions of different base-substitutions in SNSs. The individual base-substitutions are shown in different colors as indicated in the legend on the right side. The percentage (and number, in parentheses) of the total substitutions from guanine (G>) or cytosine (C>) are provided in each pie chart. (C) Number of SNSs according to their nucleotide position (Pos.) – combined data from three transfection experiments. The individual base-substitutions are shown in different colors as indicated on the right side.

**Figure 16.**
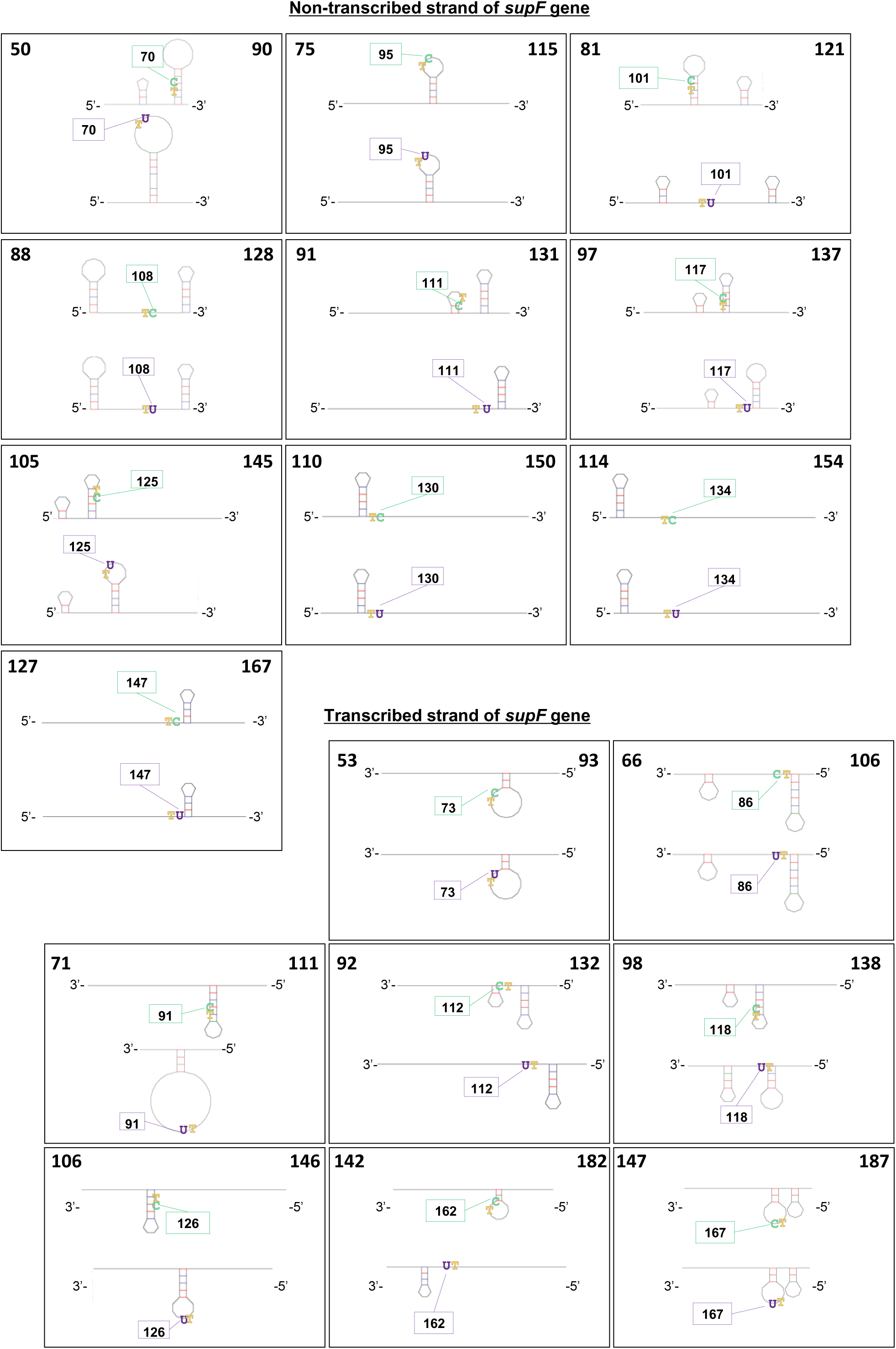
Localization of each 5’-TC-3’:5’-GA-3’ site and the predicted secondary structure of single-stranded DNA. The secondary structure of the single-stranded non-transcribed (top) or transcribed strand (bottom) of the *supF*-analyzed sequence were predicted by Mfold web server. The analyzed sequence was divided into overlapping regions of 41-bases, with the 5’-TC-3’:5’-GA-3’ sites (70, 95, 101, 108, 111, 117. 125, 130, 134, and 147 for the *supF*-non-transcribed strand; 73, 86. 91, 112, 118, 126, 162, and 167 for the *supF*-transcribed strand) located at the center of the sequence. The bottom structure in each frame is the predicted single-stranded DNA secondary structure in the case of the replacement of cytosine with uracil at the site. The first and last positions of the analyzed sequence are shown in the upper left and right corners of each frame. The positions of 5’-T<u>C</u>-3’ sites in the non-transcribed strand are shown in a green-outline box, and the 5’-T<u>C</u>-3’ sites in the transcribed strand are shown in a purple-outline box for each region.

### The hotspot sites for mutations induced by the insertion of 8-oxo-G and irradiation in addition to 5’-T<u>C</u>-3’:5’-<u>G</u>A-3’ sites

Furthermore, abundant mutational spectra from all analyses revealed the common mutation sites that were detected at even higher frequency than some 5’-T<u>C</u>-3’:5’-<u>G</u>A-3’ sites among all the libraries pNGS2-K1 to -K4, at positions 71 (5’-C<u>C</u>C-3’:5’-G<u>G</u>G-3’), 85 (5’-G<u>G</u>G-3’:5’-C<u>C</u>C-3’), and 131 (5’-C<u>C</u>T:A<u>G</u>G-3’), (Figures 7, 12, and 15-figure supplements 4, 5). All these positions were C:G sites and were positioned next to 5’-TC-3’:5’-GA-3’ sites, namely 70 (5’-TC<u>C</u>-3’:5’-<u>G</u>GA-3’), 86 (5’-<u>G</u>GA-3’:5’-TC<u>C</u>-3’), and 130 (5’-TC<u>C</u>-3’:5’-<u>G</u>GA-3’). As one possible mechanism, the C:G to T:A transition due to the deamination of C to U at 5’-T<u>C</u>-3’:5’-<u>G</u>A-3’ sites may result in the generation of secondary 5’-TT<u>C</u>-3’:5’-<u>G</u>AA-3’ sites for deamination. In fact, at all these positions the double nucleotide substitutions (DNSs), CC to TT/TA/AT/TG, were also detected with similar frequencies to the single mutations at the secondary sites. However, since the single mutations at the secondary sites were detected independently from the primary mutations, it is more likely that the deamination occurred at those two sites simultaneously rather than sequentially; another possibility is polymerase errors by TLS polymerases following the primary uracil or abasic sites, although this is unlikely because the sites of mutations were only C:G.

In addition to the series of mutational biases described so far, positional biases for the substituted bases at both 5’-T<u>C</u>-3’:5’-<u>G</u>A-3’ and 5’-<u>G</u>A-3’:5’-T<u>C</u>-3’ sites, although difficult to discern, were certainly present. The C:G to T:A transitions were the major SNSs detected at the sites in w/o-*supF*-selection and in multiple mutations from *supF*-selection experiments (Figures 3D, 6D, 10B, 11-figure supplement 4B, and 14-figure supplement 7B). The C to T transitions at 5’-TCN-3’ are associated with COSMIC SBS2 and are a type of mutation observed in various cancers. These mutations are thought to arise from during DNA replication from uracil residues generated by the deamination of cytosine by AID/APOBEC enzymes (61,62). There seem to be some positional biases affecting their proportions, but it proved difficult to make conclusions from the limited number of mutations at each position. In addition to the transition mutations, the C:G to G:C and A:T transversions, which are associated with COSMIC SBS13, were observed at the same sites and are believed to arise from the glycosylation of uracil, leading to abasic sites, and subsequent error-prone replication by TLS polymerases (63–65). The positional bias affecting the proportions of C:G to G:C and C:G to A:T transversions could also be observed (Figures 5F, 9C, 14-figure supplement 1B). The C:G to G:C transversions were predominantly observed at positions 95 and 125 for 5’-T<u>C</u>-3’:5’-<u>G</u>A-3’ sites, and 5, 91, 126, and 167 for 5’-<u>G</u>A-3’:5’-T<u>C</u>-3’ sites and were affected by both the insertion of 8-oxo-G and irradiation (Figure 14-figure supplements 3 and 5). Notably, the proportions of the mutations were not affected by any sequences other than the 5’-T<u>C</u>-3’:5’-<u>G</u>A-3’ sequence, prompting us to consider different factors that might be discerned from the data. We entertained the possibility that one such factor might be the secondary DNA structure, and therefore, the secondary structure predictions for single-stranded DNA were further explored. During replication, repair, and recombination, RPA is required for shielding exposed single-stranded DNA in order to prevent the formation of secondary structures, which plays a crucial role in maintaining genome stability (66). The interaction between RPA and single-stranded DNA needs to be highly dynamic, which aids in destabilizing secondary structures and hairpin structures of single-stranded DNA (67,68). This process creates small single-stranded regions that are essential for optimal DNA synthesis and repair. The structure of these single-stranded DNA regions is expected to exhibit high dynamics, and alterations are likely to occur around positions that have undergone deamination. Therefore, we conducted further predictions of the secondary structures of single-stranded DNA, as described in Figure 13, but the sequence at 5’-T<u>C</u>-3’ sites was changed to T/U instead of C with a focus on the 5’-T<u>C</u>-3’:5’-<u>G</u>A-3’ sequence. Position 125 at 5’-T<u>C</u>-3’:5’-<u>G</u>A-3’ sites, and positions 126 and 167 at 5’-<u>G</u>A-3’:5’-T<u>C</u>-3’ sites, were located in the loop of the newly formed stem-loop structures (Figure 16). The underlying mechanisms are not known but it is possible that the stem-loop structure, which may contain a C, U or an abasic site either in single-stranded DNA or in the adjacent base-paired helical stem-structure, makes a difference for the DNA replication mechanisms of abasic sites. Abasic sites are endogenous DNA lesions that are highly frequent in cells, and their repair pathways and TLS are crucial for cells to maintain genomic stability and cellular survival (65). Recently, there has been a significant focus on understanding the difference between the abasic sites that occur in double-stranded DNA and those that occur in single-stranded DNA. Rev1 (REV1 DNA Directed Polymerase) and Pol ζ (DNA Polymerase zeta) are involved in TLS of abasic sites by filling in dCMP, resulting in C:G to G:C transversion mutations during DNA replication (69–71). Pol η (DNA polymerase eta), another TLS polymerase, is known to contribute to both C:G to G:C and A:T transversions when filling in abasic sites (72–74). In addition, PrimPol (Primase-Polymerase) has been reported to prevent the transversion mutations at abasic sites (75,76). Furthermore, HMCES (5-hydroxymethylcytosine binding, ES cell specific) protein interacts with abasic sites in single-stranded DNA exposed at DNA replication forks and blocks TLS (77–79).

Exposure to ROS is one of the major sources for endogenous DNA-lesions in genomic DNA, and 8-oxo-G is known as one of the most frequently produced oxidative base-damages. We have previously reported that chronic gamma-irradiation at a dose-rate of 1 Gy per day introduced cellular senescence and inflammatory reactions to the normal human fibroblast cells more efficiently than acute gamma-irradiation at a dose-rate of 1 Gy per minute, and produce ROS via mitochondrial damage-response (47,48). In this study, a set of *supF*-NGS assays were carried out for the mutations induced by chronic gamma-irradiation and an 8-oxo-G, and an integrative perspective from the data of *supF*-selection and w/o-*supF*-selection provided much less biased mutation spectra comparing to an accumulated number of mutagenesis studies using various types of selective marker gene. As a consequence, the chronic gamma-irradiation induced base-substitutions from C:G at 5’-T<u>C</u>-3’:5’-<u>G</u>A-3’ sites, which were supposed to be increased at the same sites for spontaneous mutations rather than at the DNA-lesions directly induced by the radiation. A single 8-oxo-G produced by *in vitro* enzymatic reactions significantly increased the same mutations at the 5’-T<u>C</u>-3’:5’-<u>G</u>A-3’ sites apart from the introduced 8-oxo-G, and the mutations were highly strand-biased depending on a strand introduced with an 8-oxo-G. And, the strand-biased mutations were significantly further increased by chronic gamma-irradiation via the mechanism that remains undetermined yet. It is possible that it is because of the induction of APOBECs-expressions by exposure to the radiation and DNA-damages (79,80). It has been reported that the analysis for mutational signatures implicated APOBEC-mutagenesis following radiotherapy for patients with glioma (81). The frequencies and the substituted bases of mutations at each position, especially 5’-T<u>C</u>-3’ sites, are strongly associated with the predicted secondary structure of single-stranded DNA. The underlying mechanisms for the achievement of the mutations are intricate and even stochastic, which is dependent on the balance of a number of physiological pathways influenced by many intra-and extracellular factors. The NGS analysis and large public data resources such as The Cancer Genome Atlas (TCGA) (82) and the Genomic Data Commons (GDC) (83), will keep providing results that are factually certain. However, in order to fully utilize the available data, the underlying individual and combined mechanisms must be addressed and confirmed by reproducible methods in an ordinary research laboratory setting without specialized equipment or techniques. The high-throughput *supF*-NGS assay provided in this study, that is developed as a more versatile and simple method than that presented in the previous study, can provide more prospect for various designed experiments in any laboratories. Especially, the data from w/o-*supF*-selection could provide less-biased mutational spectra, and the results indicate any nucleotide sequences, even a non-coding gene or artificially designed sequences, can be used for this assay. Therefore, it is promising that the further development of the sequence for the analysis or applying to long read sequencing, such as PacBio Revio system, will provide us more informative data. Accumulating data from future research will reveal the intricacies of the mutagenesis, and will hopefully contribute to machine learning for the discovering the patterns for the mutations induced by the complexed mechanisms, or even to improve base editing techniques for the treating genetic diseases.

## Materials and methods

### Construction of pNGS2-N_12_-BC libraries

The vector maps of four series of *supF* shuttle vectors – pNGS2-K1, -K2, -K3, and -K4, and the construction method for the pNGS2-N_12_-libraries were described in detail in our previous paper (46). Briefly, pNGS2 was digested with *Eco*RV-HF restriction enzyme (New England Biolabs, USA) and purified by agarose gel electrophoresis followed by extraction using FastGene Gel/PCR Extraction kit (NIPPON Genetics, Japan). The purified *Eco*RV-HF-digested pNGS2 was combined with single-stranded randomized N_12_-oligonucleotides (5’-dGGCCTCAGCGAATTGCAAGCTTCTAGAAGGCGATNNNNNNNNNNNNATCGAATTCGGATCCTTTCTCAACGTAA-3’, where N = A, T, G or C) and assembled using NEBuilder HiFi DNA Assembly Master Mix (New England Biolabs). The randomized N_12_-oligonucleotides were commercially synthesized and purified on a reversed-phase column (FASMAC, Japan). The *E. coli* JM109 electro-competent cells were transformed with either the assembly reaction mixture or non-assembly control mixture by electroporation. The transformants were serially diluted, seeded on LB agar plates, and incubated at 37°C overnight. The number of colonies on each plate was counted, and a higher than 99% efficiency of the assembly reaction was confirmed. Approximately 10^4^ colonies were harvested from the plates by scraping. The pNGS2-N_12_-BC libraries were prepared by purification from the harvested cells by using either GenElute Plasmid Miniprep (Merck, Germany) or NucleoBond Xtra Midi (TaKaRa Bio Inc., Japan). Each prepared library was interrogated for the presence of potential *supF* mutants by using the indicator non-SOS induced *E. coli* RF01 strain (49), and a mutation frequency of less than 10^-4^ was considered mutant free.

### Construction of pNGS2-N_12_-BC libraries containing either an 8-Oxo-7,8-dihydroguanine (8-oxo-G) or lesion-free (G)

The procedures for construction of pNGS2 libraries containing either an 8-oxo-7,8-dihydroguanine (8-oxo-G) or lesion-free (G) were described in detail previously (56). The 5’-phosphorylated oligonucleotides carrying 8-oxo-G or G (5’-P-GAGGTGCTGCG*CCGCCGGATC-3’ for pNGS2-K1 and -K4, and 5’-P-GATCCGGCGG*CGCAGCACCTC-3’ for pNGS2-K2 and -K3; P=phosphate, G*=8-oxo-G or lesion-free G) were synthesized and purified by two sequential independent purifications (Hokkaido System Sciences, Japan). Briefly, *E. coli* HB101 bearing VCSM13ΔPS (P_BAD_-pII) were transformed with each individual pNGS2-N_12_-BC library and seeded on LB agar plates. At least 10^4^ colonies were harvested in LB medium by scraping, and the optical density of the cell suspensions was measured at 610 nm (OD_610_). The cells were collected by centrifugation, resuspended in 15% glycerol/LB medium (adjusted to OD_610_ = 50-250), and stored at -80 °C. For culturing, a batch from the cryopreserved stock was added to 2×=YT medium and incubated at 37=°C with shaking until OD_610_ = 0.5. Arabinose at a final concentration of 0.2% was added, and the culture was further incubated overnight (37=°C with shaking). The supernatant containing the phage particles was collected by two consecutive centrifugations. The phage particles were precipitated by adding a one-tenth volume of 20% PEG-6000/2.5=M NaCl solution to the supernatant. The phage particles were then collected by centrifugation and resuspended in 10=mM Tris-HCl (pH=8.0). The suspension was subjected to enzymatic treatment with DNase I and RNase A in order to digest any contaminating bacterial nucleic acids. Then, the circular single-stranded DNA for the pNGS2-BC_12_-libraries was extracted from the phage particles by treatment with proteinase K and SDS, followed by purification using QIAGEN-tip20 (QIAGEN, Venlo, Netherlands). To construct the circular double-stranded DNA for pNGS2-BC_12_-libraries containing either an 8-oxo-G or lesion-free, a five-fold molar excess of oligonucleotides bearing either 8-oxo-G or G was mixed with the phage-extracted circular single-stranded DNA in 1=×=Phusion HF Buffer (New England BioLabs). After denaturation at 90=°C for 2=min followed by rapid cooling, the solutions were heated to 70=°C for 5=min, and then allowed to gradually cool to room temperature. The polymerase and ligase reactions were simultaneously carried out in 1=×=Phusion HF Buffer supplemented with 1=mM dithiothreitol, 0.2=mM dNTPs, and 1=mM NAD^+^, using Phusion High-Fidelity DNA polymerase and Taq DNA ligase (New England BioLabs). The reaction took place at 50=°C for 30=min, and then at 65=°C for 1=h. The sample was then vigorously vortexed to inactivate the enzymes, and after addition of dam methyltransferase and T5 exonuclease was incubated at 37=°C for 3=h to methylate the A in the 5’-GATC-3’ sequence and digest the DNAs other than closed circular double-stranded DNA. The sample was purified with a PureLink PCR Purification Kit (Thermo Fisher Scientific, USA) and precipitated with ethanol. The quality of the circular single-stranded DNA for pNGS2-BC_12_-libraries was checked by gel electrophoresis using agarose gels with GelRed (Biotium, USA).

### Cell culture, transfection, gamma-irradiation, and *supF* mutation analyses

After their construction in the above-outlined manner, pNGS2-N_12_-BC libraries were transfected into the human osteosarcoma U2OS cell line (HTB-96^TM^, American Type Culture Collection, USA) using Lipofectamine 2000 (Thermo Fisher Scientific), as we have previously described (46,49,50). Briefly, trypsinized cells were seeded into either 6-well plates or 35-mm dishes at 3 × 10^5^ cells/well the day before transfection. After culturing for 24=h, 400=ng of pNGS2-BC_12_-library were transfected according to the manufacturer’s instructions. Transfected cells were maintained at 37°C in a humidified, 5% CO_2_ atmosphere for 48 hours. To investigate the effects of chronic gamma-irradiation at different dose-rates (1,000 or 2,000 mGy per 24 hours) on mutations, a specially designed radiation facility was used, where cell culture incubators were placed at different distances from a ^137^Cs-radiation-device (1.11 TBq), as described in our previous paper (47). The dose-rate at each location of sample placement inside the incubators was measured using a GD-302M glass dosimeter (AGC Techno Glass, Japan). At 48 hours after transfection the library was extracted from the cells as previously described (46), To remove unreplicated shuttle vectors, the library was digested with the restriction enzyme *Dpn* I (New England BioLabs), which possesses specificity for methylated 5’-G^m^ATC-3’ sequences. The library was then precipitated by ethanol and dissolved in 15 μl of sterile deionized water (Nacalai, Japan). The *Dpn* I-treated libraries were used for both the conventional *supF* mutagenesis assay and the preparation of NGS samples (w/o-*supF*-selection). For analyzing the *supF* mutant frequency, the *Dpn* I-digested library was transformed into the indicator *E. coli* strain RF01. Serial 5-fold dilutions of transformed RF01 samples were then seeded on titer and selection LB agar plates. Titer plates contained kanamycin and chloramphenicol, while selection plates contained streptomycin and nalidixic acid in addition to kanamycin and chloramphenicol. The *supF* mutant frequencies were estimated based on the number of colonies grown on the individual plates. The presumed number of 2,000 colonies was harvested from selection plates as a cell pellet by scraping, and libraries were extracted using GenElute plasmid miniprep kit for preparation of NGS samples (*supF*-selection).

### Preparation of samples with unique indexes for multiplexed NGS

Each sample with a unique index for multiplexed NGS was prepared by PCR using KOD One PCR Master Mix -Blue-(TOYOBO, Japan). For the standard protocol, either 10 ng of the library extracted from the RF01 colonies grown on selection plates (*supF*-selection) or 1 μl of *Dpn* I-treated library (w/o-*supF*-selection), as described above, was used for a 50 μL PCR reaction (30 cycles of amplification at 98°C for 10 s, 60°C for 5 s, and 68°C for 1 s) with a series of primer sets containing a pre-designed 6-nucleotide sequence (N_6_) as an index sequence for distinguishing each sample by multiplexed NGS data analysis (46). PCR products were purified using NucleoSpin Gel and PCR Clean-up (TaKara Bio Inc.), and then analyzed and quantified by Nanodrop spectrometer (Thermo Fisher Scientific) and agarose gel electrophoresis. The uniquely indexed samples were combined together in the proper proportion according to the expected number of NGS reads for each sample. More than ten reads per N_12_-BC sequence were obtained for all samples in this study (approximately 16 Mb and 400 Mb of sequencing data for the selection plates and *Dpn* I-treated libraries, respectively).

### DNBSeq sequencing and sequencing processing

Preparation of NGS libraries and sequencing were conducted by Seibutsu Giken inc. (Japan), as previously described (46). The DNA nanoballs were prepared by using the MGIEasy Circularization Kit (MGI Tech). The samples were subjected to multiplex{Canet, 2022 #15}ed deep sequencing in 200-bp paired-end mode on the NGS using a BGISEQ-G400 platform (MGI Tech, China).

### NGS data analysis

The process of NGS data analysis was described in our previous paper (46). Briefly, the FASTQ files for each N_6_-indexed data set in an NGS run (sorting was performed by Seibutsu Giken inc.) were used in the following manner: 1) The reverse-complement sequences were obtained using SeqKit tools (85); 2) The paired-end reads were merged using CASPER (86); 3) The merged sequences were mapped to a reference sequence using minimap2 (87); 4) The error corrections were performed based on a whitelist by UMI-tools (88). For the N_12_-BC extraction, only one-base differences between N_12_-BC sequences were allowed, and corrections were implemented using the following regular expression: (".*(GGCGAT){s<=1}(?P<Cell_1>.{12})(?P<UMI_1>.{0})(ATCGAA){s<=1}.*"); 5) Using VarDict, the sequence data for all N_12_-BC sequences with more than 9 reads per N_12_-BC was employed for the alignment to the reference sequence, variant calling and calculation of allele frequencies (89); 7) Spotfire version 7.11.1 (TIBCO, USA) was used for data organization, filtering, and visualization.

## DATA AVAILABILITY

NGS raw data has been uploaded to DDBJ database (http://www.ddbj.nig. ac.jp) under the BioProject accession number PRJDB17510 and BioSample accession number SAMD00737992-SAMD00737998. See Supplementary table S1 for list of nucleotide sequences of primer sets used in multiplex sample preparation for NGS in the respective figures.

## SUPPLEMENTARY DATA

Figure 3 − figure supplements 1-2; Figure 4 − figure supplements 1-9; Figure 5 − figure supplements 1-2; Figure 6 − figure supplements 1-2; Figure 8 − figure supplement S1; Figure 9 − -figure supplement 1-3; Figure 10 − figure supplement 1; Figure 11 − figure supplements 1-10; Figure 13 − figure supplements 1-2; Figure 14 − figure supplements 1-8; Figure 15 − figure supplements 1-12; Supplementary table S1.

## ACKNOWLEDGEMENTS

A part of this study was carried out at the Joint Usage/Research Center Program of the Research Institute for Radiation Biology and Medicine (RIRBM), Hiroshima University.

## FUNDING

This work was supported in part by the Japan Society for the Promotion of Science (JSPS) KAKENHI grant numbers JP 22K12375 (H. Kawai) and 22H03751 (H. Kamiya).

## CONFLICT OF INTEREST

The authors declare no conflict of interest.

## Notes

### Competing Interest Statement

The authors have declared no competing interest.

### Summary of Updates

Abstract is updated; Typo in Reference #65 is corrected.

