## Supplementary material for "Systematic mutagenesis assay promotes comprehension of the strand-bias laws for mutations induced by oxidative DNA damage": Figure 11-figure supplement 3

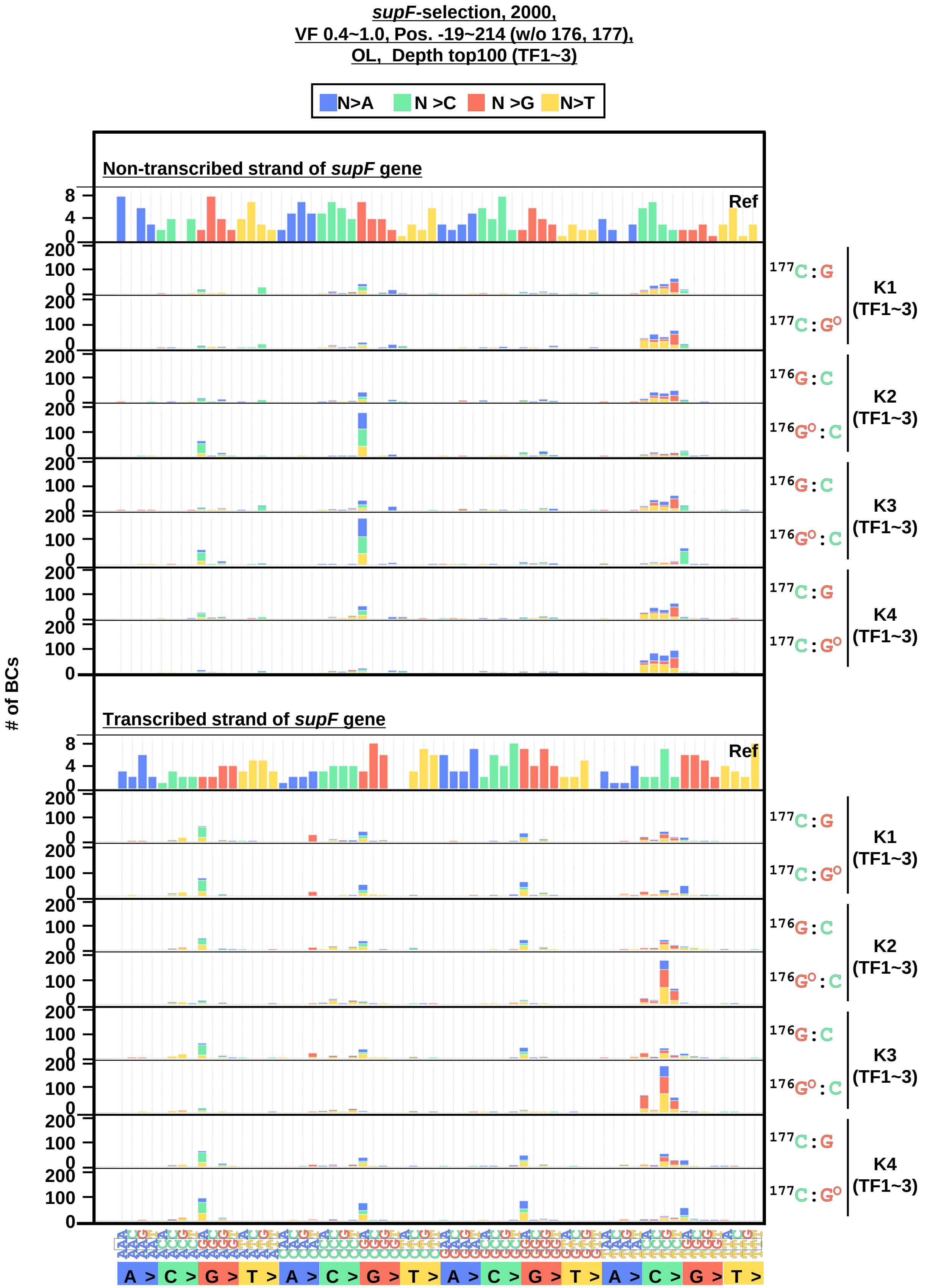


**Fig. 11-figure supplement 3.** The 192 trinucleotide contexts of SNSs (*supF*-selection, VF 0.4~1.0). The number of SNSs in192 trinucleotide contexts for the non-transcribed (top panel) and transcribed (bottom panel) strand of the *supF* gene (nucleotide positions from -19 to 214). The substituted bases are indicated in colors corresponding to the notation on the horizontal axis; the combined data from three transfection experiments (TF1, TF2, and TF3) in pNGS2-K1~K4 with either G or 8-oxo-G were shown on the right side of the graph; the reference sequence for the analysis is denoted as Ref.
