## Supplementary material for "Systematic mutagenesis assay promotes comprehension of the strand-bias laws for mutations induced by oxidative DNA damage": Figure 11-figure supplement 4

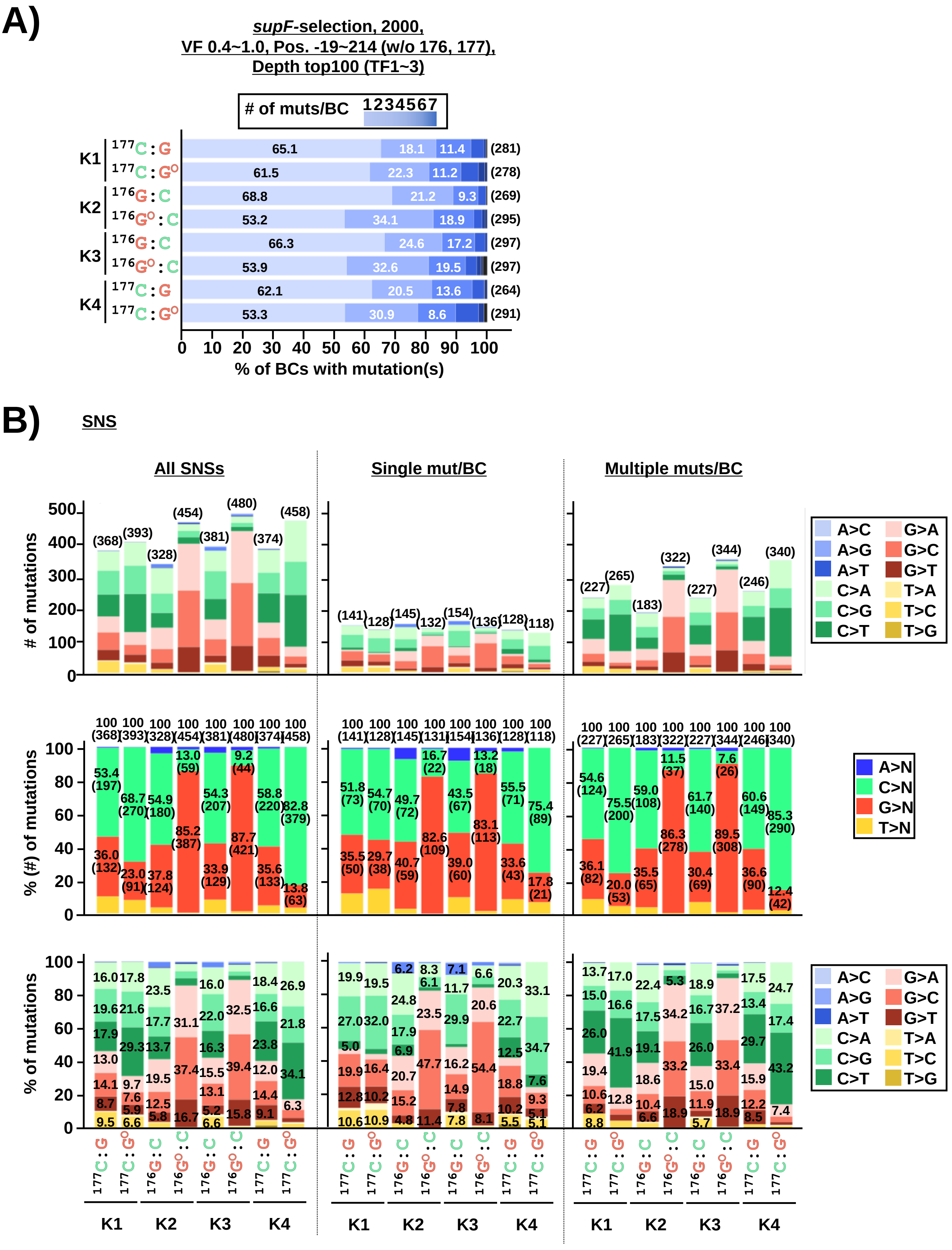


**Fig. 11-figure supplement 4.** Comparing the base-substitutions on the number of *supF* mutations per $N_{12}$-BC induced by the insertion of an 8-oxo-G (*supF*-selection, VF 0.4~1.0). (A) Proportion of N12-BC sequences with single (1) or multiple (2~7) mutations for pNGS2-K1~K4 inserted with either a G or an 8-oxo-G — combined data from three independent transfection experiments (TF1, TF2, and TF3). The percentage of $N_{12}$-BCs with different number of mutations (single, 2, and 3 per $N_{12}$-BC) are indicated inside each bar, and the total number of mutations are presented outside each bar in the parentheses. (B) The layout of the figure is analogous to Figure 10, this time for the data from *supF*-selection.
