## Supplementary material for "Systematic mutagenesis assay promotes comprehension of the strand-bias laws for mutations induced by oxidative DNA damage": Figure 11-figure supplement 8

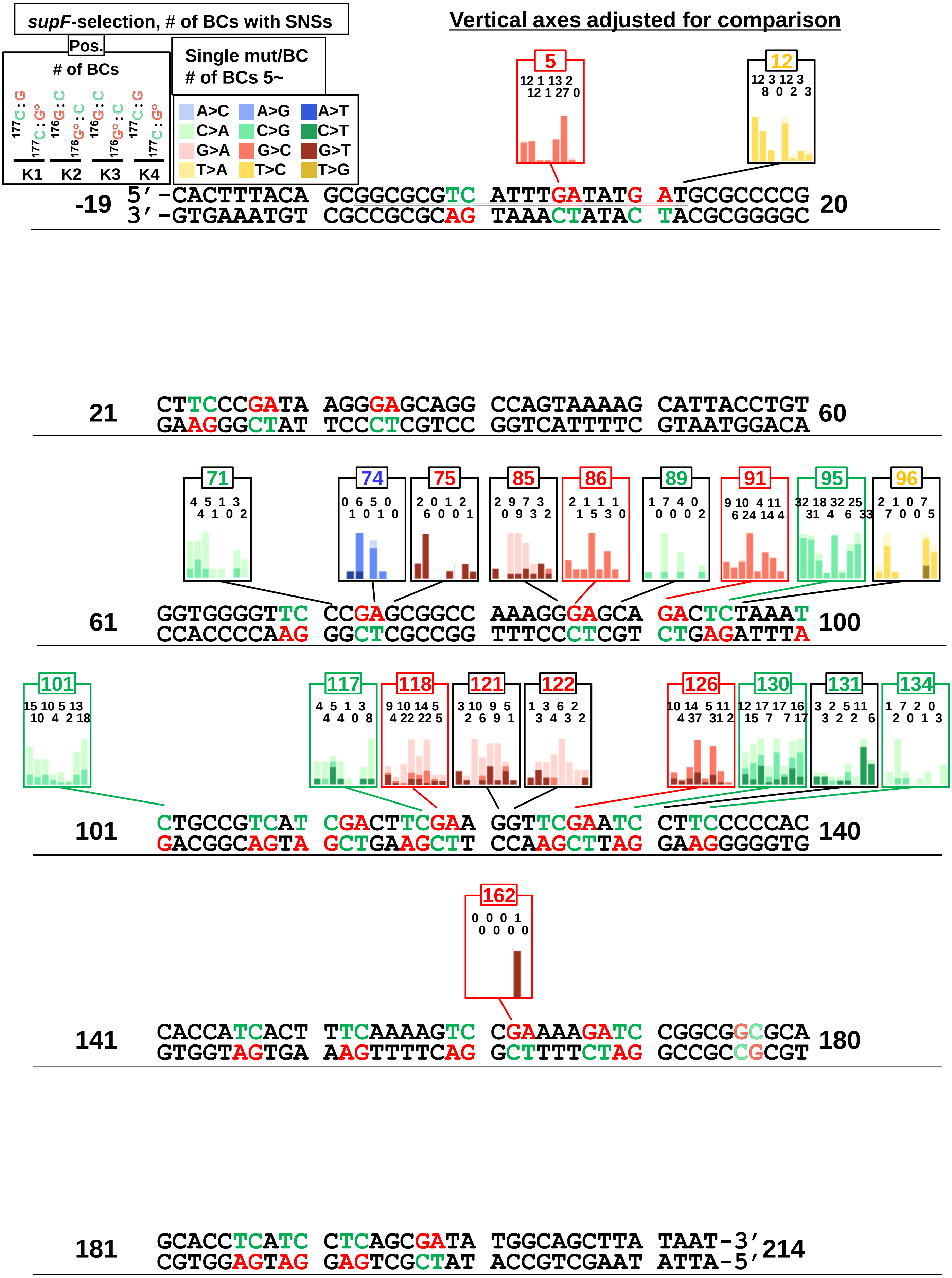


**Fig. 11-figure supplement 8.** Proportion of substituted bases in SNSs for single mutation per $N_{12}$-BC according to their positions and the insertion of an 8-oxo-G (*supF*-selection). Bar graphs provide combined data for the number of SNSs from three transfection experiments (TF1, TF2, and TF3). The position and proportion for individual SNSs are shown in different colors as indicated on the top-left side of the figure. 5'-TC-3' sites and 5'-GA-3' sites are shown in green and red respectively. Vertical axes are adjusted for comparison.
