## Supplementary material for "Systematic mutagenesis assay promotes comprehension of the strand-bias laws for mutations induced by oxidative DNA damage": Figure 3-figure supplement 1

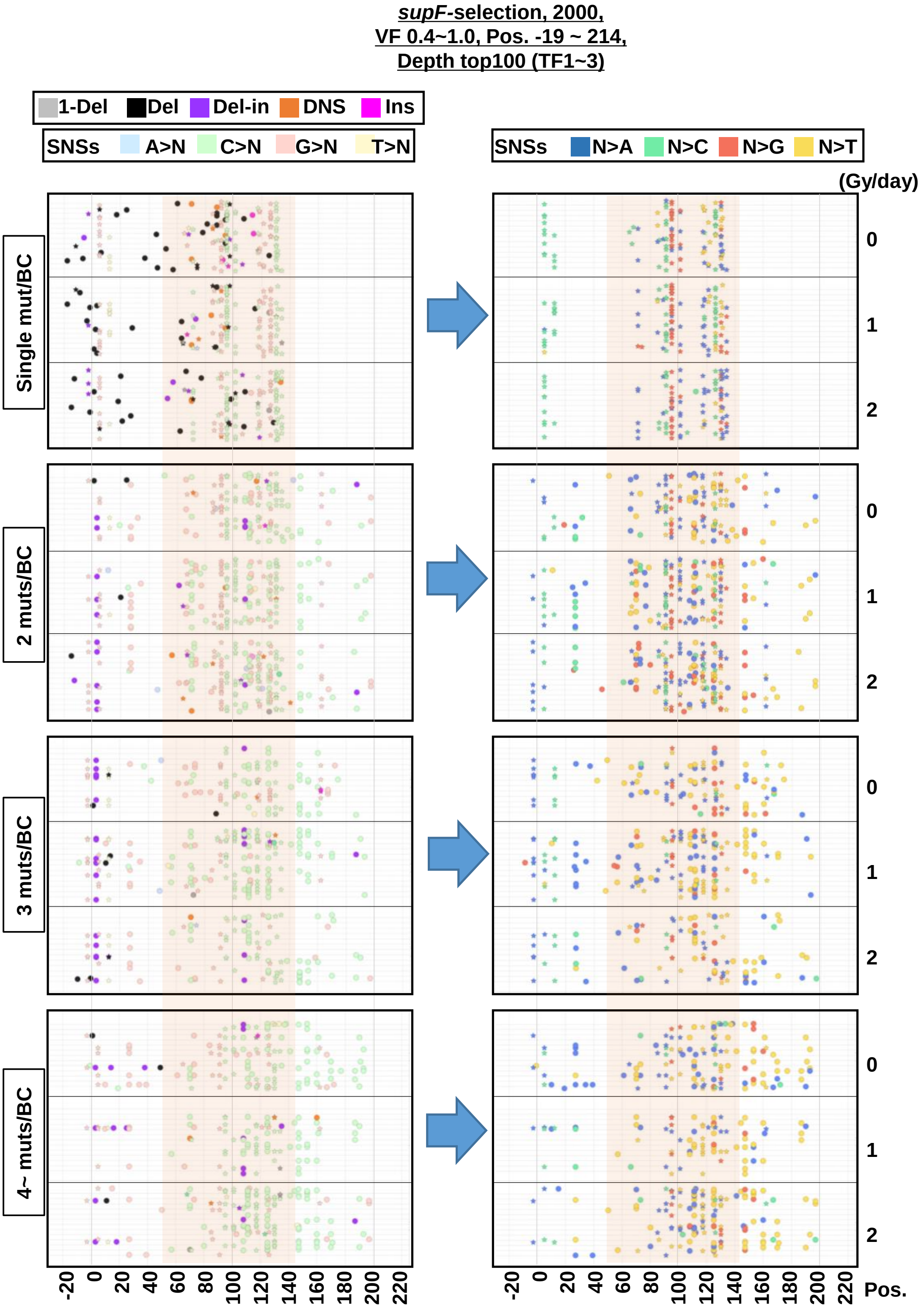


**Fig. 3-figure supplement 1.** The position and mutation type of each identified mutation (*supF*-selection). The position and mutation type of each identified mutation are presented in dot plots for the cases of 1, 2, 3, or 4~ mutations per $N_{12}$-BC. The *supF*-tRNA cloverleaf-region is highlighted by light-red background. Each type of mutation is indicated by a different color in the left-side plots (legend on top). The bases after the substitution are indicated by a different color for only SNSs in the right-side plots (legend on top). The mutations in identical BCs are plotted on the same horizontal line. Data is combined from TF1, TF2, and TF3.
