## Supplementary material for "Systematic mutagenesis assay promotes comprehension of the strand-bias laws for mutations induced by oxidative DNA damage": Figure 3-figure supplement 2

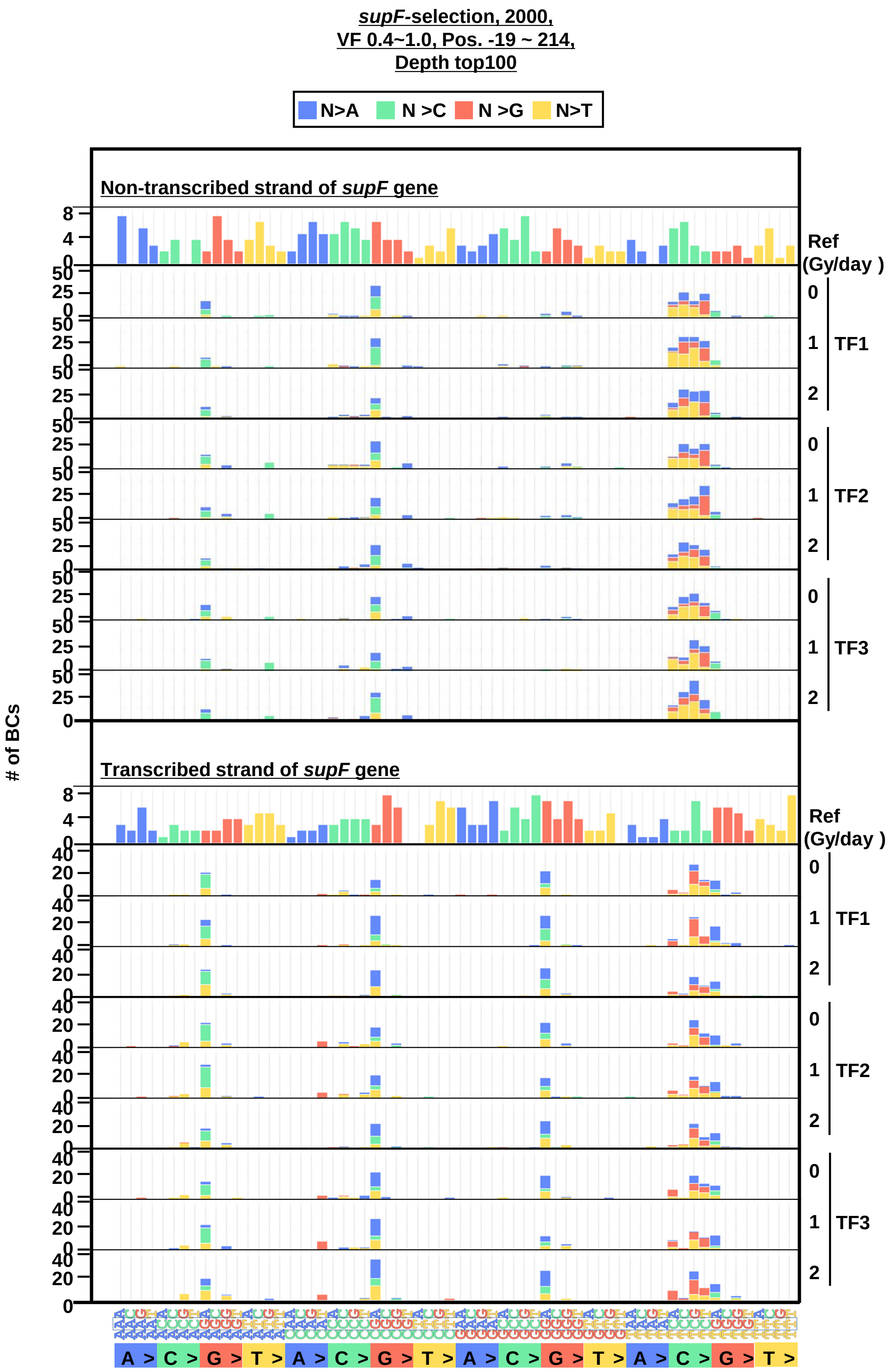


**Fig. 3-figure supplement 2.** The 192 trinucleotide contexts of chronic gamma-irradiation-induced SNSs (*supF*-selection). The number of SNSs in192 trinucleotide contexts for the non-transcribed (top panel) and transcribed (bottom panel) strand of the *supF* gene (nucleotide positions from -19 to 214). The substituted bases are indicated in colors corresponding to the notation on the horizontal axis; the transfection experiments (TF1, TF2, and TF3) and dose-rates (0, 1, and 2 Gy/day) are shown on the right side of the graph; the reference sequence for the analysis is denoted as Ref.
