## Supplementary material for "Systematic mutagenesis assay promotes comprehension of the strand-bias laws for mutations induced by oxidative DNA damage": Figure 4-figure supplement 1

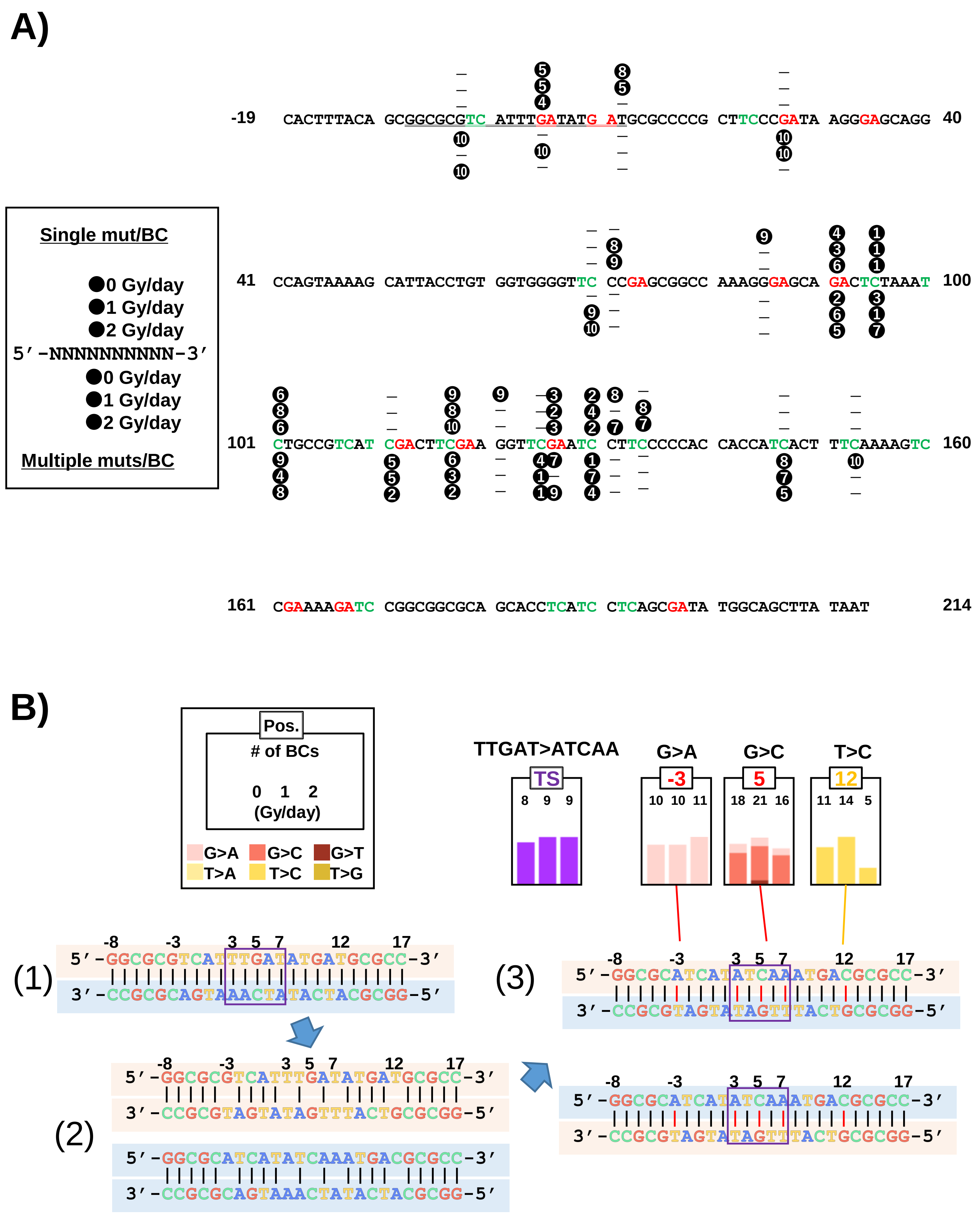


**Fig. 4-figure supplement 1.** The *supF* mutant phenotype resulting from either single mutations per $N_{12}$-BC or template switching at the specific regions in the analyzed sequence. (A) Comparison of the ranking by frequency of detection between single mutation per $N_{12}$-BC and multiple mutations per $N_{12}$-BC. The analyzed nucleotide sequence of the non-transcribed strand (sense strand, positions -19 to 214) with a number in black circle corresponding to the rank indicated in Figure 4A, as presented in the upper-left box.(B) The characteristic template switching mutations (TS, TTGAT>ATCAA) at the potential hairpin loop of the quasi-palindromic sequence. Reference sequence (1), the template switched sequences (2), and the mutated sequence (3). The potential Watson-Crick hydrogen bonded base pair is shown by a vertical line. The SNSs at positions -3, 5 and 12 are detected as indicated in the box, and for deletion/insertion at positions 3 to 7 – in purple color. The proportion of individual base-substitutions is shown in different colors as indicated in the upper-left box.
