## Supplementary material for "Systematic mutagenesis assay promotes comprehension of the strand-bias laws for mutations induced by oxidative DNA damage": Figure 4-figure supplement 2

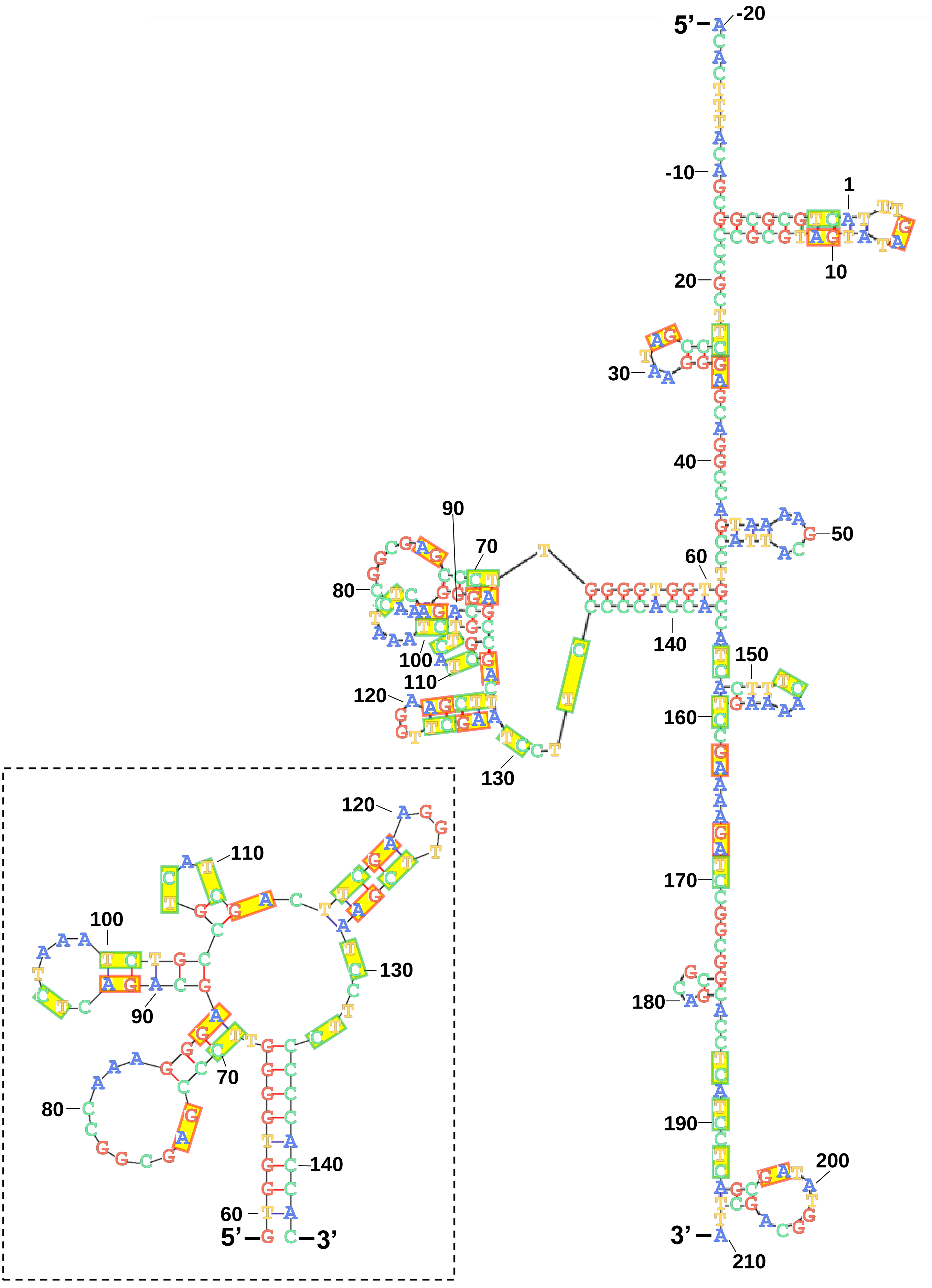


**Fig. 4-figure supplement 2.** The single-stranded DNA secondary structure for the non-transcribed strand (sense strand) of the analyzed sequence (either from -20 to 210, or from 59 to 143 in the dashed-line box). Mfold web server for nucleic acid folding and hybridization prediction for DNA folding form was used (Folding at 37°C, and ionic conditions were $Na^{+}$=10 mM, $Mg^{++}$=1 mM). The 5'-TC-3' sites are shown in the green line boxex, and the 5'-GA-3' sites are shown in the red line boxes.
