## Supplementary material for "Systematic mutagenesis assay promotes comprehension of the strand-bias laws for mutations induced by oxidative DNA damage": Figure 4-figure supplement 3

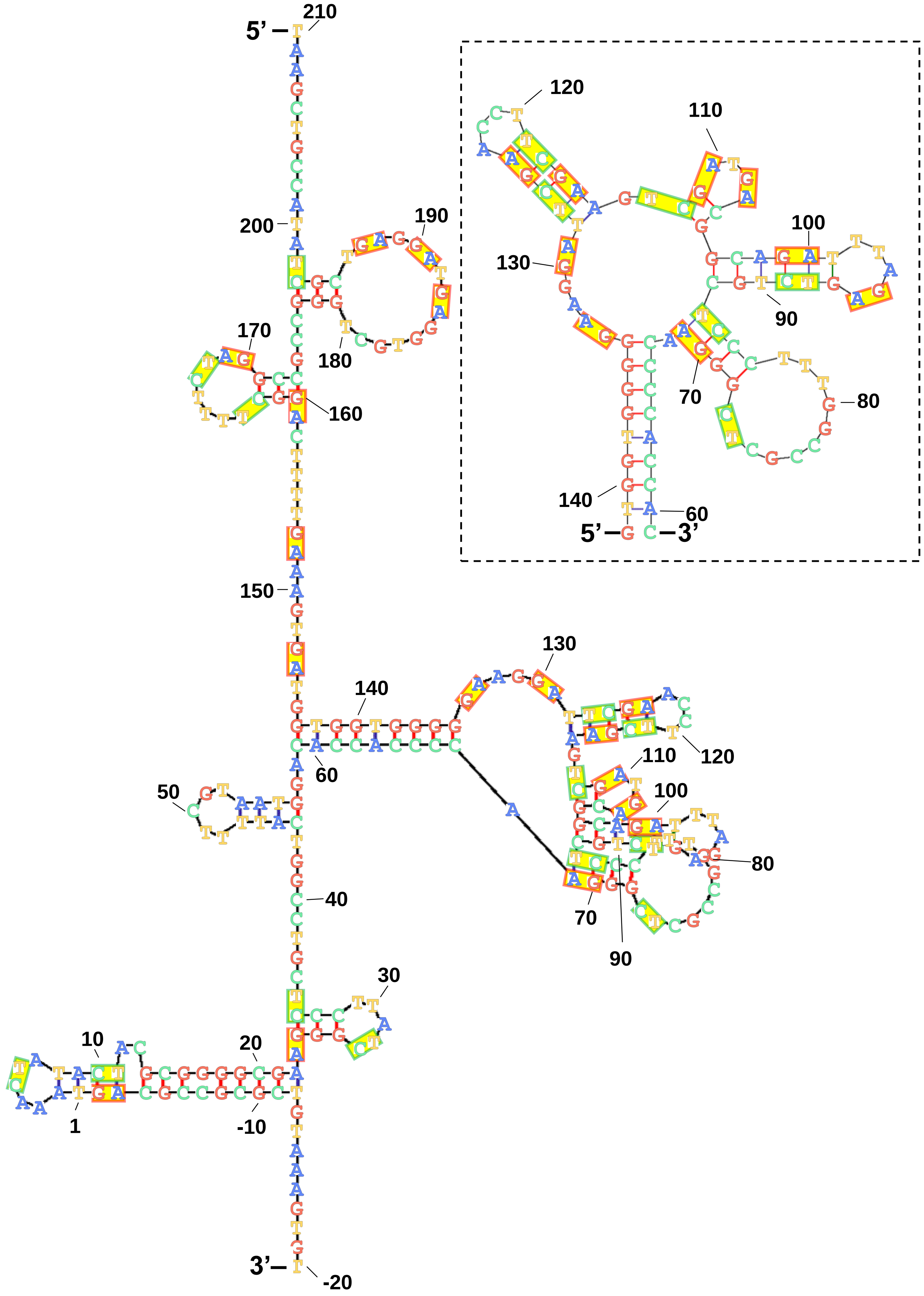


**Fig. 4-figure supplement 3.** The single-stranded DNA secondary structure for the transcribed strand (antisense strand) of the analyzed sequence (either from -20 to 210, or from 59 to 143 in the dashed-line box), as shown in Fig.4-figure supplement 2.
