## Supplementary material for "Systematic mutagenesis assay promotes comprehension of the strand-bias laws for mutations induced by oxidative DNA damage": Figure 4-figure supplement 8

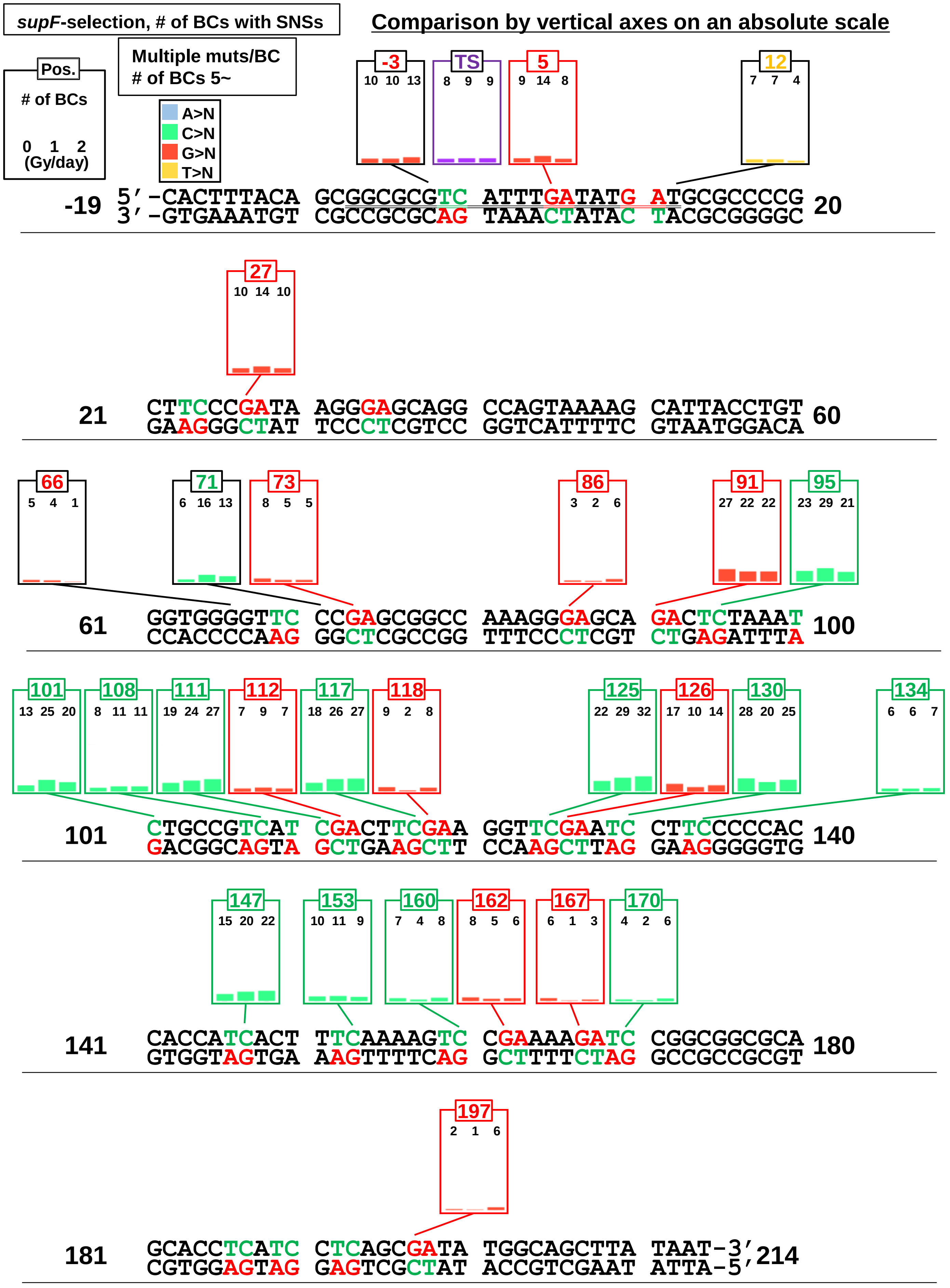


**Fig. 4-figure supplement 8.** The number of SNSs for single mutation per $N_{12}$-BC according to their positions and the dose-rates of irradiation (*supF*-selection). Bar graphs provide combined data from three transfection experiments (TF1, TF2, and TF3). The position and reference base for individual SNSs are shown in different colors as indicated on the top-left side of the figure. 5'-TC-3' sites and 5'-GA-3' sites are shown in green and red respectively. Vertical axes are on an absolute scale adjusted for comparison.
