## Supplementary material for "Systematic mutagenesis assay promotes comprehension of the strand-bias laws for mutations induced by oxidative DNA damage": Figure 4-figure supplement 9

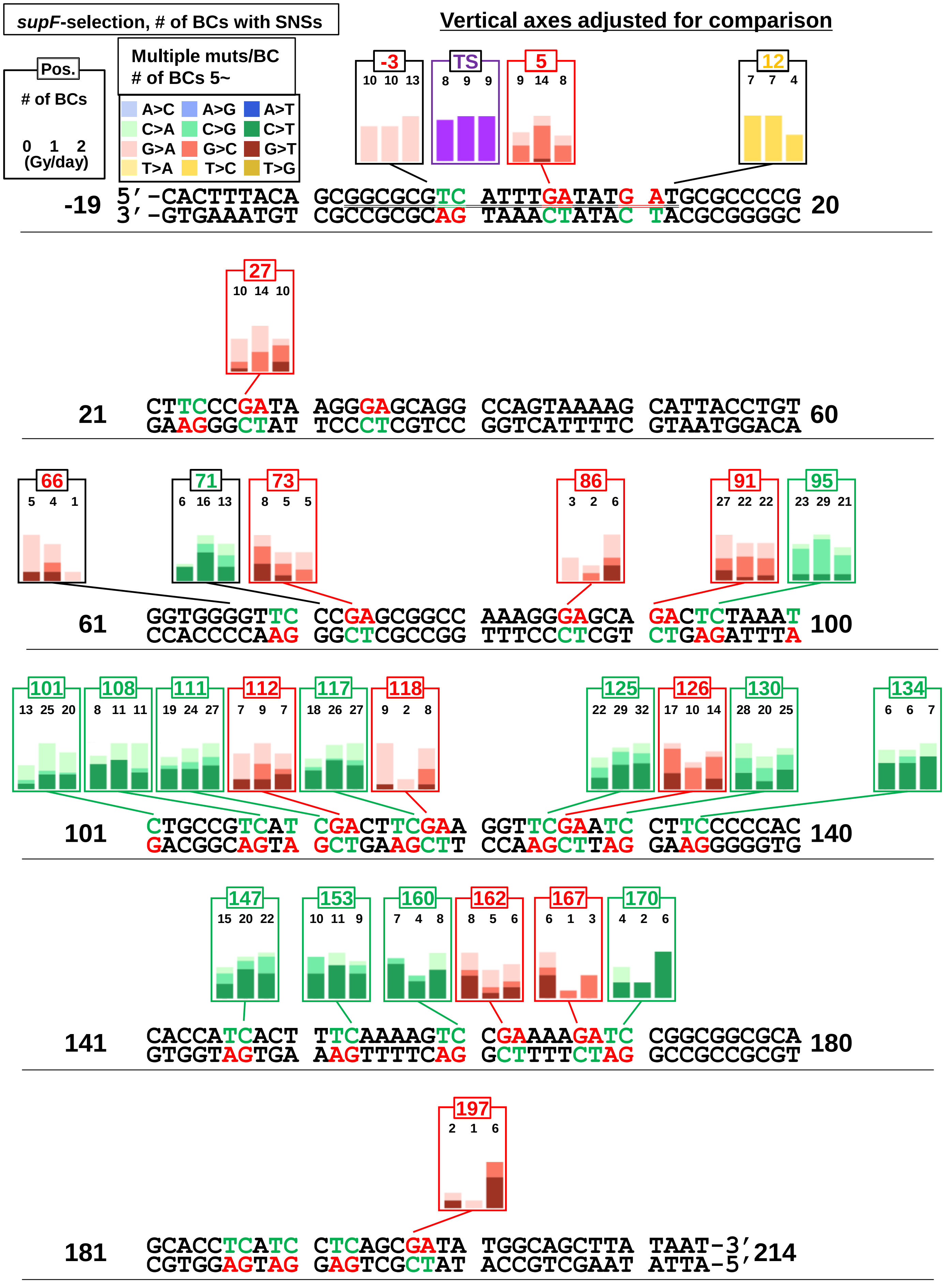


**Fig. 4-figure supplement 9.** Proportion of substituted bases in SNSs for multiple mutations per $N_{12}$-BC according to their positions and the dose-rates of irradiation (*supF*-selection). Bar graphs provide the number of SNSs for the combined data from three transfection-experiments (TF1, TF2, and TF3). The position and reference base for individual SNSs are shown in different colors as indicated on the top-left side of the figure. 5'-TC-3' sites and 5'-GA-3' sites are shown in green and red respectively. Vertical axes are adjusted for comparison.
