## Supplementary material for "Systematic mutagenesis assay promotes comprehension of the strand-bias laws for mutations induced by oxidative DNA damage": Figure 5-figure supplement 2

## A)

| SNSs | 0 Gy/day | 1 Gy/day | 2 Gy/day |
|---|---|---|---|
| *w/o-supF*-selection | ① 27 CCGAT/ATCGG (29)<br>② 5 TTGAT/ATCAA (15)<br>③ 147 ATCAC/GTGAT (15)<br>④ 125 TTCGA/TCGAA (13)<br>⑤ -3 GCGTC/GACGC (9)<br>⑤ 95 CTCTA/TAGAG (9)<br>⑤ 111 ATCGA/TCGAT (9)<br>⑧ 91 CAGAC/GTCTG (8)<br>⑨ 12 GATGC/GCATC (7)<br>⑨ 130 ATGCG/CGCAT (7)<br>⑨ 153 TTCAA/TTGAA (7)<br>⑨ 162 CCGAA/TTCGG (7)<br>*TS TTGAT/ATCAA (5) | ① 27 CCGAT/ATCGG (48)<br>② 111 ATCGA/TCGAT (35)<br>③ 5 TTGAT/ATCAA (30)<br>④ 125 TTCGA/TCGAA (28)<br>⑤ 95 CTCTA/TAGAG (20)<br>⑥ 73 CCGAG/CTCGG (14)<br>⑥ 197 GCGAT/ATCGC (14)<br>⑧ 91 CAGAC/GTCTG (13)<br>⑧ 147 ATCAC/GTGAT (13)<br>⑧ -3 GCGTC/GACGC (12)<br>*TS TTGAT/ATCAA (6) | ① 27 CCGAT/ATCGG (56)<br>② 5 TTGAT/ATCAA (35)<br>② 111 ATCGA/TCGAT (35)<br>④ 125 TTCGA/TCGAA (19)<br>⑤ -3 GCGTC/GACGC (18)<br>⑥ 147 ATCAC/GTGAT (16)<br>⑥ 197 GCGAT/ATCGC (16)<br>⑧ 95 CTCTA/TAGAG (14)<br>⑧ 117 GATGC/GCATC (14)<br>⑩ 91 CAGAC/GTCTG (13)<br>⑩ 101 ATCTG/CAGAT (13)<br>⑩ 130 ATGCG/CGCAT (13)<br>*TS TTGAT/ATCAA (3) |
| *supF*-selection | ❶ 130 ATCCT/AGGAT (44)<br>❷ 95 CTCTA/TAGAG (43)<br>❸ 91 CAGAC/GTCTG (39)<br>❹ 126 TCGAA/TTCGA (30)<br>❺ 125 TTCGA/TCGAA (22)<br>❻ 101 ATCTG/CAGAT (21)<br>❻ 117 TTCGA/TCGAA (21)<br>❽ 111 ATCGA/TCGAT (19)<br>❾ 5 TTGAT/ATCAA (18)<br>❿ 118 TCGAA/TTCGA (16)<br>*TS TTGAT/ATCAA (8) | ❶ 95 CTCTA/TAGAG (54)<br>❷ 91 CAGAC/GTCTG (31)<br>❸ 101 ATCTG/CAGAT (29)<br>❸ 117 TTCGA/TCGAA (29)<br>❸ 125 TTCGA/TCGAA (29)<br>❻ 130 ATCCT/AGGAT (28)<br>❼ 126 TCGAA/TTCGA (25)<br>❽ 111 ATCGA/TCGAT (24)<br>❾ 5 TTGAT/ATCAA (21)<br>❿ 147 ATCAC/GTGAT (20)<br>*TS TTGAT/ATCAA (9) | ❶ 130 ATCCT/AGGAT (40)<br>❷ 95 CTCTA/TAGAG (39)<br>❸ 125 TTCGA/TCGAA (32)<br>❹ 117 TTCGA/TCGAA (31)<br>❺ 91 CAGAC/GTCTG (30)<br>❻ 101 ATCTG/CAGAT (29)<br>❼ 111 ATCGA/TCGAT (27)<br>❽ 126 TCGAA/TTCGA (24)<br>❾ 147 ATCAC/GTGAT (22)<br>❿ 5 TTGAT/ATCAA (17)<br>*TS TTGAT/ATCAA (9) |

## B)

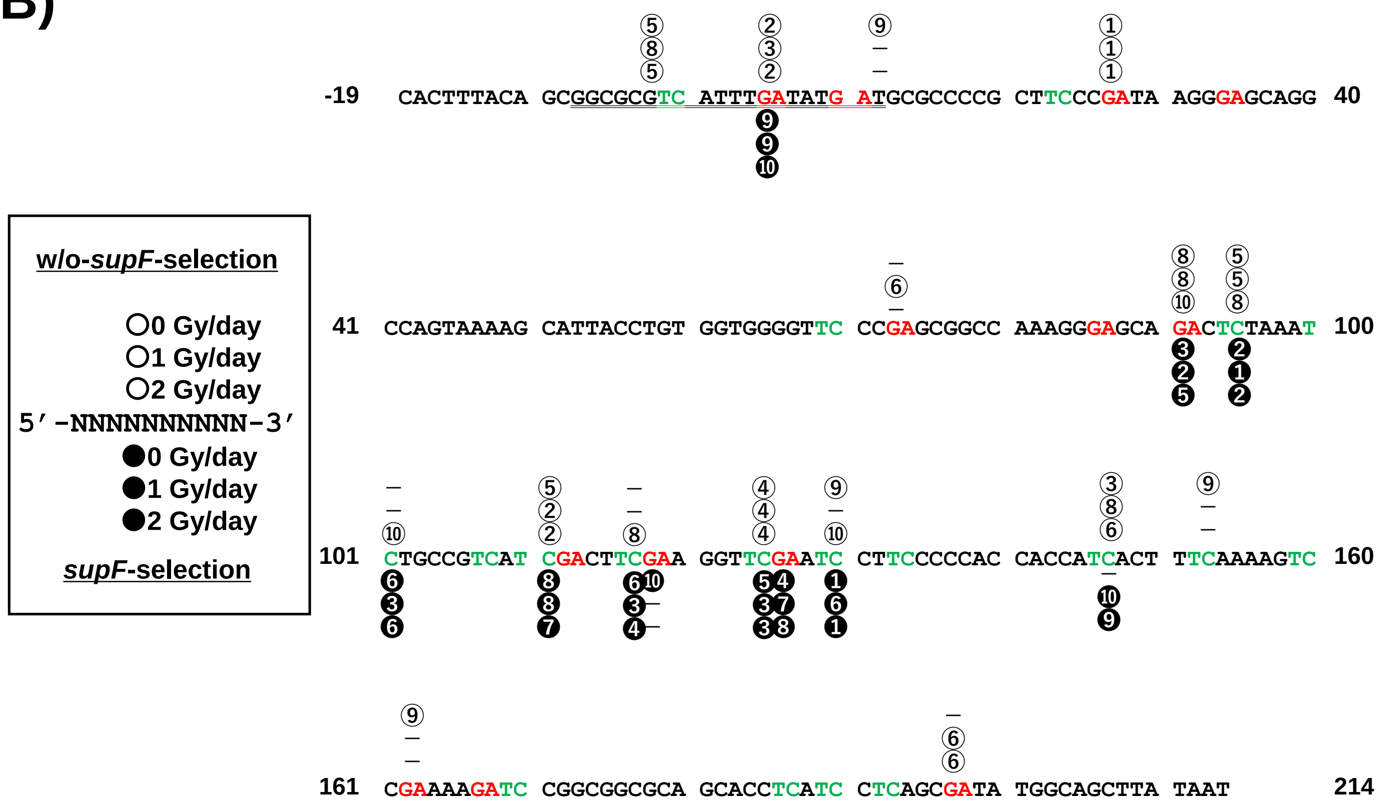


**Fig. 5-figure supplement 2.** Comparison of the ranking of SNSs by frequency of detection between w/o-*supF*-selection and *supF*-selection. (A) The ranks were listed for w/o-*supF*-selection (the upper row) or *supF*-selection (the lower row) (refer to the legend of Figure 4A). (B) The analyzed nucleotide sequence of the non-transcribed strand (sense strand, positions -19 to 214) with a number in a white circle correspond to the correspond to the rank referred to w/o-*supF*-selection, and black circle correspond to the rank referred to *supF*-selection, as presented in the upper box.
