## Supplementary material for "Systematic mutagenesis assay promotes comprehension of the strand-bias laws for mutations induced by oxidative DNA damage": Figure 8-figure supplement 1

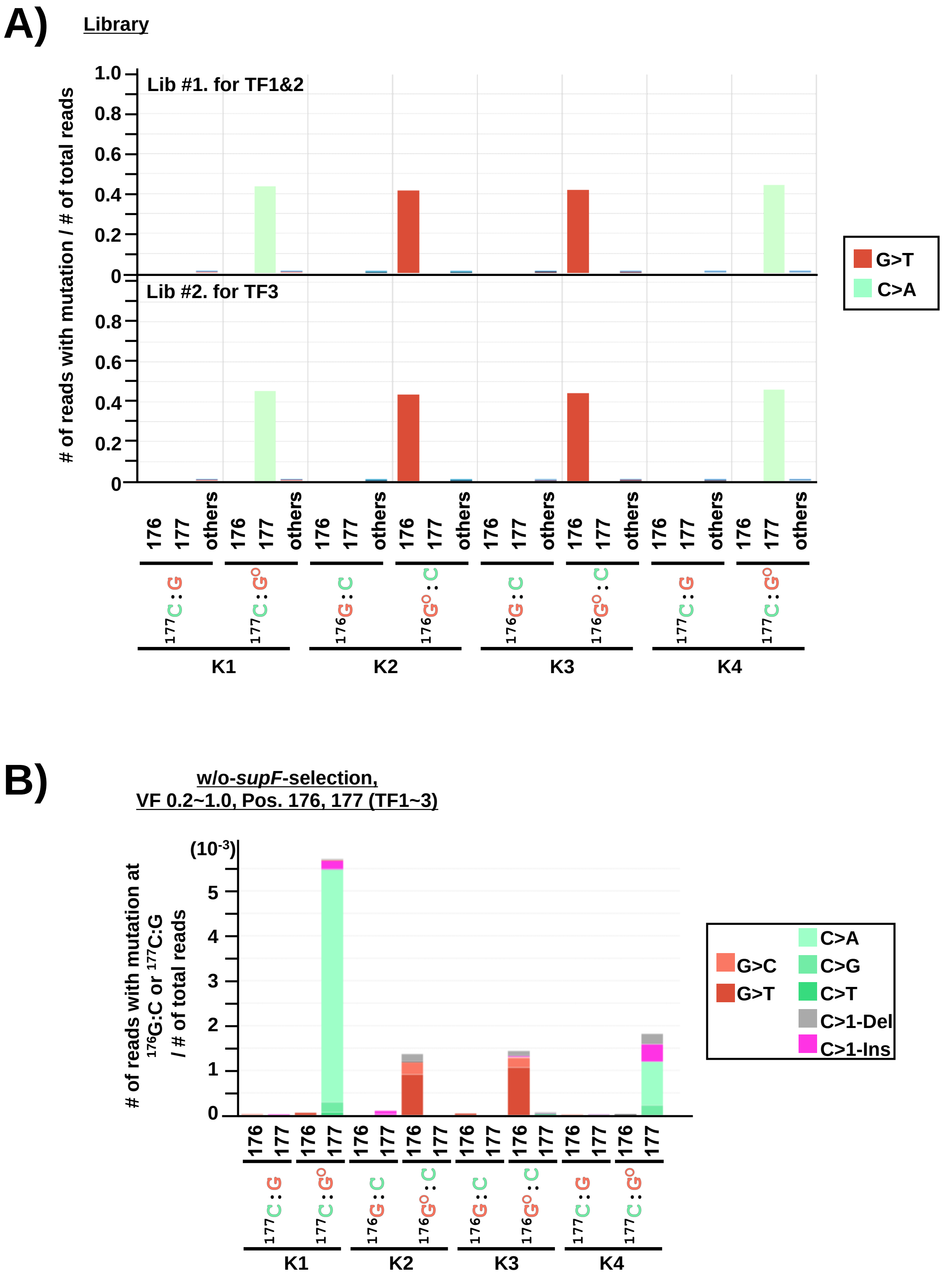


**Fig. 8-figure supplement 1.** The sequencing analysis for the sites inserted with an 8-oxo-G (at 176 for K2 and K3, at 177 for K1 and K4). (A) The sequence of two $N_{12}$-BC libraries independently constructed *in vitro* for each pNGS2 -K1, -K2, -K3, and –K4 (Library #1 for TF1 and TF2, Library #2 for TF3), was analyzed by NGS before the transfection experiment. The background mutation frequencies for each library were determined. The mutation frequencies at positions 176 and 177 are shown separately from the rest of the sequence (denoted "others"). (B) The mutation frequencies at the 8-oxo-G inserted sites for each library from the data of w/o-*supF*-selection. The mutation frequency was calculated by dividing the number of reads with mutation at the site of insertion by the number of total reads. The combined data from three transfection-experiments is shown in a bar graph. The mutation types and substituted bases are shown in different colors as indicated on the right side of the graph.
