## Supplementary material for "Systematic mutagenesis assay promotes comprehension of the strand-bias laws for mutations induced by oxidative DNA damage": Figure 10-figure supplement 1

<u>*w/o-supF*-selection,</u>
<u>VF 0.2~1.0. Pos. -19 ~ 214 (TF1~3)</u>

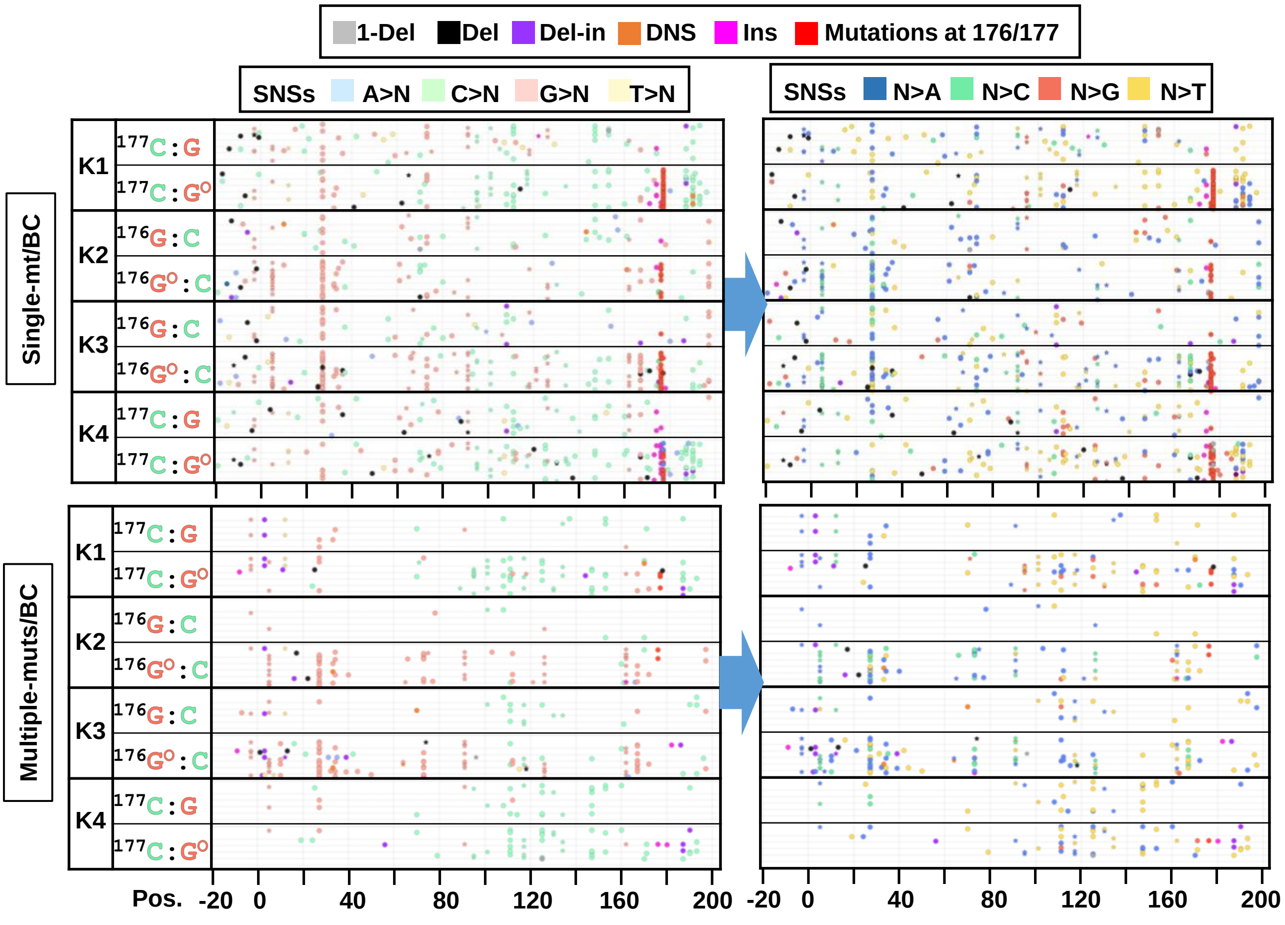


**Fig. 10-figure supplement 1.** The position and mutation type of each identified mutation (w/o-*supF*-selection). The position and mutation type of each identified mutation are presented in dot plots for the cases of either single mutation per $N_{12}$-BC (upper plots) or multiple mutations per $N_{12}$-BC (lower plots). Each type of mutation and reference bases for SNSs are indicated by a different color in the left-side plots (legend on top). The bases after SNSs are indicated by a different color for only SNSs in the right-side plots (legend on top). The mutations in identical BCs are plotted on the same horizontal line. Data is combined from TF1, TF2, and TF3.
