## Supplementary material for "Systematic mutagenesis assay promotes comprehension of the strand-bias laws for mutations induced by oxidative DNA damage": Figure 11-figure supplement 1

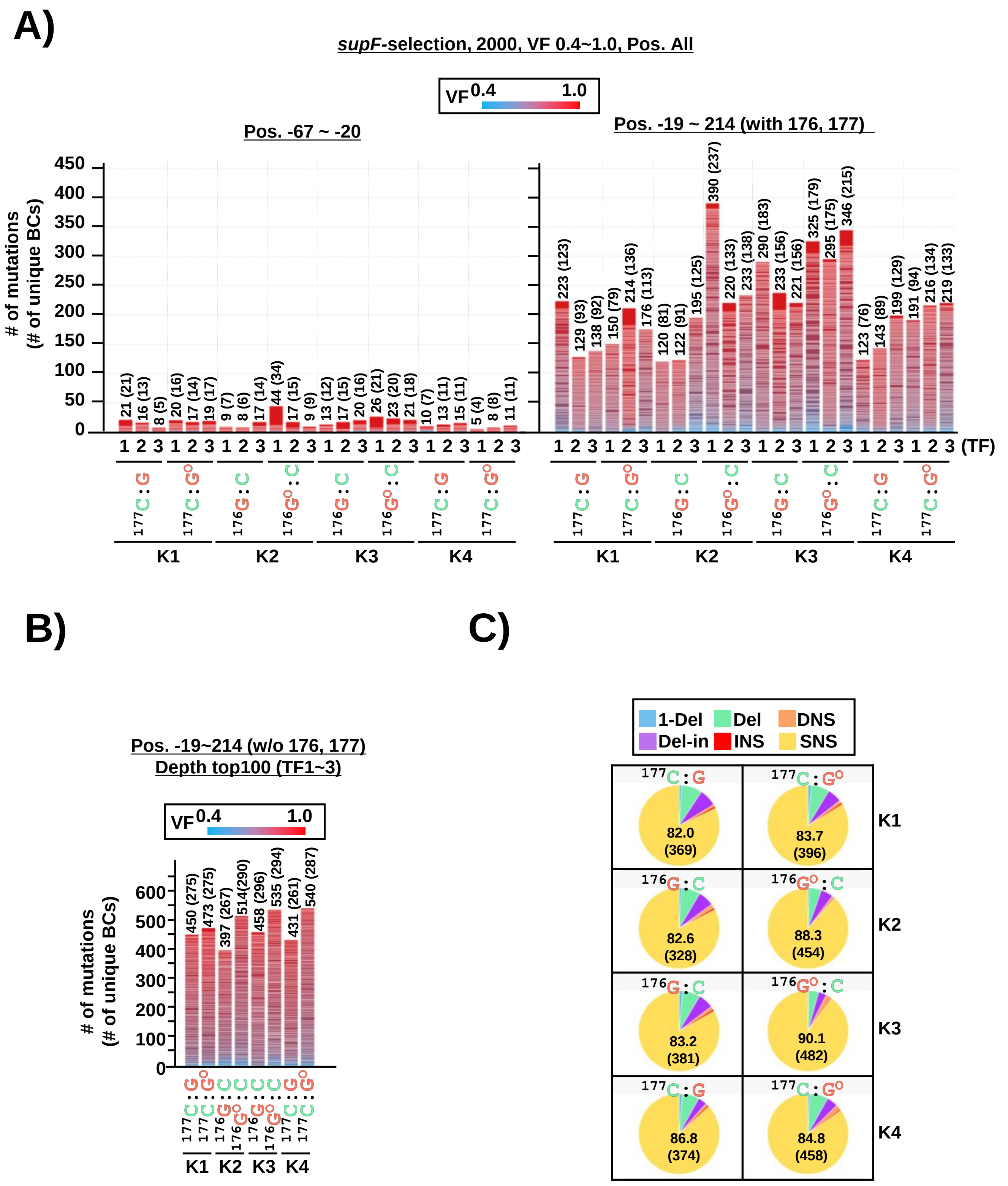


**Fig. 11-figure supplement 1.** The *supF* NGS assay for the mutations induced by the insertion of an 8-oxo-G (*supF*-selection, VF 0.4~1.0). (A) Stack bar graph representing the number of $N_{12}$-BC sequences with mutations obtained from each sample used for data analysis. The data were separated into two graphs: the graph on the left side represents positions from -67 to -20, and the graph on the right side – positions from -19 to 214 containing the *supF* gene. (B) Stack bar graph representing the number of $N_{12}$-BC sequences with mutations obtained from each sample used for data analysis. The one hundred $N_{12}$-BCs with at least one mutation or the maximum number of available $N_{12}$-BCs with mutations extracted from the group with the highest number of reads per unique $N_{12}$-BC. The number in parentheses refers to the number of unique $N_{12}$-BC sequences with mutations. (C) Pie charts of the proportions of mutation types in different colors as indicated at the top of the panel. The percentage (the number) of SNSs are provided in each pie chart.
