## Supplementary material for "Systematic mutagenesis assay promotes comprehension of the strand-bias laws for mutations induced by oxidative DNA damage": Figure 15-figure supplement 4

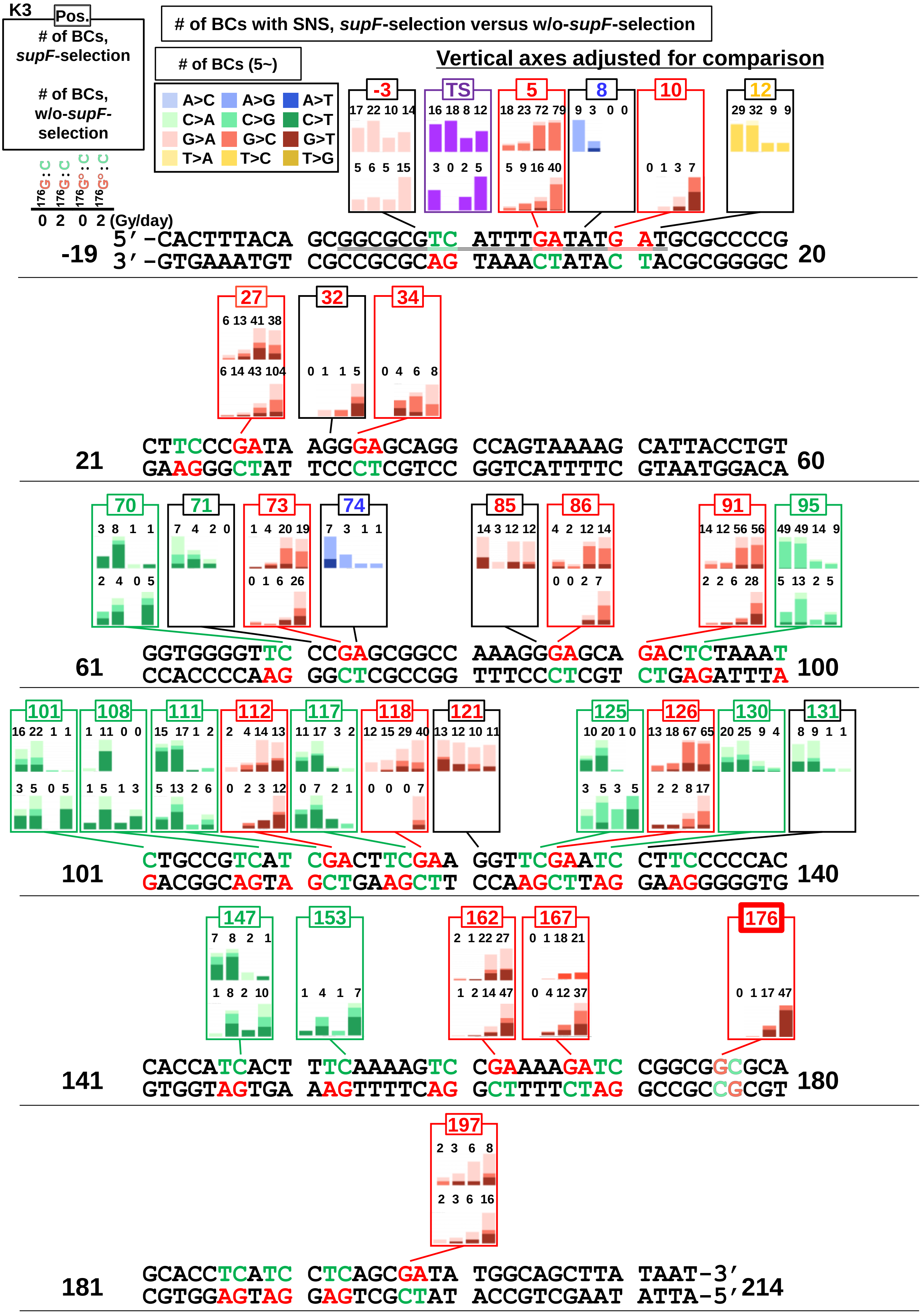


**Fig. 15-figure supplement 4.** Proportion of substituted bases in SNSs according to their positions for pNGS2-K3 inserted with either a G or an 8-oxo-G and irradiation (*supF*-selection versus w/o-*supF*-selection). The layout of the figure is analogous to Figure 12. Scales are adjusted for comparison.
