## Supplementary material for "Systematic mutagenesis assay promotes comprehension of the strand-bias laws for mutations induced by oxidative DNA damage": Figure 15-figure supplement 6

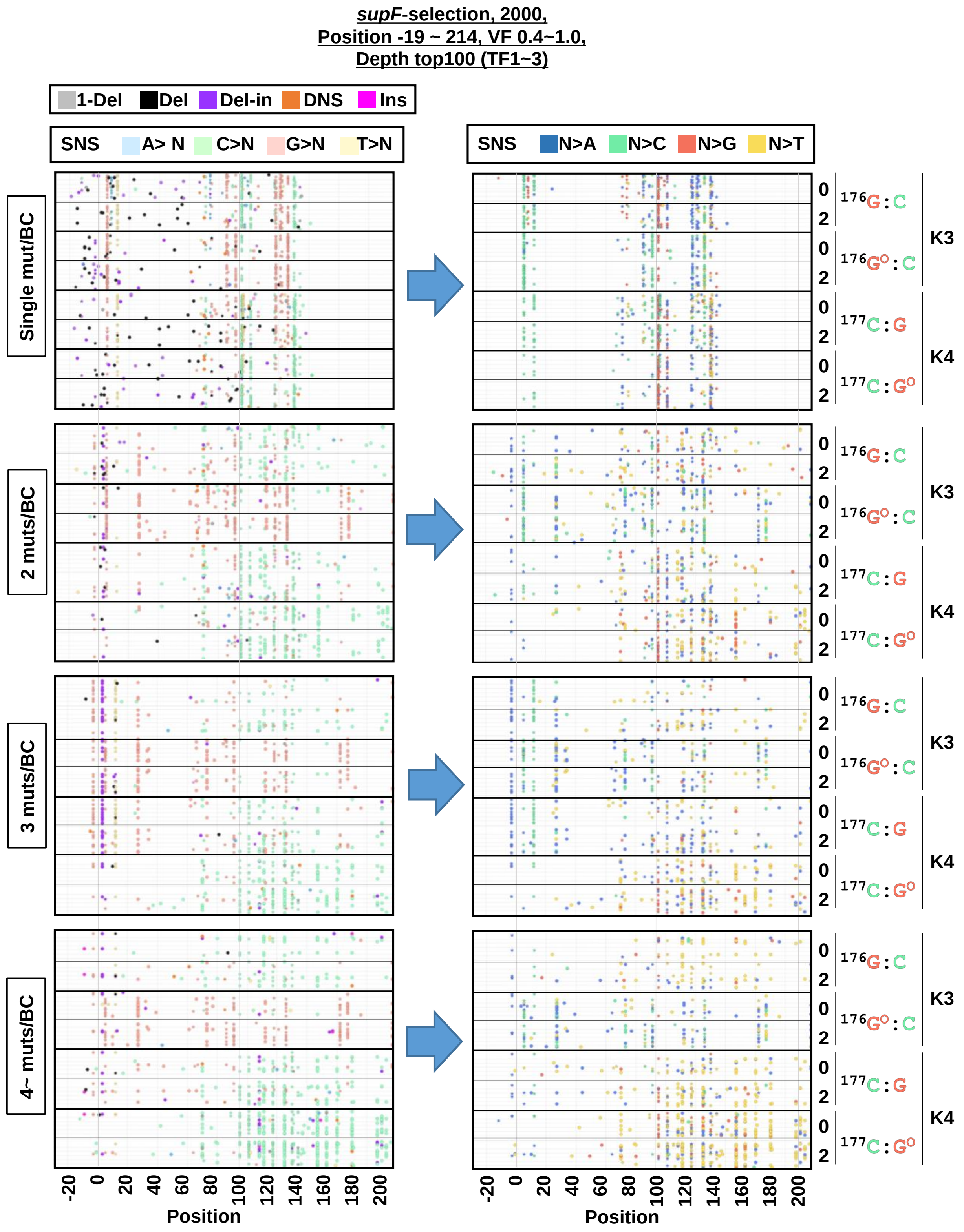


**Fig. 15-figure supplement 6.** The position and mutation type of each identified mutation (*supF*-selection, VF 0.4~1.0). The position and mutation type of each identified mutation are presented in dot plots for the cases of either single (1), 2, 3, or more than 3 (4~) mutations per $N_{12}$-BC. Each type of mutation and reference bases for SNSs are indicated by a different color in the left-side plots (legend on top). The bases after SNSs are indicated by a different color for only SNSs in the right-side plots (legend on top). The mutations in identical BCs are plotted on the same horizontal line. Data is combined from TF1, TF2, and TF3.
