## Supplementary material for "Systematic mutagenesis assay promotes comprehension of the strand-bias laws for mutations induced by oxidative DNA damage": Figure 15-figure supplement 11

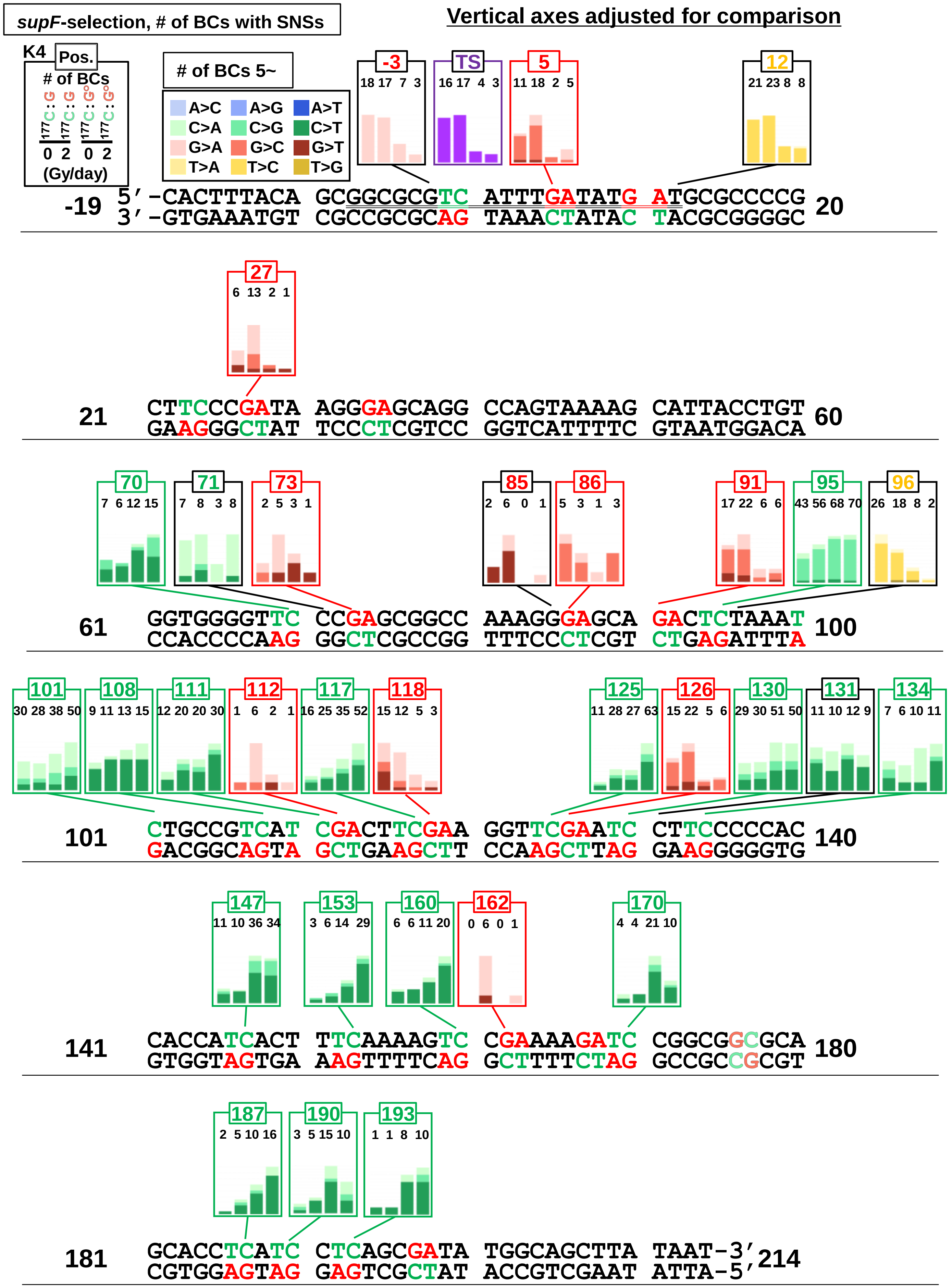


**Fig. 15-figure supplement 11.** Effects of 8-oxo-G and chronic gamma-irradiation (2 Gy/day) on the proportion of substituted bases in SNSs according to their positions (pNGS2-K4, *supF*-selection). Bar graphs provide the number of SNSs for the combined data from three transfection-experiments (TF1, TF2, and TF3). The both position and reference base for individual SNSs are shown in different colors as indicated on the top-left side of the figure. 5'-TC-3' sites and 5'-GA-3' sites are shown in green and red respectively. Comparison with adjusted scales for each graph.
