## Supplementary material for "Systematic mutagenesis assay promotes comprehension of the strand-bias laws for mutations induced by oxidative DNA damage": Figure 13-figure supplement 2

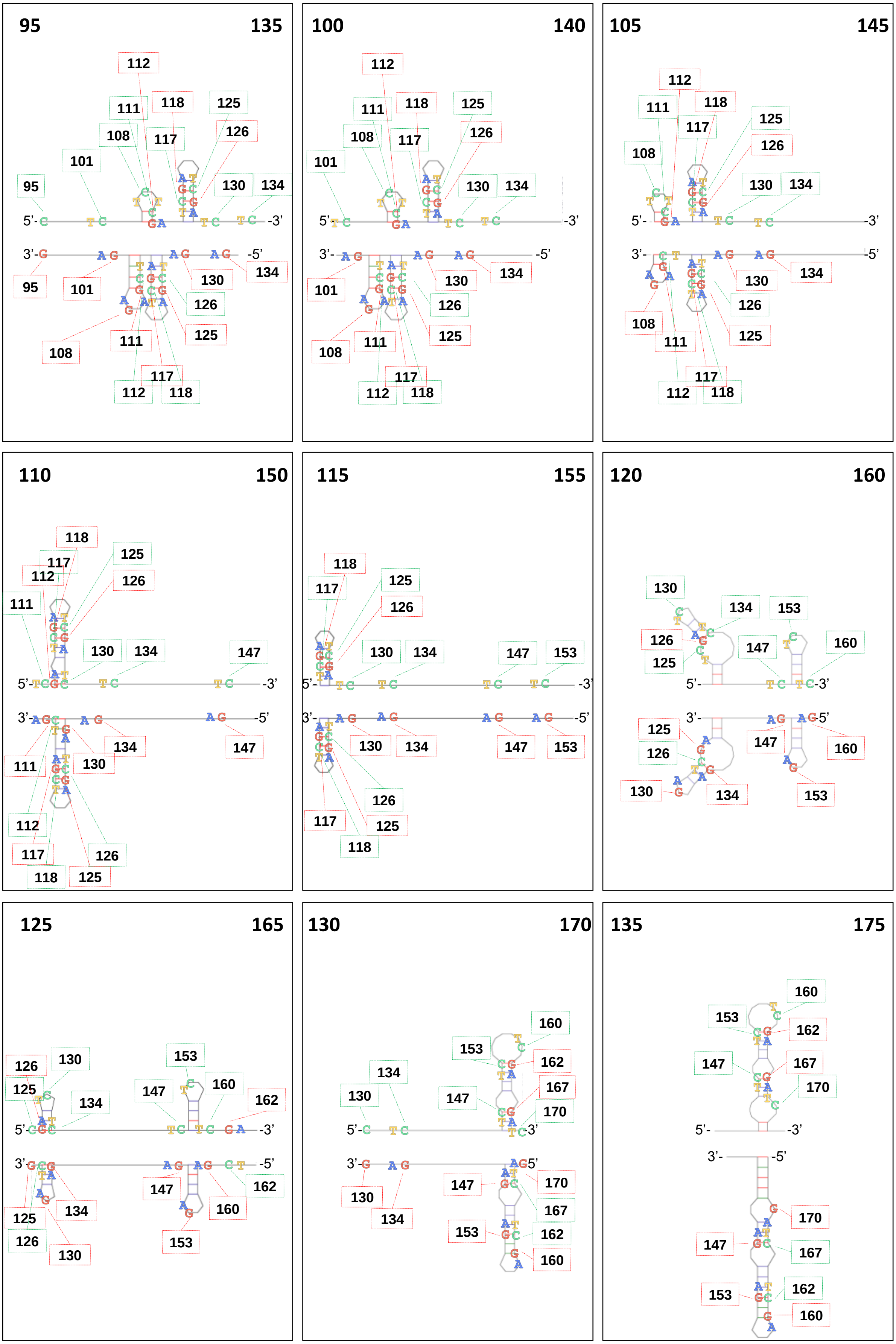


**Fig. 13-figure supplement 2.** The localizations of each 5’-TC-3’:5’-GA-3’ site and the predicted secondary structure for single-stranded DNA (positions from 95 to 175)
