## Supplementary material for "Systematic mutagenesis assay promotes comprehension of the strand-bias laws for mutations induced by oxidative DNA damage": Figure 14-figure supplement 1

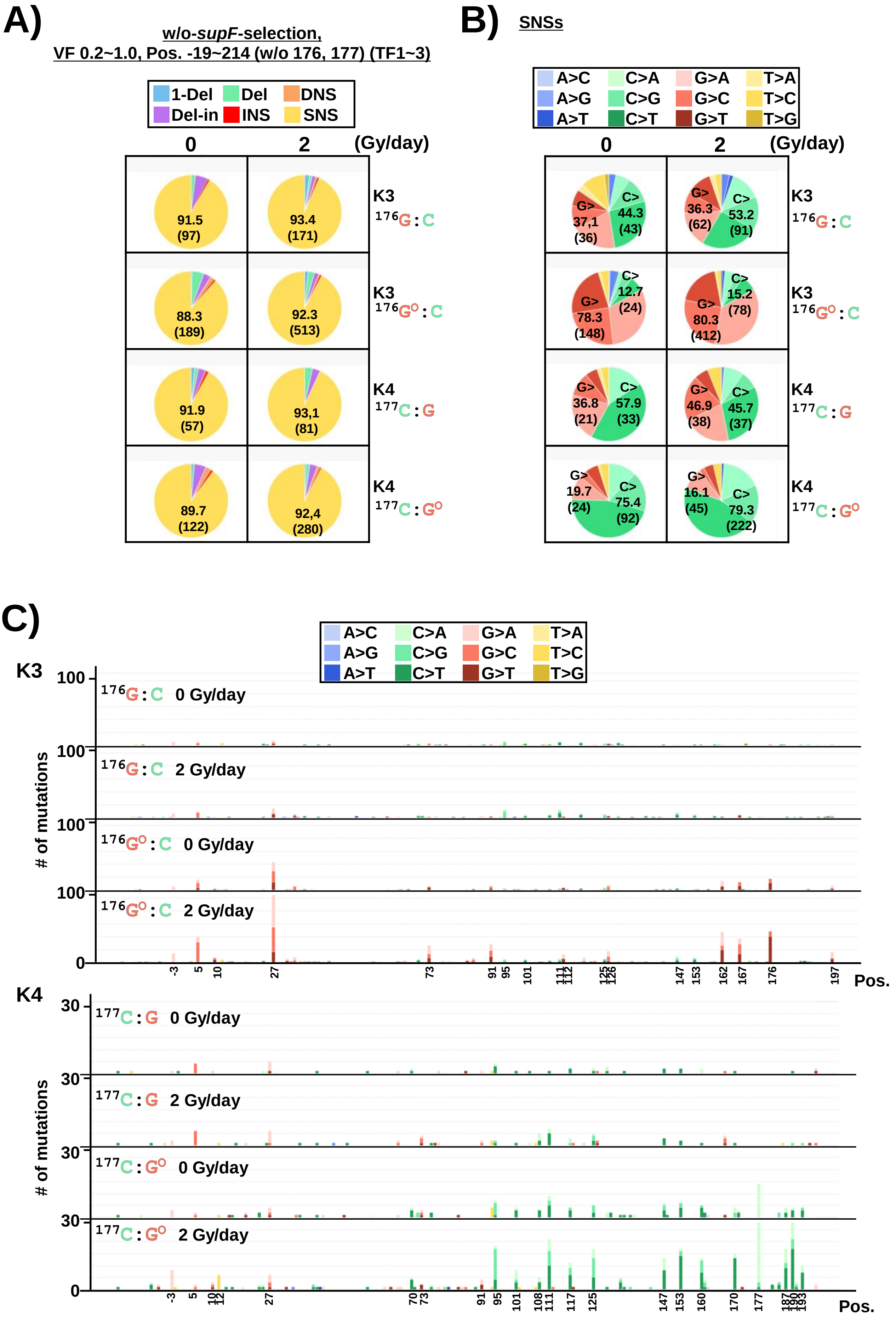


**Fig. 14-figure supplement 1.** The effect of chronic gamma-irradiation on 8-oxo-G induced action-at-a-distance mutations in cells (pNGS2-K3 and K4 inserted with either a non-damaged G or an 8-oxo-G, dose-rate of irradiation at 0 or 2 Gy/day for 2 days, w/o-supF-selection, VF 0.2~1.0) (A) Pie charts of the proportions of mutation types (refer to the legend in Figure 2C). (B) Pie charts of the proportions of different base-substitutions for SNSs (refer to the legend in Figure 2D). (C) Number of SNSs according to their nucleotide position (refer to the legend in Figure 2E).
