## Supplementary material for "Systematic mutagenesis assay promotes comprehension of the strand-bias laws for mutations induced by oxidative DNA damage": Figure 14-figure supplement 4

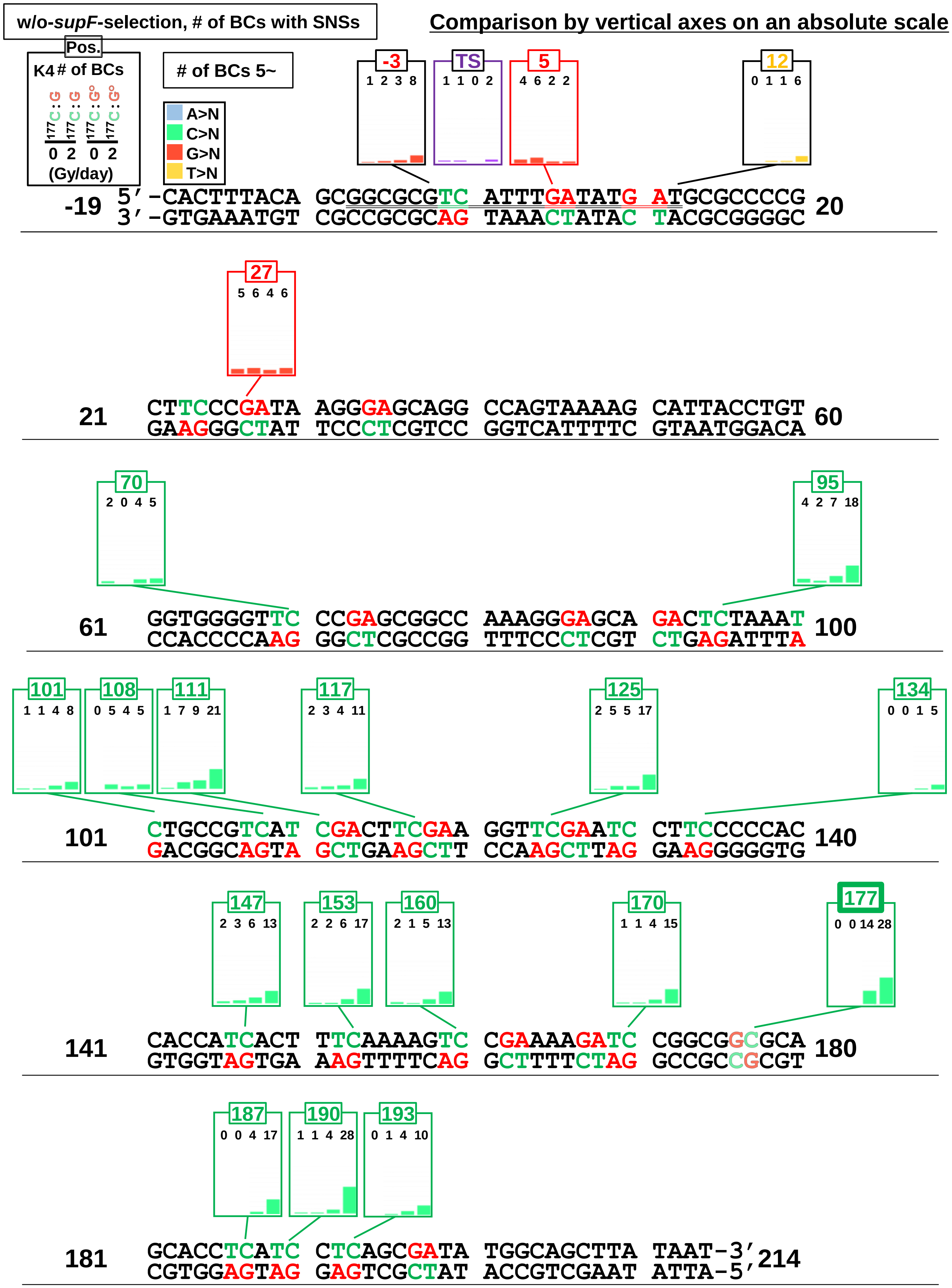


**Fig. 14-figure supplement 4.** Effects of 8-oxo-G insertion and chronic gamma-irradiation at 2 Gy/day on the number of SNSs according to their positions in the sequence (pNGS2-K4, w/o-*supF*-selection). Bar graphs provide combined from three transfection experiments (TF1, TF2, and TF3). The positions and reference bases for individual SNSs are shown in different colors as indicated on the top-left side of the figure. 5'-TC-3' sites and 5'-GA-3' sites are shown in green and red respectively. Vertical axes are  on an absolute scale adjusted for comparison.
