## Supplementary material for "Systematic mutagenesis assay promotes comprehension of the strand-bias laws for mutations induced by oxidative DNA damage": Figure 14-figure supplement 5

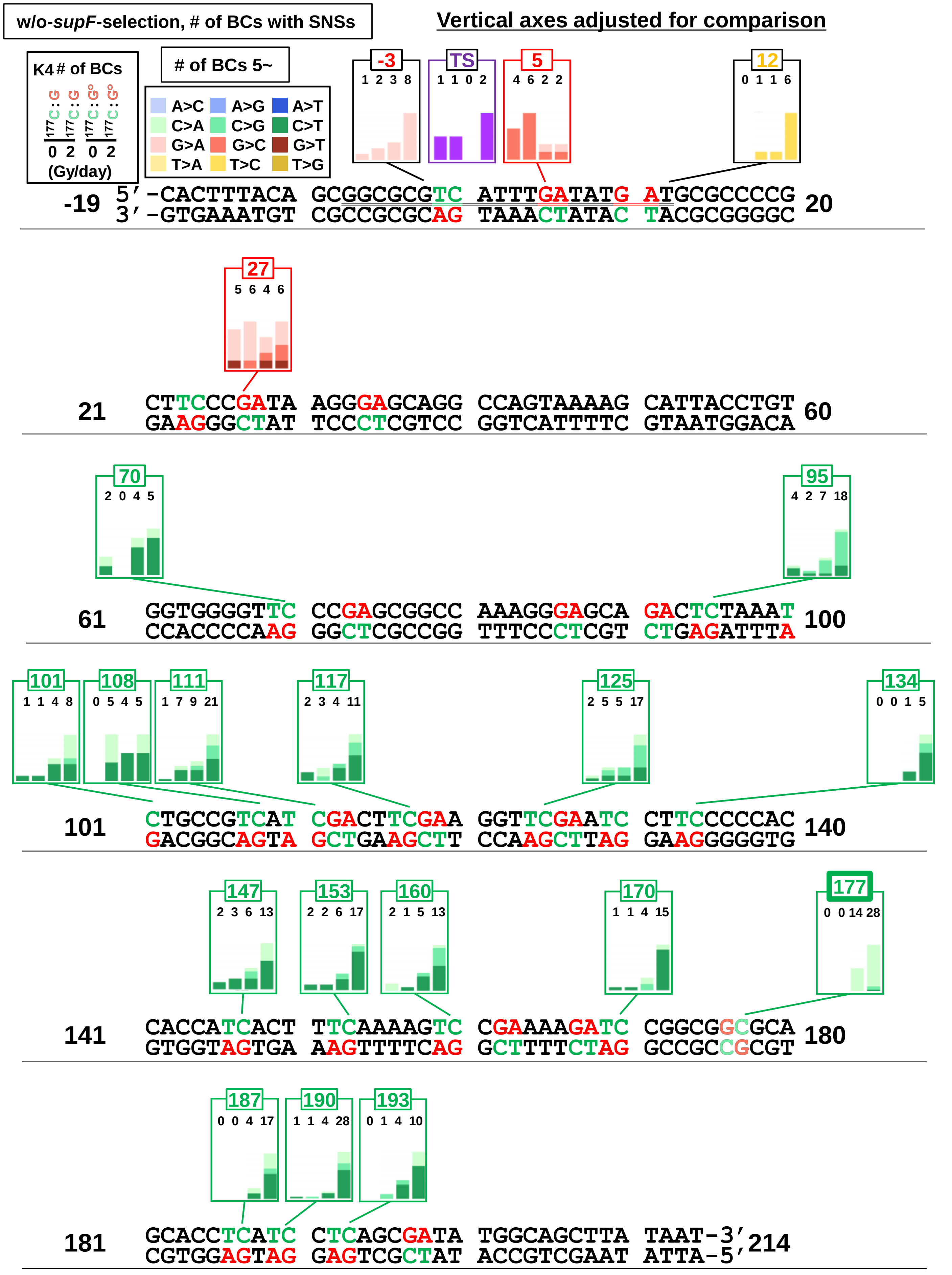


**Fig. 14-figure supplement 5.** Effects of 8-oxo-G insertion and chronic gamma-irradiation at 2 Gy/day on the proportion of substituted bases in SNSs according to their positions in the sequence (pNGS2-K4, w/o-*supF*-selection). Bar graphs provide the number of SNSs combined from three transfection experiments (TF1, TF2, and TF3). The positions and reference bases for individual SNSs are shown in different colors as indicated on the top-left side of the figure. 5'-TC-3' sites and 5'-GA-3' sites are shown in green and red respectively. Vertical axes are adjusted for comparison.
