## Supplementary material for "Systematic mutagenesis assay promotes comprehension of the strand-bias laws for mutations induced by oxidative DNA damage": Figure 14-figure supplement 6

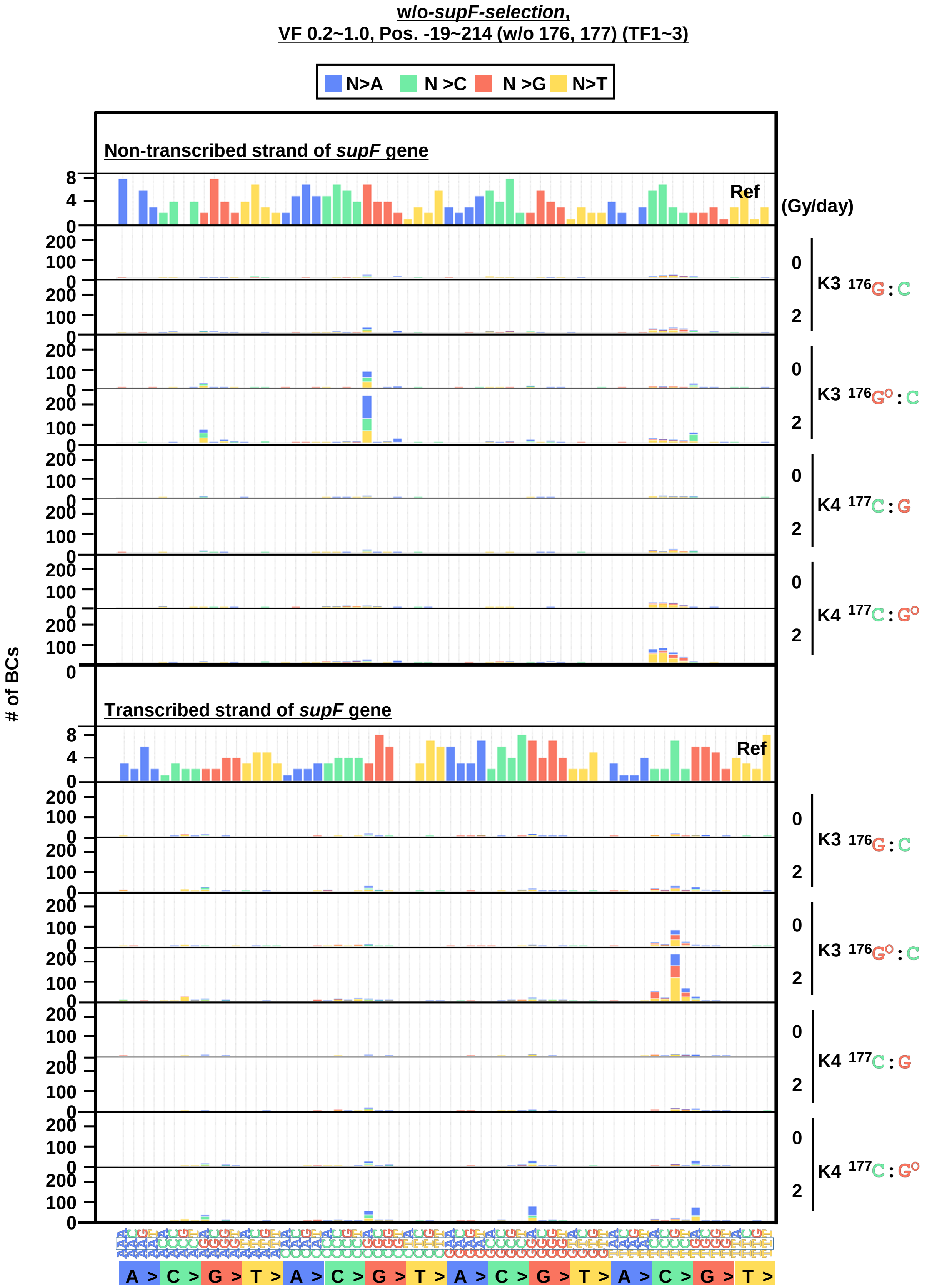


**Fig. 14-figure supplement 6.** The 192 trinucleotide contexts of SNSs for the pNGS2-K3 and K4 inserted with either a non-damaged G or an 8-oxo-G, and dose rates of chronic gamma-irradiation 0 or 2 Gy/day for 2 days (w/o-*supF*-selection, VF 0.2~1.0) (refer to the legend in Fig. 3-figure supplement 2).
