## Supplementary material for "Systematic mutagenesis assay promotes comprehension of the strand-bias laws for mutations induced by oxidative DNA damage": Figure 14-figure supplement 7

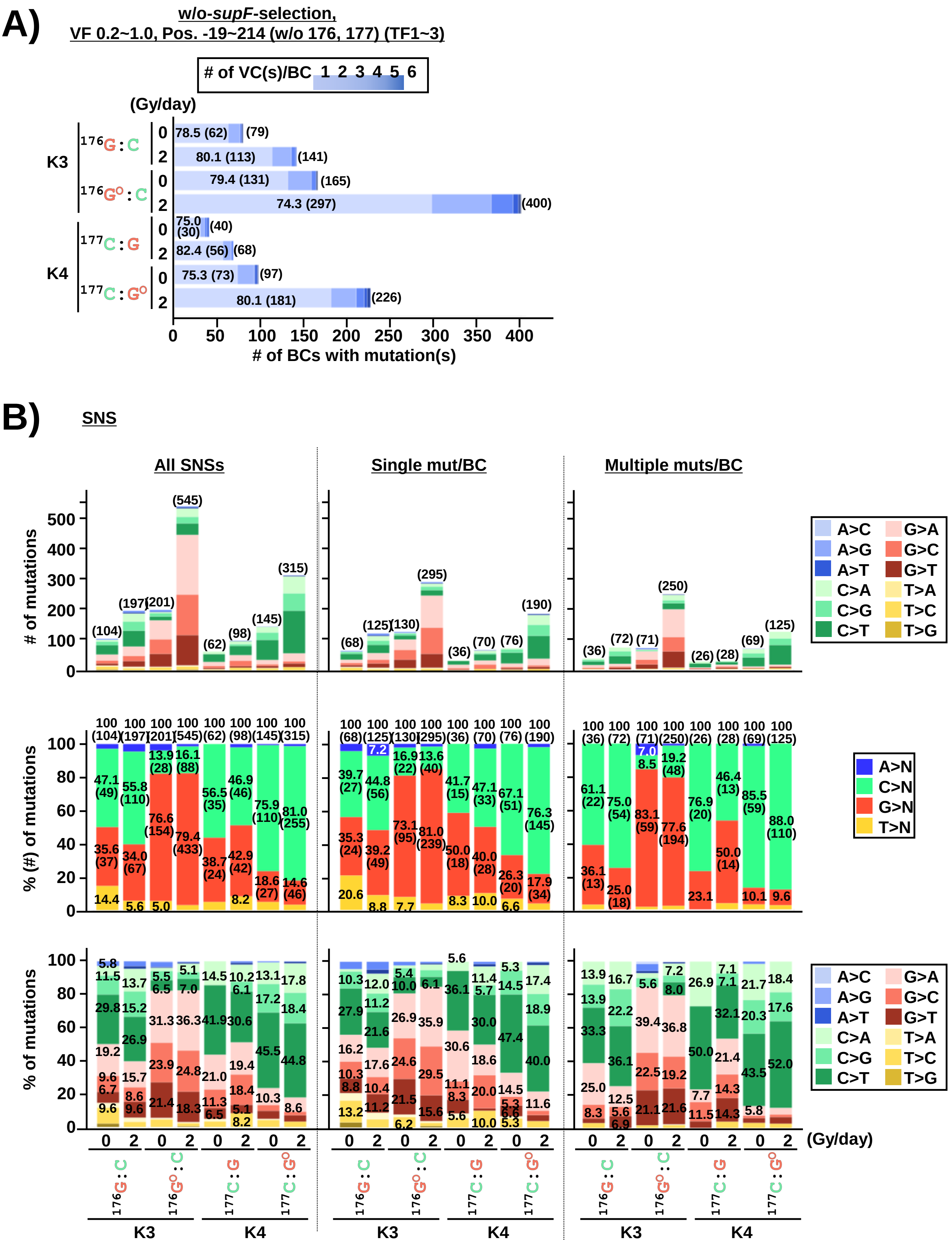


**Fig. 14-figure supplement 7.** Comparison of base substitutions by number of *supF* mutations per $N_{12}$-BC induced by the insertion of an 8-oxo-G and irradiation (w/o-*supF*-selection, VF 0.2~1.0). The layout of the figure is analogous to Figure 10. (A) The proportion of $N_{12}$-BC sequences with single (1) or multiple (2~6) mutations for pNGS2-K3 or K4 inserted with either a G or an 8-oxo-G and subjected to chronic gamma-irradiation (0 or 2 Gy/ day for 2 days). The combined data from three independent transfection experiments (TF1, TF2, and TF3) are presented. The percentage (the number) of $N_{12}$-BCs with single mutation per $N_{12}$-BC is indicated inside each bar, and the total number of mutations is presented outside each bar in the parentheses. (B) The number in each segment indicates the percentage of SNSs, and the numbers in parentheses indicate the number of SNSs.
