## Supplementary material for "Systematic mutagenesis assay promotes comprehension of the strand-bias laws for mutations induced by oxidative DNA damage": Figure 14-figure supplement 8

<u>*w/o-supF*-selection,</u>
<u>VF 0.2~1.0, Pos. -19~214 (TF1~3)</u>

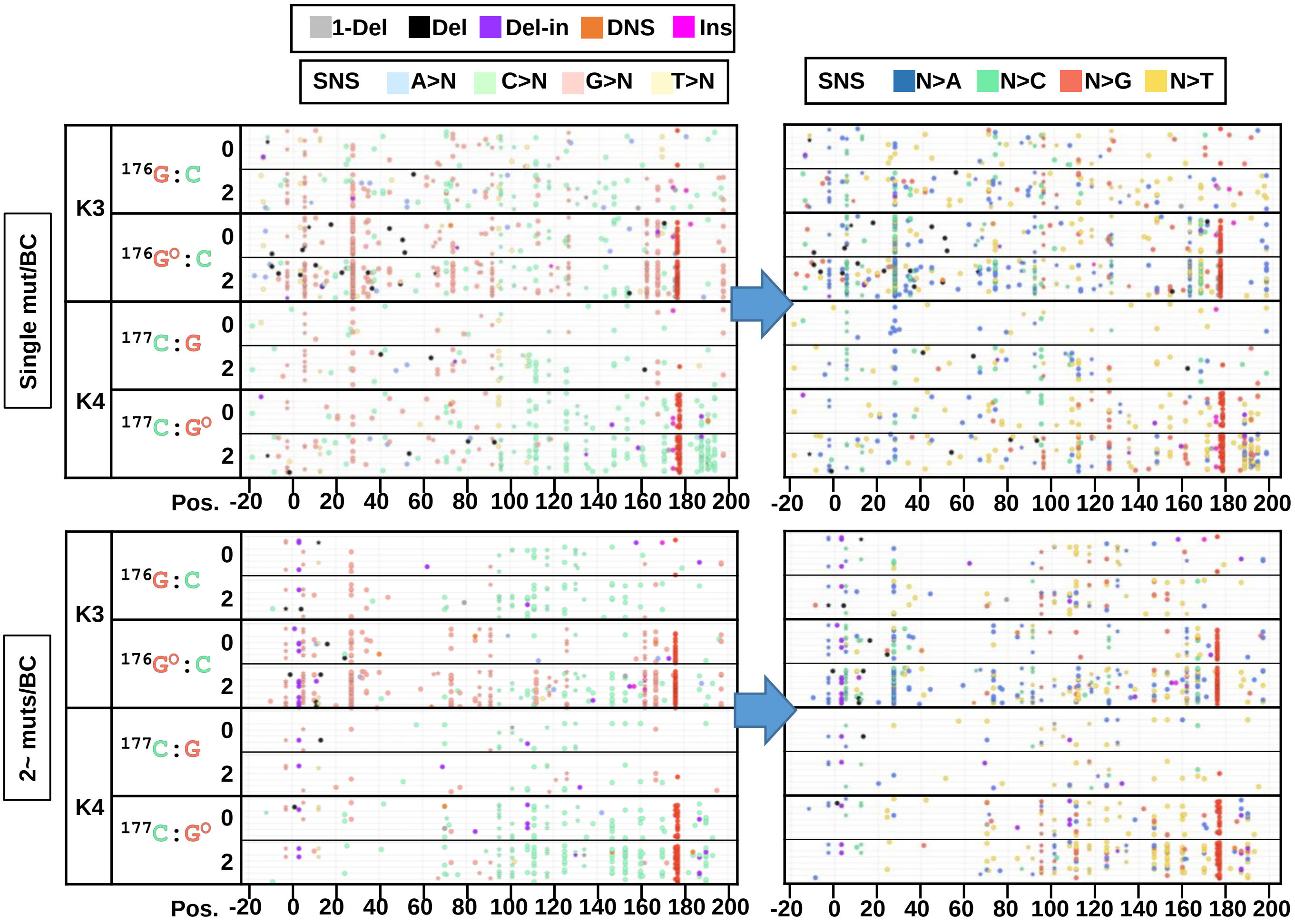


**Fig. 14-figure supplement 8.** The position and mutation type of each identified mutation (w/o-*supF*-selection, VF 0.2~1.0). The position and mutation type of each identified mutation are presented in dot plots for the cases of either single or more than one mutations (2~ muts) per $N_{12}$-BC. Each type of mutation and reference bases for SNSs are indicated by a different color in the left-side plots (legend on top). The bases after SNSs are indicated by a different color for only SNSs in the right-side plots (legend on top). The mutations in identical BCs are plotted on the same horizontal line. Data is combined from TF1, TF2, and TF3.
