## Supplementary material for "Systematic mutagenesis assay promotes comprehension of the strand-bias laws for mutations induced by oxidative DNA damage": Figure 15-figure supplement 1

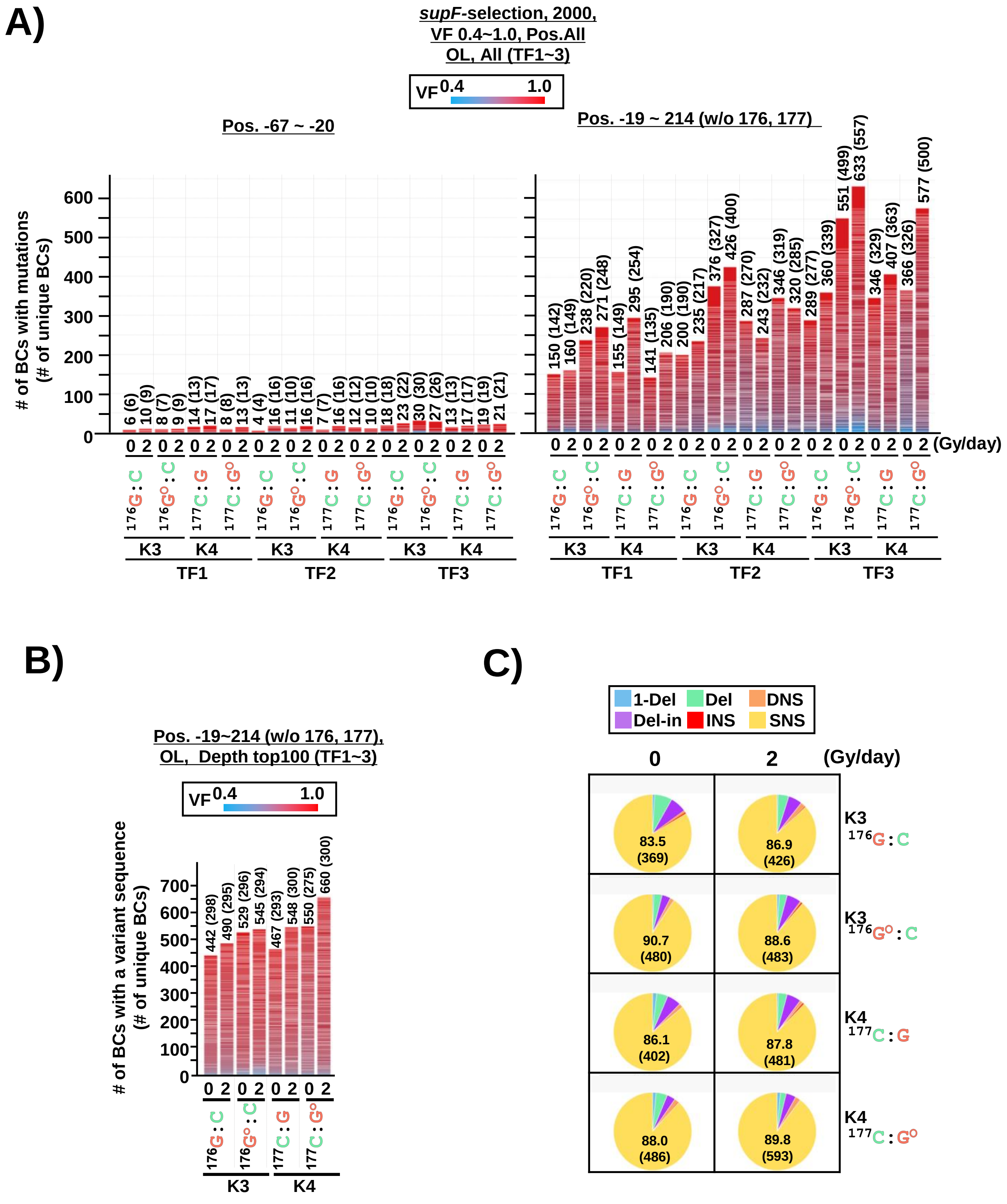


**Fig. 15-figure supplement 1.** The *supF* NGS assay for mutations in pNGS2-K3 and -K4 induced by the insertion of an 8-oxo-G and chronic gamma-irradiation (*supF*-selection, VF 0.4~1.0). (A) Stack bar graph representing the number of $N_{12}$-BC sequences with mutations obtained from each sample used for data analysis. The data are separated into two graphs: the graph on the left side represents positions from -67 to -20, and the graph on the right side – positions from -19 to 214 containing the *supF* gene. (B) Stack bar graph representing the number of $N_{12}$-BC sequences with mutations obtained from each sample used for data analysis. The one hundred $N_{12}$-BCs with at least one mutation or the maximum number of available $N_{12}$-BCs with mutations extracted from the group with the highest number of reads per unique $N_{12}$-BC. The number in parentheses refers to the number of unique $N_{12}$-BC sequences with mutations. (C) Pie charts of the proportions of mutation types in different colors as indicated at the top of panel. The percentage (the number) of SNSs are provided in each pie chart.
